# Fluorescence Blinking Patterns Fingerprint the Local Protein Environment

**DOI:** 10.64898/2026.06.22.733774

**Authors:** Salome Püntener, Dorothea Kossmann, Krzysztof Bielec, Pablo Rivera-Fuentes

## Abstract

The function of a protein depends not only on its sequence but on post-translational modifications and folding that produce functionally distinct proteoforms. Single-molecule methods for protein identification, such as nanopore sequencing, typically require denaturation or proteolysis, sacrificing conformational information that contributes to proteoform diversity. Here, we identify intact, folded proteins by recording an optical fingerprint of their local surface chemistry using a single covalent label. The signal is produced by a spontaneously blinking fluorophore attached to the protein through established bioconjugation reactions. The thermodynamics and kinetics of its switching between a fluorescent and a dark state are influenced by the immediate protein environment in a chemically interpretable manner. Further discriminative information can be extracted using deep learning to achieve excellent identification accuracy. Using this approach, we distinguish different proteins, different pockets of the same protein, and the presence of a single post-translational modification, in each case tracing the classification back to a distinct physicochemical mechanism. These results establish single-molecule fluorescence blinking as both a protein fingerprinting method and a probe of local chemistry on the surface of folded proteins.

## Introduction

The function of a protein is only partially defined by the sequence of the gene that encodes it. Splice variants, proteolytic processing, and post-translational modifications (PTMs) can produce numerous functionally distinct proteoforms from a single gene^1^. Proteoform diversity is the molecular basis of most biological regulation and disease^2,3^. Consequently, direct analysis of proteoforms is essential for a complete understanding of biological processes. The challenge is that proteoform diversity can present itself as subtle chemical modifications that are undetectable by most existing methods.

Current approaches to proteoform analysis face a fundamental challenge: methods sensitive enough to resolve subtle chemical differences typically require proteolytic digestion, destroying conformational information encoded in the intact protein. State-of-the-art protein analysis uses a bottom-up approach in which proteins are digested and the resulting peptides analyzed, either in bulk using mass spectrometry (MS)^4–6^ or at the single-molecule level using recently developed methods^7–14^, both of which sacrifice proteoform information accessible only in the full-length, folded protein. Intact proteins can be profiled using high-resolution native mass spectrometry to resolve multiple proteoforms^15–17^. Despite its advantages, native MS is challenging, may not be suitable for determining the specific location of chemical modifications, and has limited ability to distinguish isobaric species. Single-molecule methods based on nanopore readouts have also been developed to profile full-length proteins, but they still require protein denaturation with loss of native structure and conformational information^18,19^. In contrast, optical readouts such as Förster resonance energy transfer (FRET) have been used for decades to study the single-molecule dynamics and conformation^20^, and more recently the identity^21^, of proteoforms close to their fully folded, native state. However, FRET requires two labeling sites, reports on nanoscale distances rather than local surface chemistry, and is generally unsuitable to study subtle surface modifications that produce only small conformational changes.

We sought to develop a method that simultaneously achieves single-molecule sensitivity, works on intact folded proteins, and can detect subtle local surface chemistry with a single label. Towards this goal, we showed that spontaneously blinking fluorophores, which were originally developed for super-resolution microscopy^22–25^, encode information about the environment in their blinking kinetics^26,27^. We leveraged this behavior in a peptide fingerprinting method termed “blinkognition” and demonstrated that it can distinguish between synthetic peptides of different sequences, presence, and location of PTMs (Figure 1A)^27^. However, short peptides are usually conformationally flexible, and it remained unclear whether the constrained, well-defined microenvironment of a folded protein surface would enhance or ablate the information in blinking patterns.

**Figure 1.**
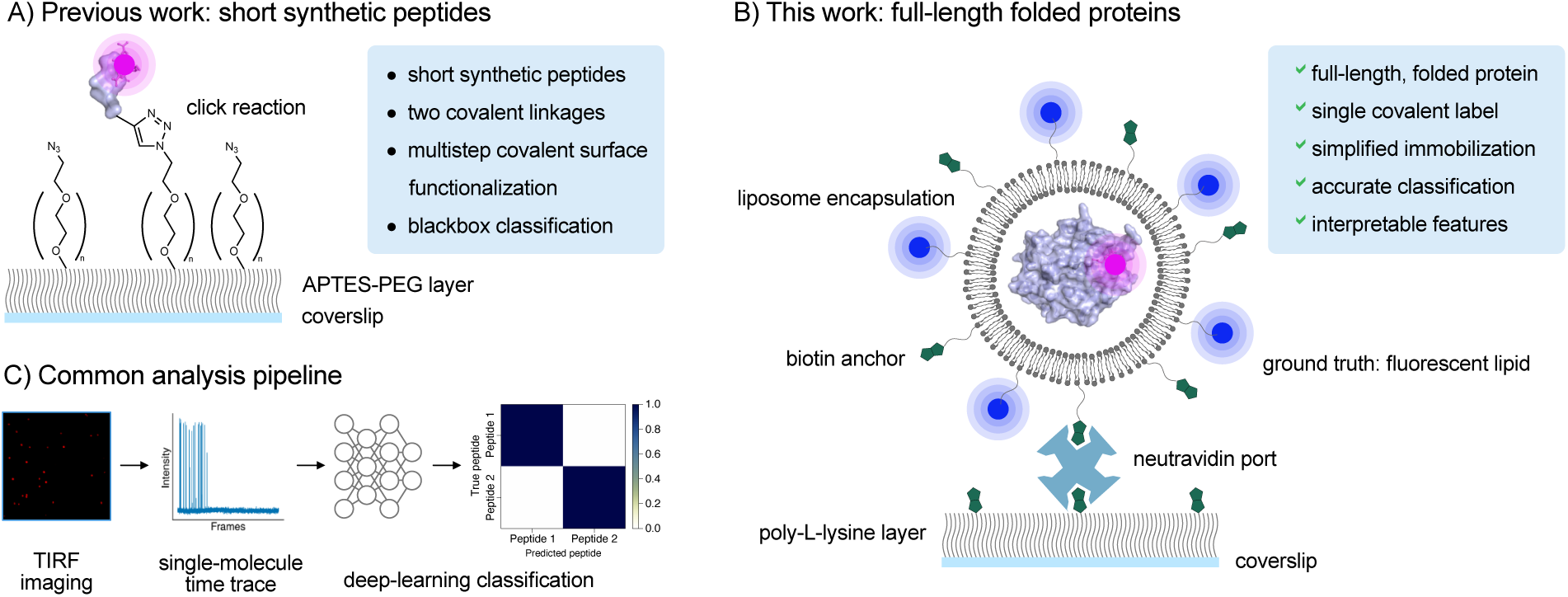
Blinkognition method for synthetic peptides (previous work) and full-length, folded proteins (this work). A) Synthetic peptides were prepared with a cysteine residue pre-labeled with a fluorophore and a modified N-terminus for surface immobilization through click chemistry. B) Proteins were only bioconjugated to the dye and non-covalently immobilized by liposome encapsulation, followed by immobilization of the vesicle to the surface via biotin-neutravidin interactions. C) General analysis pipeline for either peptide or protein single-molecule identification using fluorescent blinking patterns. APTES: (3-aminopropyl)triethoxysilane; PEG: polyethylene glycol; TIRF: total-internal reflection fluorescence.

Herein, we present protein blinkognition, a method for the accurate identification of proteins, including full-length, folded proteins and PTMs using a single covalent label (Figure 1B). In addition to accurately differentiating between two proteins, we show that protein blinkognition provides rich information about the local chemical environment of selected regions of the protein surface, extending this method from fingerprinting to single-molecule protein biophysics.

## Results and Discussion

### Liposome encapsulation enables non-covalent immobilization of proteins for single-molecule blinking analysis

In our previous work, we employed short peptides prepared by solid-phase synthesis^27^. These model systems facilitated the site-specific conjugation of the spontaneously blinking fluorophore hydroxymethyl silicon rhodamine (HMSiR)^22^ to cysteine and anchoring of the peptide to the surface via copper-catalyzed click chemistry using an alkyne at the C-terminus of the peptide (Figure 1A)^27^. To test the blinkognition concept in full-length, folded proteins, we sought an alternative immobilization strategy that required no unnatural modifications of the protein, no covalent chemistry, and minimized immobilization artifacts (Figure 1B). Our solution was to encapsulate single protein molecules in liposomes^28^, inspired by similar work in single-molecule imaging of nucleic acids^29^ and proteins^30^. Liposomes of defined sizes can be formed using a variety of lipids and are tolerant of lipid modifications^30^. This flexibility allows for surface immobilization using biotinylated lipids that can be non-covalently bound to streptavidin-functionalized surfaces^30^. Furthermore, by including a small fraction of fluorescently labeled lipids, liposomes can be imaged by flow-based methods^31,32^ and microscopy^30^, offering many opportunities for characterization.

Liposomes were generated by resuspension of 1-stearoyl-2-myristoyl-sn-glycero-3-phosphocholine in a buffered solution containing the cargo. Resuspension alone generates liposomes of varying sizes (Figure 2A), but extrusion through a 100 nm membrane homogenized the sizes of the vesicles to 120 ± 1.5 nm according to dynamic light scattering (DLS) measurements (Figure 2A). Further purification of the vesicles by size-exclusion chromatography (SEC) did not affect the size distribution of the vesicles (Figure 2A), allowing for a clean separation of vesicles and proteins that were not encapsulated and remained in solution. The liposome size distribution remained unchanged over 24 h (Figure S1), suggesting that vesicles do not undergo spontaneous fusion or fission. Furthermore, using the self-labeling protein HaloTag^33^ labeled with a fluorogenic silicon rhodamine dye^34^ and vesicles labelled with 2-dipalmitoyl-sn-glycero-3-phosphoethanolamine (DPPE) modified with dyes Atto520 (λ_ex_ = 488 nm) or Atto425 (λ_ex_ = 405 nm), we demonstrated that there is only marginal exchange of either lipids or cargo between liposomes (Figures S2 and S3).

**Figure 2.**
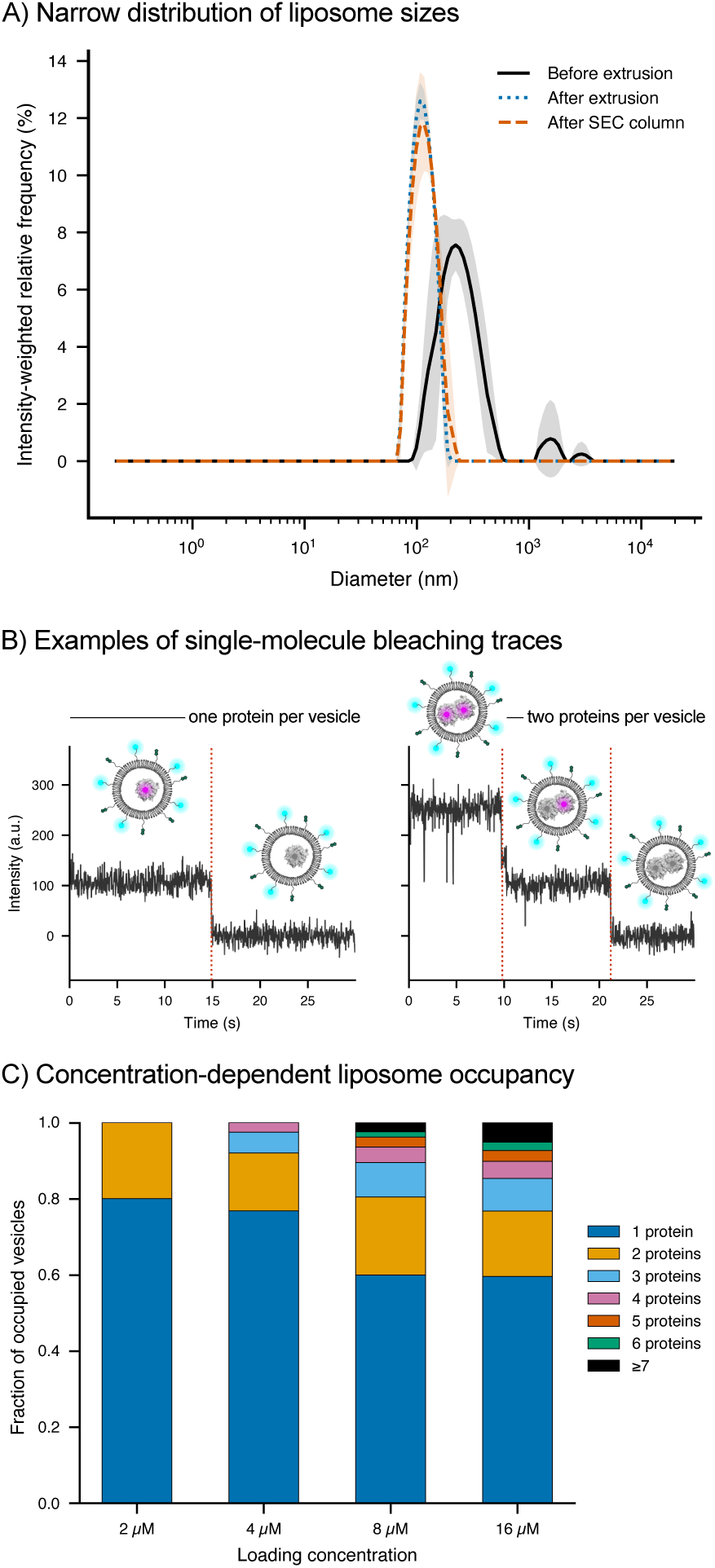
Characterization of liposomes for protein encapsulation. A) Distribution of liposome sizes measured by DLS. Solid lines are means and shaded areas represent standard deviation of three preparative replicates. B) Baseline-corrected photobleaching traces obtained from continuous imaging of liposomes with single or double occupation. Every photobleaching step (vertical dotted lines) corresponds to a single molecule. C) Number of proteins per vesicle in occupied liposomes as a function of concentration. Occupation was determined by counting photobleaching steps. Bars represent fractions for a single preparative replicate (for a second replicate see Figure S4).

We observed that many DPPE-Atto520 liposomes did not contain any protein (Figure S3). We set out to determine how many proteins exist per vesicle at different loading concentrations in the range 2-16 μM. To determine the occupancy of liposomes loaded with proteins we imaged the proteins in DPPE-Atto520 vesicles and counted the number of photobleaching events, each of which corresponds to a single molecule (Figure 2B)^35^. We found that, at all concentrations, most vesicles are empty (Figure S4) but the number of proteins per vesicle in occupied liposomes increased with loading concentration (Figure 2C and S4). At low concentrations (2-4 μM), however, a substantial majority of all occupied vesicles contain only a single protein. Moreover, these experiments also revealed that liposomes do not drift on the glass surface during the experiment, allowing for sustained imaging of the fluorescently labeled protein until the dye has photobleached. Overall, these experiments confirmed that liposome encapsulation is a simple and robust strategy to immobilize protein for single molecule imaging, with no further modification of the protein other than a single bioconjugation reaction with a functionalized blinking dye.

### Blinkognition can distinguish between two different proteins with high accuracy

Having validated the method for immobilization of single molecules of full-length, folded, and minimally modified proteins, we set out to demonstrate that blinking patterns of the spontaneously blinking dye HMSiR could be used to distinguish between two proteins. We first labeled two self-labeling proteins, HaloTag and SNAP-tag, with HMSiR functionalized with either a chloroalkane ligand (HMSiR-CA, Figure 3A) or an iodoacetamide group that reacts with reactive cysteine residues (HMSiR-IA, Figure 3A). HaloTag and SNAP-tag provide a stringent test for blinkognition because their surfaces present a similar mixture of hydrophobic and hydrophilic surface amino acids despite their unrelated sequences, and both proteins allow for site-selective labeling at defined residues (Figure 3B). Labeling of HaloTag with a chloroalkane-functionalized rhodamine dye is well established to be selective for residue D106 and quantitative (Figure 3C)^33,36^. SNAP-tag is usually labeled with a benzylguanine substrate^37^, but using HMSiR-IA led to efficient (Figure S5) and similarly selective labeling of residue C148 (SNAP-C148, Figure 3D and Figure S6). Each HMSiR-labelled protein was encapsulated in DPPE-Atto520-labeled liposomes and deposited onto a coverslip as described in the previous section. Single-molecule fluorescence data were acquired from at least two independent experiments carried out on different days and collecting data from multiple fields of view (Supporting Information).

**Figure 3.**
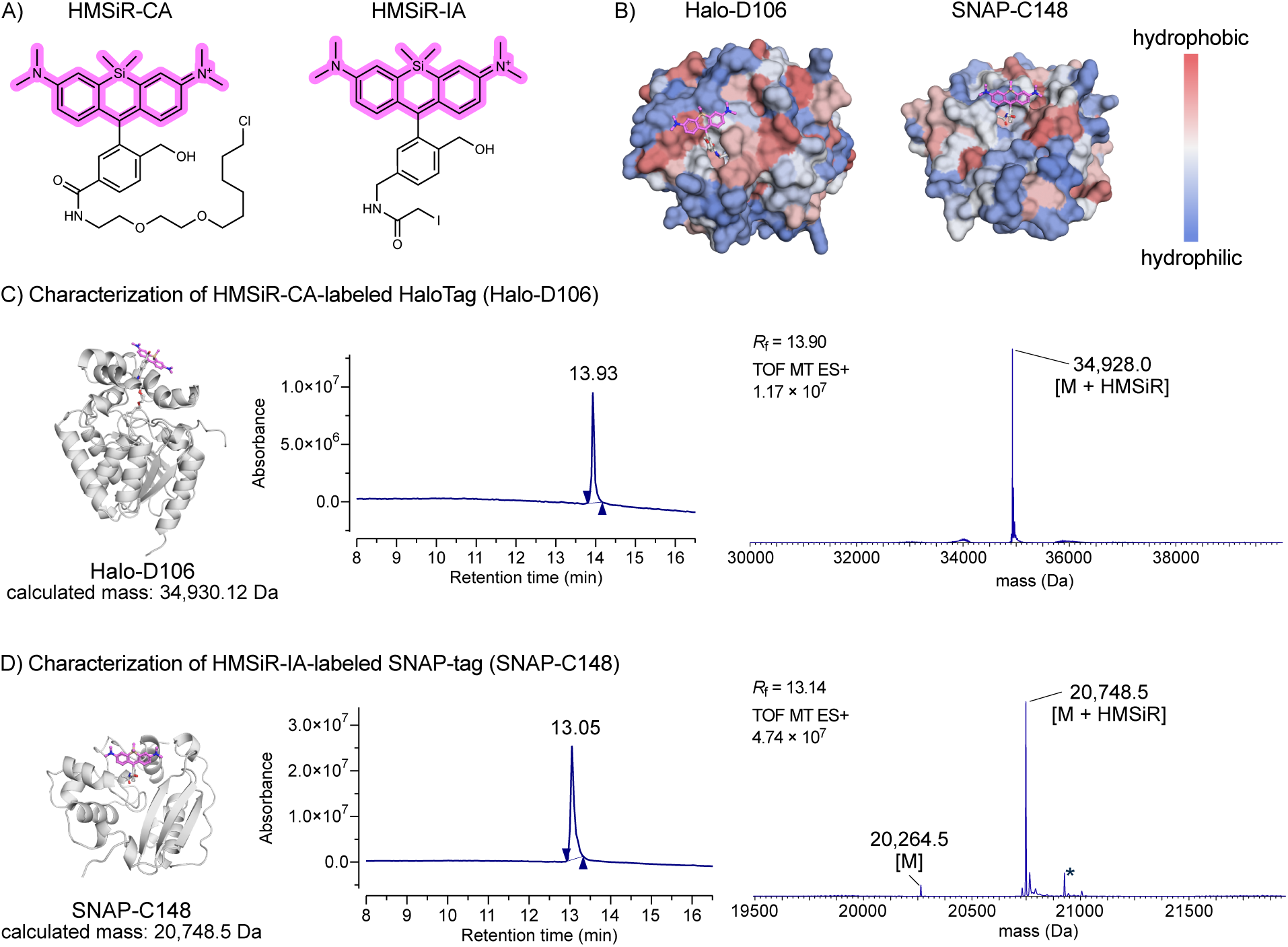
Structures and characterization of HaloTag-D106 and SNAP-C148. A) Chemical structures of HMSiR-CA and HMSiR-IA. B) Kyte-Doolittle^38^ hydrophobicity maps of HaloTag-D106 and SNAP-C148 surfaces. Models were obtained by covalent co-folding using Boltz-2^39^. C) Liquid chromatography and high-resolution mass spectrometry (LC-HRMS) analysis of HaloTag labeled with HMSiR-CA. The deconvolved mass spectrum is displayed. D) LC-HRMS analysis of SNAP-tag labeled with HMSiR-IA. The deconvolved mass spectrum is displayed. The signal marked with a star corresponds to gluconoylation, a common modification observed in N-terminal, hexahistidine-tagged proteins expressed in *Escherichia coli*^40^.

In our previous work with synthetic peptides, we developed an analysis pipeline to extract single-molecule fluorescence time traces, filter them by number of blinking events, augment the filtered traces by time reversal, and used two normalization channels for training^27^. For classification, we employed a hybrid convolutional neural network and gated recurrent units (CNN-GRU) model that leverages Monte Carlo dropout at inference time to estimate epistemic uncertainty^41^. Here, we followed a similar logic, but refactored and adapted our original code to automatically choose the optimal localization parameters per experiment and select only those proteins that colocalize with a vesicle signal (Figure 4A and Supporting Information, Figure S7). We also reformulated our trace filtering rubric to remove noisy traces more efficiently (Supporting Information, Figure S8). We used this new pipeline to extract traces for Halo-D106 and SNAP-C148 and evaluated the performance of the original CNN-GRU model for binary classification in a four-fold, cross-validation experiment (Supporting Information). Additionally, we compared this performance against those of other deep learning models, focusing on a one-dimensional residual neural network (ResNet)^42^ and a temporal convolutional neural network (TCN)^43^. Both architectures were chosen because of their computational efficiency and their strong precedent in time series classification tasks^44,45^. Furthermore, we tested whether data augmentation (time reversal or introduction of time warping, noise, and jitter, Figure S9) or using a single normalization channel affected the performance of these models in binary classification.

**Figure 4.**
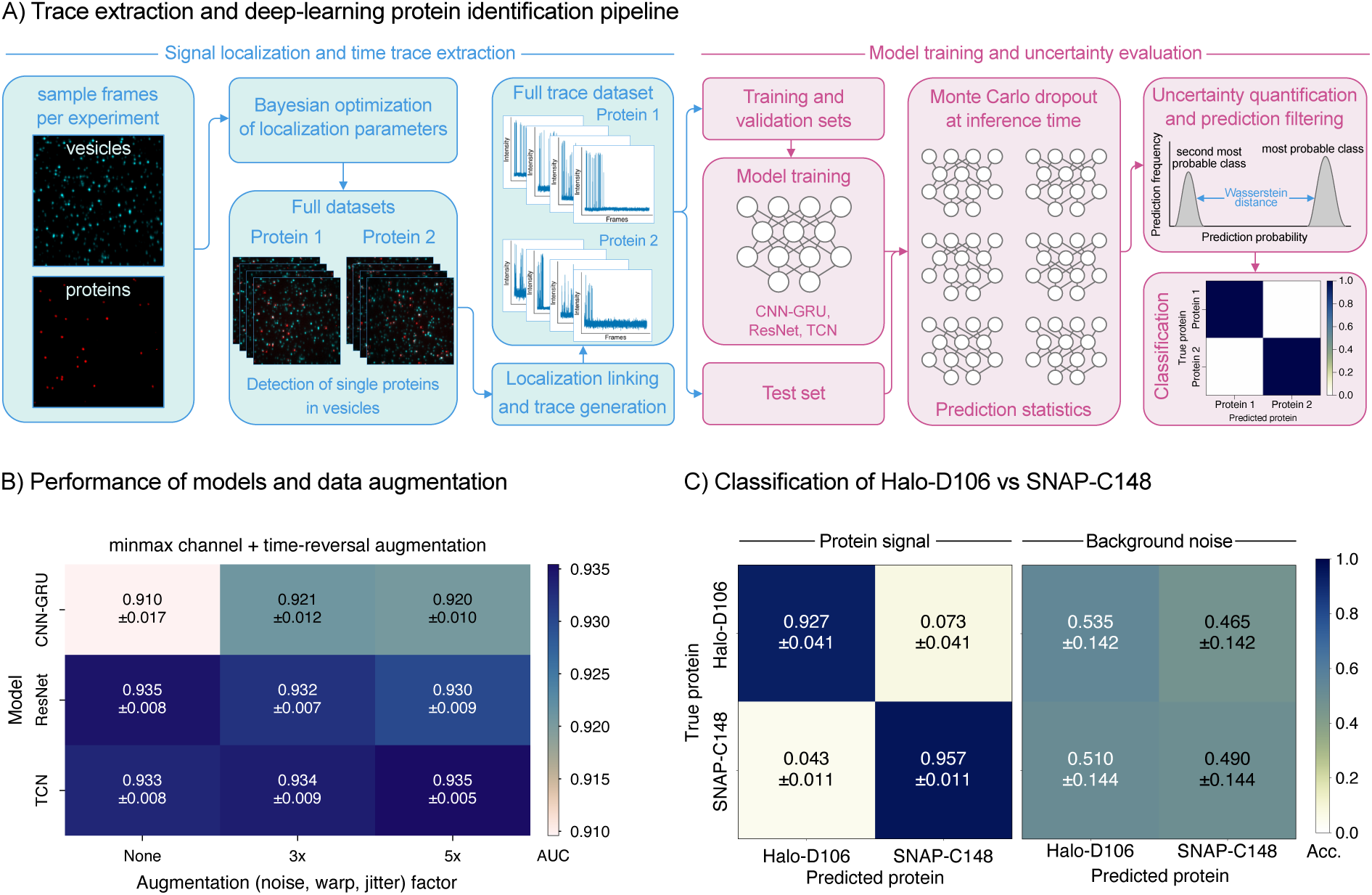
Deep-learning identification of proteins from their single-molecule blinking patterns. A) Computational pipeline from single-molecule image acquisition to assignment of protein identity. Details of each step of the analysis can be found in the Supplementary Information. B) Performance of various deep learning models and data augmentation strategies. Mean and standard deviation of four-fold cross validation are displayed in each cell. C) Confusion matrices for protein identification using filtered predictions and their corresponding background traces using the CNN-GRU model using minmax normalized traces, with time-reversal but no other further augmentation. Accuracy (Acc.) is reported as mean and standard deviation of four-fold cross validation.

All three models performed well, with values for the area under the receiver operating characteristic curve (AUC) between ∼0.910-0.935 (Figure 4B). However, closer inspection revealed important differences between these architectures in their robustness to spurious correlations. We noticed that for this dataset, augmentation and the use of two normalization channels only had a noticeable, yet modest, effect on the performance of the CNN-GRU model (Figure S10). Both the ResNet and TCN models performed slightly better than the CNN-GRU model under all augmentation and normalization conditions, and they were both trained faster than the CNN-GRU model (Figure S11). However, when we performed a noise-only training experiment (Supporting Information and Figure S12), both ResNet and TCN models seemed to learn patterns from baseline noise (Figure S13). This behavior seemed slightly worse for both the dual-channel or z-score-normalized traces (Figure S13). Therefore, despite the apparent excellent performance and short training times of the ResNet and TCN models, only the CNN-GRU model seems to learn from blinking signals without learning patterns from baseline noise for this dataset. Nevertheless, the CNN-GRU model with minmax normalized traces and uncertainty filtering gave excellent classification accuracy (Figure 3C). These results establish that blinkognition, combined with liposome encapsulation, can accurately identify full-length folded proteins at the single-molecule level using only a single covalent label.

### Blinking patterns contain chemically interpretable information of the local protein environment

A central hypothesis of blinkognition is that the blinking pattern responds to the chemical environment of the fluorophore, which in the case of proteins, varies depending on the specific amino acid that is labeled with the blinking dye. We tested this hypothesis by labeling HaloTag with HMSiR-IA, which led to the selective labeling of residue K117 (Figure 5A and Figures S14 and S15). This alternative covalent binding site provides the dye with a very different local environment and therefore should be distinguishable from the local environment in Halo-D106 (Figure 5A). We encapsulated Halo-D106 and Halo-K117 as described before, imaged them, and trained all combinations of models and augmentation strategies described in the previous section (Figure S16). Once again, we found that the CNN-GRU model using minmax-normalized traces gave the highest accuracy without learning patterns from baseline noise (Figure 5B).

**Figure 5.**
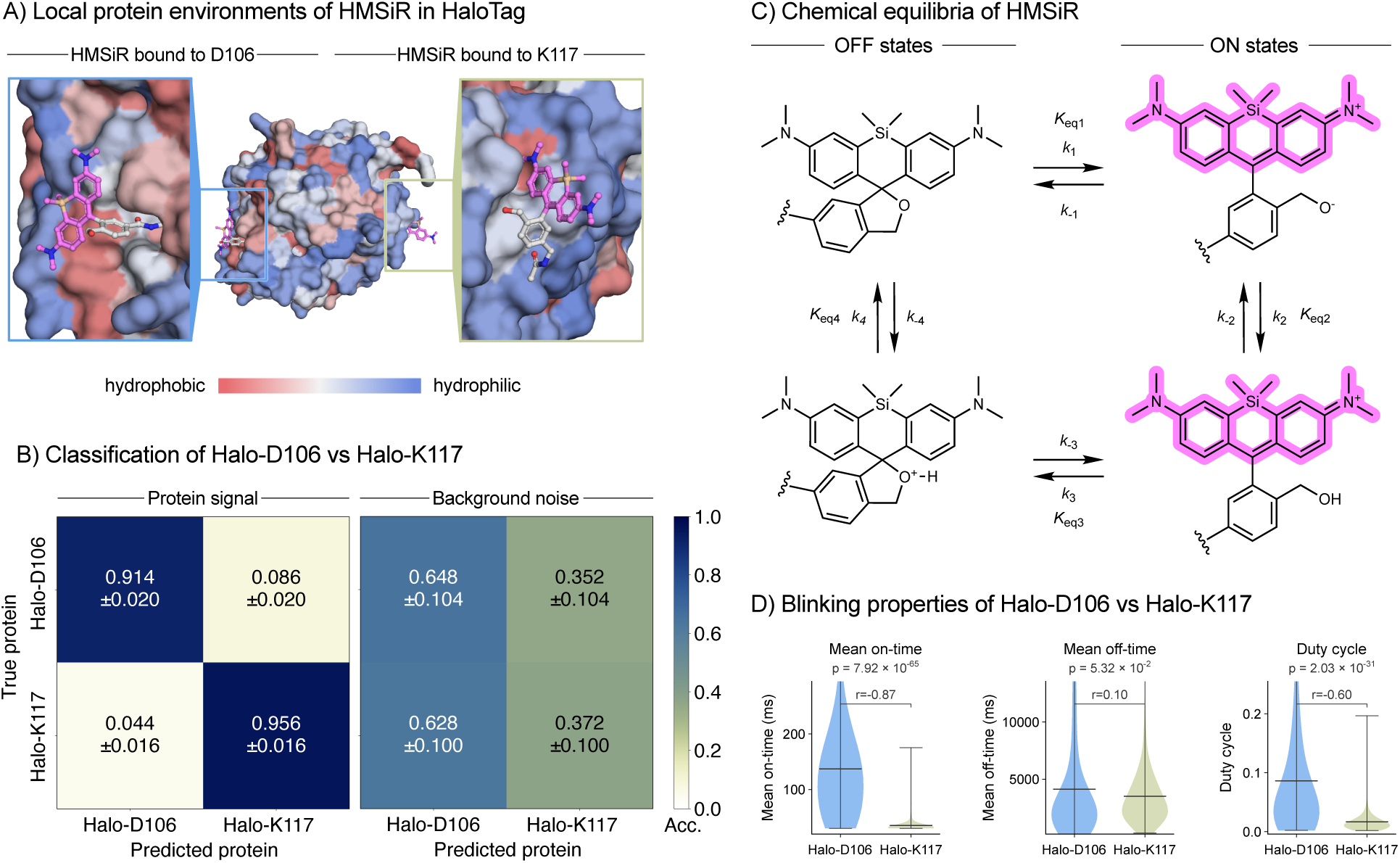
Blinkognition discriminates between two different protein local environments. A) Overlay of two Boltz-2 models of the protonated, ON state of HMSiR bound to HaloTag at either D106 or K117. Protein surface colored according to the Kyte-Doolittle^38^ hydrophobicity scale. Alternative poses found by Boltz-2 are displayed in Figures S17 and S18. B) Confusion matrices for protein identification using filtered predictions and their corresponding background traces using the CNN-GRU model without time-reversal or any further augmentation. Accuracy (Acc.) is reported as mean and standard deviation of four-fold cross validation. C) Chemical structures of the OFF and ON states of HMSiR and equilibrium and rate constants for the main transformations. D) Chemically interpretable blinking properties of HMSiR bound to HaloTag at residues D106 or K117. Mean off- and on-times, and duty cycle were calculated per single-molecule trace (Halo-D106: n=262, Halo-K117: n=248 single molecules pooled from two preparative replicates). The violin plots aggregate single-molecule features and mean and standard deviation are shown. Significance brackets show pairwise comparisons between protein classes for each feature (Mann-Whitney U test, two-sided; p-values corrected for three simultaneous comparisons using the Benjamini-Hochberg false discovery rate procedure). Effect size is reported as the rank-biserial correlation r.

The ability of blinkognition to distinguish between two different subregions of the same protein indicates that there must be local chemical properties of the protein that lead to different blinking features. Recent work with HMSiR and synthetic peptides suggested that chemically interpretable properties such as hydrophobicity or H-bond partners may be mapped to blinking features^46^. The blinking pattern of HMSiR is determined by: 1) the relative populations of the ring-open fluorescent (ON) form, which itself is an equilibrium of a protonated cationic and a deprotonated zwitterionic state, and the ring-closed dark (OFF) form, which may also exist as the neutral ether or protonated form (Figure 5C); and 2) the kinetics of interconversion between all four states. These thermodynamic and kinetic parameters are affected by the environment provided by the protein.

We inspected the mean ON times, mean OFF times, and the duty cycle (total ON time over total ON + OFF time) for every single molecule of Halo-D106 and Halo-K117 that was correctly and confidently classified by the CNN-GRU model (Figure 5D). Each of these blinking properties maps to distinct processes in the blinking equilibrium (Figure 5C). The mean ON time reflects the ring-closing process, which is governed by *k*_-1_, *k*_3_ and the protonation equilibrium *K*_eq2_ = *k*_2_/*k*_-2_. Similarly, mean OFF time reflects the ring-opening process, which is governed by rate constants *k*_1_ and *k*_-3_, and the protonation equilibrium *K*_eq4_ = *k*_4_/*k_-_*_4_. If a single molecule samples enough ON and OFF states before photobleaching (ergodic assumption), the duty cycle is related to the thermodynamic equilibrium between the ON and OFF species *K*_eq,ON/OFF_ = *K*_eq1_/*K*_eq2_. The combination of shorter ON times (Halo-D106: 137 ± 139 ms vs Halo-K117: 35 ± 15 ms, Figure 5D), lower duty cycle (Halo-D106: 0.08 ± 0.1 vs Halo-K117: 0.017 ± 0.02, Figure 5D), and mostly unchanged OFF times in Halo-K117 (Halo-D106: 4135 ± 4891 ms vs Halo-K117: 3509 ± 2897 ms, Figure 5D) indicates that the equilibrium is shifted toward the OFF states in Halo-K117 compared to Halo-D106.

Since the ON states are charged (Figure 5C), one would expect destabilization in hydrophobic environments. Inspection of covalent co-folding models obtained using Boltz-2 (Figure 5A)^39^ suggests that the protein surface around HMSiR in Halo-D106 is more hydrophobic than the environment in Halo-K117. At first glance this appears contradictory, since the charged, ON states should be destabilized in hydrophobic environments. However, the vast majority of Boltz-2-predicted structures clustered around a pose in which only one face of the HMSiR dye in D106 is exposed to the hydrophobic surface, and the opposite side, which bears the critical hydroxymethyl group is completely exposed to the solvent, where it could be solvated and eventually protonated by water molecules (Figure 5A and Figure S17). The mean ON time in Halo-D106 of 137 ± 139 ms sits in between the value measured for the free dye in solution by flash photolysis (245 ms) and that measured for the dye attached to an antibody (96 ms)^22^, and is therefore consistent with the Boltz-2 model that shows minimal interaction of the hydroxymethyl substituent with the protein.

In contrast, all the main poses predicted for HMSiR in Halo-K117, where the local protein environment is more polar, place both sides of the dye in contact with the protein, with the hydroxymethyl substituent substantially less exposed to the solvent (Figure 5A and Figure S18). Since the zwitterionic form, in which the hydroxymethyl is deprotonated, is the most reactive species for ring closure, any interaction that stabilizes this form or facilitates its generation would accelerate ring closure. This effect may arise from protein residues near the hydroxymethyl group that act as general bases facilitating deprotonation (increasing *k*_-2_) or slow down protonation by the solvent (decreasing *k*_2_) by stabilization of the alkoxide. If ring closure (*k*_-1_) is fast, both effects would shorten the mean ON time and shift the equilibrium toward the OFF state without drastic effects on the mean OFF time (unaffected *k*_1_), which is consistent with our observations. Together, these results demonstrate that blinking patterns encode mechanistically interpretable information about the chemical environment of individual residues on a folded protein surface, establishing blinkognition as both a fingerprinting method and a single-molecule probe of local protein chemistry.

### Blinkognition can detect post-translational modifications with single-molecule sensitivity

The accurate detection of post-translational modifications in proteins is challenging and single-molecule methods for this task are only starting to emerge^27,47–51,21,52,53^. Given the excellent sensitivity of blinking patterns to the chemical environment on the protein surface, we tested whether the blinking fingerprint was sufficiently sensitive to detect the presence of a post-translational modification on an amino acid in the vicinity of the HMSiR dye. For this experiment we employed a single-cysteine (sc) variant of the human redox protein glutaredoxin 1 (Grx1, Uniprot: P35754, Figure 6A). Grx1 is an essential protein in the modulation of reversible protein *S*-glutathionylation^54^. In humans, this protein can be acetylated at lysine residue K20^55^. The only cysteine residue in scGrx1 is C23, and we could label this residue efficiently with HMSiR-IA (Figures S19 and S20). Additionally, we expressed and purified scGrx1 with acetylated lysine K20 (scGrx1-AcK20, Figure 6A and Figures S21 and S22), which was site-selectively introduced by genetic code expansion^56^. Labeling of C23 with HMSiR-IA also proceeded with high selectivity in the acetylated protein (Figure S22). Both labeled scGrx1 variants were encapsulated in liposomes and imaged as described before to obtain single-molecule blinking traces of both proteins.

**Figure 6.**
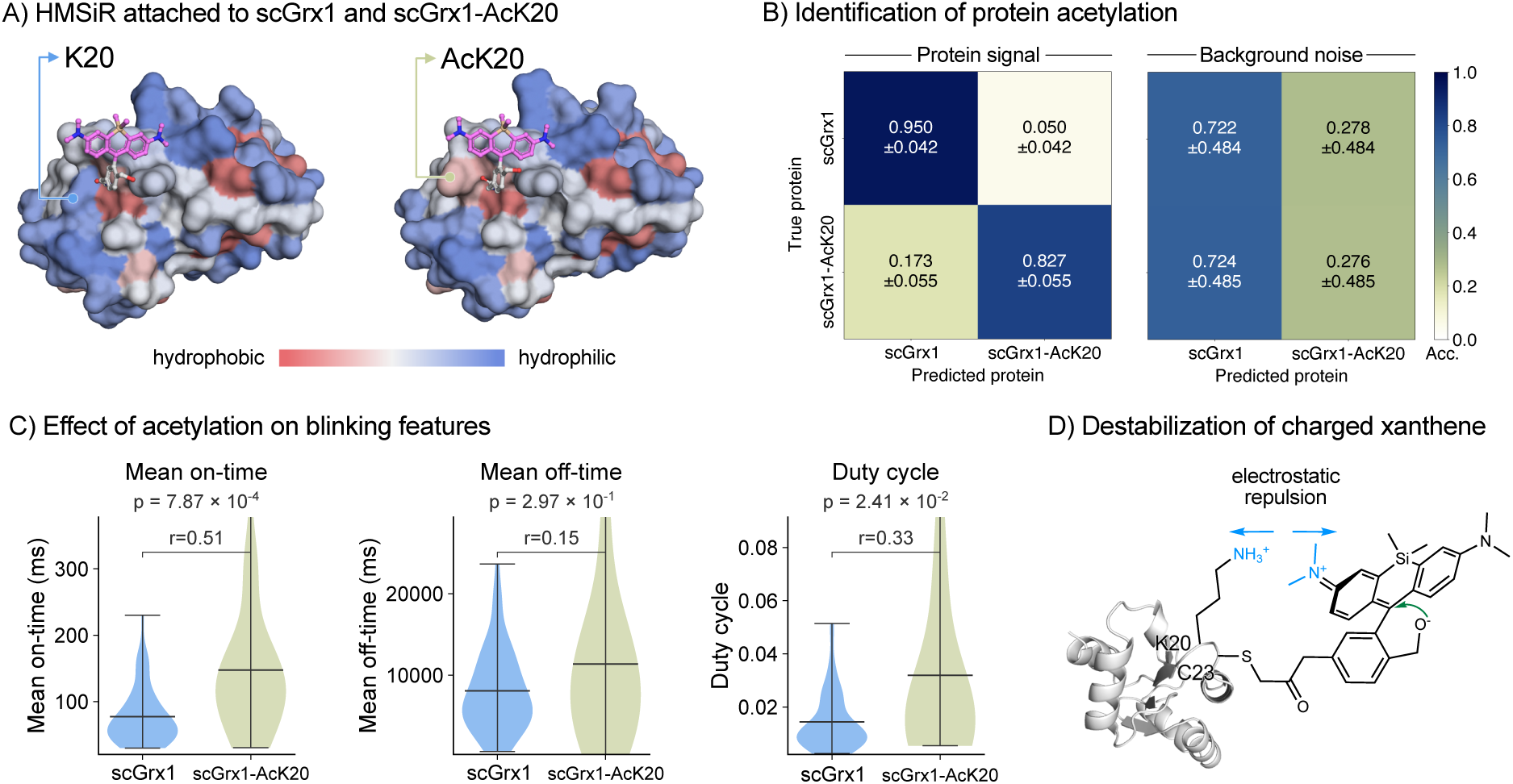
Detection of lysine acetylation. A) Boltz-2 structures of scGrx1 and its K20 acetylated version labeled at C23 with HMSiR. Protein surface colored according to the Kyte-Doolittle^38^ hydrophobicity scale. Alternative poses found by Boltz-2 are displayed in Figures S27 and S28. B) Confusion matrices for protein identification using filtered predictions and their corresponding background traces using the TCN model with time-reversal but no further augmentation. Accuracy (Acc.) is reported as mean and standard deviation of four-fold cross validation. C) Chemically interpretable blinking properties of HMSiR bound to scGrx1 or scGrx1-AcK20 at residue C23. Mean off- and on-times, and duty cycles were calculated per single-molecule trace (scGrx1: n=59, scGrx1-AcK20: n=25 single molecules pooled from four preparative replicates). The violin plots aggregate single-molecule features and mean and standard deviation are shown. Significance brackets show pairwise comparisons between protein classes for each feature (Mann-Whitney U test, two-sided; p-values corrected for four simultaneous comparisons using the Benjamini-Hochberg false discovery rate procedure). Effect size is reported as the rank-biserial correlation r. D) Proposed electrostatic destabilization of the charged, ON species in scGrx1 by K20, leading to shorter ON times and duty cycles.

Distinguishing between scGrx1 and scGrx1-AcK20 is a challenging task, and the CNN-GRU model gave only moderate AUC values across augmentation strategies (Figure S23). In contrast, we observed that the TCN model performed robustly without substantial learning from background noise (Figure 6B and Figure S23). We further confirmed that the TCN model does not learn background-related features by performing a label-shuffling classification experiment and using the model trained on protein traces to classify noise traces (Figure S24), observing in each case no learning or classification ability. Having established the suitability of the TCN model for this dataset, we used it for binary classification and the confusion matrix revealed that the model correctly identifies scGrx1 traces with high confidence, whereas about 17% of the traces of scGrx1-AcK20 are identified as scGrx1 with high certainty. This result is also reproduced by the ResNet model (Figure S25). Closer inspection of the MS data revealed that about 13% of the proteins either incorporated regular lysine or were deacetylated during expression and purification (Figure S26), suggesting that the misclassification might be at least partially a consequence of contamination of scGrx1-AcK20 traces with genuine scGrx1 traces from non-acetylated proteins.

Analysis of the blinking features revealed that scGrx1-AcK20 has substantially longer mean ON times (147 ± 103 ms) than scGrx1 (77 ± 43 ms), indicating that the fluorescent charged state is longer-lived in the acetylated protein (Figure 6C). The Boltz-2-predicted models show very similar poses of HMSiR bound to scGrx1 and scGrx1-AcK20 (Figure 6A and Figures S27 and FigureS28), with K20 behind the xanthene dye and the hydroxymethyl group on the opposite side pointing toward the solvent. Based on these models, we hypothesize that the positive charge of K20 has an unfavorable electrostatic interaction with the positive charge of the xanthene chromophore in its ON state, whereas acetylation at K20 removes this electrostatic penalty leading to longer ON times in scGrx1-AcK20. The uncharged OFF state does not seem to have significantly different lifetimes between the two proteins, leading to a smaller duty cycle in scGrx1. These observations are consistent with electrostatic destabilization of the charged species of HMSiR in the positively charged pocket of scGrx1, compared to the neutral scGrx1-AcK20.

Finally, we tested whether acetylation of scGrx1 could be detected using only the mean, standard deviation and sum of ON and OFF times per trace as features. A simple random forest classifier with uncertainty filtering (Supporting Information) could indeed leverage these six features to distinguish between scGrx1 and scGrx1-AcK20 with an accuracy of 74% (Figure S29). Importantly, the features derived from the OFF states have the highest Gini importance in the classification task, which is consistent with our analysis (Figure S29). The accuracy of the random forest is lower than that of the TCN classifier (Figure 6B), suggesting that the deep learning model finds additional information in the kinetic fine structure of blinking. Nevertheless, the random forest classifier provides a baseline model and supports the conclusion that proteoform identification using blinking fingerprints can be mapped to chemically interpretable features.

These results demonstrate that blinkognition can detect a single-residue post-translational modification at the single-molecule level, and the underlying blinking features are traceable to a specific electrostatic mechanism. Together with the Halo-D106 versus Halo-K117 comparison, this establishes that blinking patterns can distinguish between fundamentally different types of chemical perturbations. These examples demonstrate that protein blinkognition provides chemically interpretable information about the nature of the modification, not just its presence.

## Conclusions

The spontaneous blinking pattern of HMSiR is sensitive to its local environment. When it is covalently attached to a protein, the chemical environment imposed by nearby amino acids affects the blinking pattern of HMSiR in a chemically interpretable manner. However, deep learning methods learn more than only these intuitive features and lead to very accurate identification of separate proteins, different conjugation sites of the same protein, or the presence of post-translational modifications. The single-molecule nature of this method makes it suitable to study effects that might be obscured in bulk measurements and makes it potentially more sensitive to small conformational changes than existing alternatives. The fact that this method uses full-length, folded proteins makes it suitable to study conformation-specific effects that would get lost in other single-molecule protein identification methods that rely on protein unfolding, proteolytic cleavage, or multiple bioconjugations with large labels. Additionally, the requirement of only a single bioconjugation reaction greatly simplifies sample preparation and makes it widely applicable to PTMs that do not tolerate the introduction of additional labels. The lack of a direct linkage between the protein and the surface alleviates immobilization artifacts^28^. These characteristics make protein blinkognition a promising tool for single-molecule proteomics and biophysics.

We have identified a few areas of improvement that would make protein blinkognition a more generally applicable method. The most important aspect is the development of efficient and selective bioconjugation reactions for proteins that contain multiple cysteines or none. Although activated esters partially fulfill this task^57–60^, they still lack in site specificity. The development of brighter and less hydrophobic dyes could improve signal and bioconjugation yields. Additionally, dyes with an inherently longer ON times might be able to catch more environment-specific nuances that are currently obscured by the time resolution (30 ms) of our experiments. From a mechanistic point of view, molecular dynamics simulations would be valuable to understand, and potentially predict, changes in blinking features depending on the chemical nature of the protein surface. Finally, a method that could directly couple the fractionation of complex mixtures of proteins to single-molecule imaging would facilitate the application of protein blinkognition to disease-relevant samples.

## Supporting Information

Detailed description of experimental procedures and computational analysis pipelines. Supporting figures and NMR spectra.

## Data availability

Raw data to reproduce all the results of this paper are available on Zenodo (DOI: 10.5281/zenodo.20394521). The only exception is the original ND2 movies (1 TB of data), which are available from the authors upon request.

## Code availability

Python scripts to reproduce every figure in the paper are provided along with the raw data (DOI: 10.5281/zenodo.20394521). The blinkognition codebase is available on GitHub: https://github.com/locbp-uzh/blinkognition.git

## Acknowledgments

Protein MS analyses were carried out at the Functional Genomics Center Zurich by Dr. Chia-Wei Tan-Lin and Dr. Serge Chesnov. Computational work was carried out using infrastructure of the scientific computing center S3IT of University of Zurich.

## Use of AI-assisted tools

Substantial amounts of code were written or refactored by Claude Code (Sonnet 4.6 and Opus 4.6 and 4.7). Claude’s Opus 4.7 was also used to edit the manuscript for clarity.

## Funding

This work was funded by the European Research Council (Starting Grant: HDPROBES, 801572) and University of Zurich.

## Author contributions

S. P. and P. R.-F. formulated hypotheses and planned the experiments. S. P., D. K., and K. B. performed experiments. S. P. and P. R.-F. wrote code and carried out data analysis. S. P., D. K., and P. R.-F. wrote the manuscript with input from K. B. P. R.-F. supervised the project and acquired funding.

## Conflicts of Interest

S. P. and P. R.-F. are named inventors on European patent application EP 23 813 767.3 (“Biomolecule analysis by fluorescence intermittence signal recognition”), filed by the University of Zurich, which relates to the method described in this manuscript. D. K. and K. B. declare no competing interests.

## Notes

https://zenodo.org/records/20394521

https://github.com/locbp-uzh/blinkognition

