## Supplementary Material for "Fluorescence Blinking Patterns Fingerprint the Local Protein Environment"

### General Methods

#### Reagents and analyses

Lipids for encapsulation were purchased from Avanti Polar Lipids (1-Stearoyl-2-myristoyl-sn-glycero-3-phosphocholine (SMPC): 850464C, 16:0 NBD PE: 810144P 16:0 biotinyl PE: 870277P). The fluorescent lipids DPPE-Atto425 and DPPE-Atto520 were obtained from ATTO-TEC. Reagents for PEGylation were purchased from Laysan Bio. (biotin-$PEG_{5000}$-NHS: #Biotin-PEG-SVA-5000-1g) and PLL-g-PEGs from SuSoS (biotinyl PLL-g-PEG: #PLL(20)-g[3.5]-PEG(2)/PEG(3.4)biotin20%, PLL-g-PEG: # PLL(20)-g[3.5]-PEG(2)). DLS measurements were conducted on a Litesizer 500 (Anton Paar) using a refractive index of 1.334 and the absorption settings for proteins were applied. Vesicles and their fluorescence were further characterized with a flow nanoanalyzer (NanoFCM Inc.) equipped with a 488 nm and 640 nm laser, and records SSC and up to two fluorescent channels with highly sensitive photon avalanche diodes. For size calibration in the NanoAnalyzer S16M-Exo-beads were used. Stock solutions of synthesized chemical probes were prepared in DMSO spectrophotometric grade > 99.9% (Acros Organics) at a concentration of 20 mM, diluted aliquots were prepared and stored at –20 °C. The fluorogenic silicon rhodamine dye SiBr was reported previously[1]. All stock solutions were thawed immediately before use.

#### Protein Expression

BL21(DE3) competent cells (New England Biolabs) were transformed with the respective plasmid by heat shock following the provided standard procedure of the vendor. LB medium was prepared with the appropriate antibiotic. A single colony was inoculated into the LB medium and incubated at 37 °C overnight. The desired volume of LB medium was inoculated from the starter culture and incubated at 37 °C until OD=0.4–0.8 was reached. Isopropyl β-D-1-thiogalactopyranoside was added to a final concentration of 0.5–1 mM, and the culture was further incubated at 18 °C overnight. *E. coli* cells were harvested by centrifugation. The bacterial pellet was resuspended in lysis buffer (300 mM NaCl, 50 mM PB at pH = 7.5, 10% glycerol, 50 mM tris(2-carboxyethyl)phosphine) as well as Turbonuclease (5 µL per L of culture) and a protease inhibitor cocktail tablet (Roche). The cells were lysed by sonication (70% amplitude,10 s pulse-10 s pulse off for 2.5 min). The lysate was cleared by centrifugation and the protein was purified by Ni-His-affinity chromatography in batch mode on HisPur™ Ni-NTA resin (ThermoFisher Scientific) or using an ÄKTA start chromatography system (Cytiva) equipped with a 1 mL HisTrap column (Cytiva). Fractions as well as induction controls were analyzed by SDS page gel electrophoresis, and pure fractions were pooled and dialyzed against phosphate-buffered saline (PBS). The protein samples in PBS were aliquoted and stored at –70 °C.

**Expression of scGrx1-AcK20 by genetic code expansion**

Electrocompetent *E. coli* B-95.Δ*A*Δ*fabR* cells were transformed with pET28a-His6-scGrx1-K20Ac (containing the Amber stop codon UAG at position 20) and pTECH-chAcK3RS(IPYE) (Addgene, #104069)[2] by an electric pulse (2500 V, 25 µF, 200 ohm) using a Gene Pulser XCell ShockPod (Bio-Rad). Double-transformed cells were selected by cultivation on LB agar plates with kanamycin and chloramphenicol. A single colony was picked and used to inoculate a liquid LB culture containing kanamycin and chloramphenicol, which was incubated at 37 °C and 180 rpm overnight. The overnight culture was used to inoculate a fresh culture to an $OD_{600}$ of 0.1 and incubated at 37 °C and 180 rpm until an $OD_{600}$ of 0.5 was reached. The culture was supplemented with 20 mM nicotinamide, 80 mM acetylated lysine, and 1 mM IPTG to induce protein expression and further incubated at 18 °C overnight. The bacterial cells were harvested by centrifugation (5,000 × g, 15 min, 4 °C) and stored at -20 °C overnight. The thawed bacterial pellet was resuspended in lysis buffer (300 mM NaCl, 50 mM PB at pH = 7.5, 10% glycerol, 20 mM nicotinamide), Turbonuclease (5 µL per L of culture), and a protease inhibitor cocktail tablet (Roche). The cells were lysed by sonication (70% amplitude,10 s pulse-10 s pulse off for 2.5 min). The lysate was cleared by centrifugation (18,500 × g, 30 min, 4 °C), and the supernatant was filtered through a 0.45 µm polyether sulfone filter and supplemented with 40 mM imidazole. The protein was purified by Ni-His-affinity chromatography using an ÄKTA start chromatography system (Cytiva) equipped with a 1 mL HisTrap column (Cytiva). Fractions as well as induction controls were analyzed by SDS page gel electrophoresis, and pure fractions were pooled and dialyzed against PBS containing 4% glycerol. The dialyzed fractions were concentrated to 1 mg $mL^{-1}$ using 3 kDa membrane filters (Amicon). The protein samples in PBS containing 20% glycerol were aliquoted and stored at –70 °C.

**Tobacco Etch Protease (TEV) cleavage**

The proteins *human* scGrx1 and *human* scGrx1-AcK20 were incubated with purified His10-SuperTev (Addgene #193833)[3] at a molar ratio of 1:3 in PBS containing 20% glycerol. The cleavage reactions were incubated at 37 °C overnight. The His-tag cleaved proteins were purified by Ni-His-affinity chromatography using an ÄKTA start chromatography system (Cytiva) equipped with a 1 mL HisTrap column (Cytiva), collecting the unbound protein fraction. Fractions were analyzed by SDS page gel electrophoresis, and pure fractions were pooled and dialyzed against PBS containing 4% glycerol. The dialyzed fractions were concentrated to 1 mg $mL^{-1}$ using 3 kDa membrane filters (Amicon). The protein samples in PBS containing 20% glycerol were aliquoted, snap-frozen and stored at –70 °C.

**Optical spectroscopy measurements for protein quantification**

Absorbance was measured on a Multiskan SkyHigh (ThermoFisher Scientific) with a µDrop Duo Plate. Protein concentration before labeling was determined based on the protein extinction coefficients or bicinchoninic acid (BCA) assay using the Pierce BCA Protein Assay Kit (ThermoFisher Scientific). Final protein concentrations of the labeled proteins were determined using the BCA assay.

**Protein bioconjugations**

The reactive dye was added to the protein sample in 1x PBS (pH 7.4). The reaction was incubated for 1-4 h at 37 °C and 180-600 rpm in a thermal shaker (ThermoFisher Scientific). For reaction monitoring, the sample was transferred into a conical LC-MS vial or a conical 96-well plate. The Protein masses and labeling efficiency were determined by deconvolving the total ion count (TIC+) peak using SAMMI (Cerno Biosciences).[4,5] The labeling conditions are specified in Table S1. Final protein concentrations of purified and labeled proteins were determined by bicinchoninic acid assay using the Pierce BCA Protein Assay Kit for labeled proteins (ThermoFisher Scientific).

**Size-exclusion chromatography (SEC)**

The HMSiR-labeled scGrx1 and scGrx1-ACK20 proteins were purified by gravity SEC chromatography (Cytiva, PD-10 column, Sephadex G-25 M) using PBS buffer. The fractions were analyzed with gel electrophoresis, pooled, and concentrated with 3 kDa membrane filters (Amicon) by centrifugation (14,000 × g, 10 min). The labeled protein samples were aliquoted, snap-frozen, and stored at –70 °C.

**Protein analysis by LC-MS**

Liquid chromatography electrospray-ionization mass spectrometry (LC-ESI-MS) was performed on a 1290 Infinity II ultra-high-performance liquid chromatography (UPLC; Agilent, USA) system connected to a single quadrupole mass spectrometer (SQ/MS; module type: G6125C; Agilent, USA). The samples were injected on an Acquity BEH C4 UPLC column (1.7 µm particle size, 2.1 × 50 mm, Waters) kept at 50 °C; with A: $H_2O$ + 1% $HCO_2H$ + 0.04% TFA and B: $CH_3CN$ + 0.1% $HCO_2H$ + 0.04% TFA, full scan MS in (+)-ESI mass ranges 500–1700 m/z. Data were analyzed using the OpenLab CDS software (Agilent) and calibrated with MassWorks software 4.0.0.0 (Cerno Biosciences) using a reference LC/MS tuning mix for ESI (Agilent), and deconvoluted using SAMMI (Cerno Biosciences)[5].

**Protein modification analysis by LC-MS/MS**

The sample preparation, measurement, and analysis have been performed by Dr. Chia-Wei Tan-Lin (staff member of the Functional Genomics Center Zurich—FGCZ). The HaloTag and SNAP-tag protein samples were alkylated with 15 mM chloroacetamide in aqueous buffer (10 mM Tris, 2 mM $CaCl_2$, pH 8.2) and incubated (30 min, 30 °C, 700 rpm) protected from light. All samples were enzymatically digested in a buffered trypsin solution (10 mM Tris and 2 mM $CaCl_2$, pH 8.2). Following enzymatic digests, the samples were dried. MS analysis was conducted by LC-MS/MS (Orbitrap, ThermoFisher Scientific). The digested samples were dissolved in aqueous 3% $CH_3CN$ with 0.1% formic acid, and the peptide concentration was estimated with the Lunatic UV/Vis absorbance spectrometer (Unchained Lab). Peptides were separated on an M-class UPLC and analyzed on an Orbitrap mass spectrometer (ThermoFisher Scientific). The acquired MS data were processed for identification using PEAKS Studio XPlus (Bioinformatic Solutions). The spectra were searched against the following modifications: oxidation (methionine), carbamidomethyl (cysteine), and the conjugated fluorophore (cysteine or lysine). The results were obtained and viewed in Scaffold 5 (Proteome Software) or Byonic (Protein Metrics).

**Full protein mass determination by UPLC/HR-ES-MS**

The sample preparation, measurement, and analysis have been performed by Dr. Serge Chesnov (staff member of the FGCZ). Samples were diluted in 1% TFA, clarified through an AttractFiltra RC Micro Spin column, and transferred to autosampler vials. Injection was performed onto a BioResolve Premier RP-mAb column (2.7 µm, 2.1 mm × 20 mm, 450 Å; Waters, USA) on an Acquity UPLC system. Desalting and separation were performed using a 30-minute gradient of buffer A (0.1% DFA in water) and buffer B (0.1% DFA in acetonitrile/75% 2-propanol) at 200 µL $min^{-1}$ and 60 °C. Eluate was introduced directly into a Synapt G2-Si mass spectrometer operated in positive-ion mode (m/z 400–5000 Da; 1 s scan, 0.1 s inter-scan delay; spray voltage 3 kV, cone voltage 50 V, source temperature 100 °C). Data were acquired with MassLynx 4.2 (Waters, UK), and deconvolution was carried out using the MaxEnt 1 algorithm with a resolution of 0.5 Da per channel and a Uniform Gaussian damage model (half-height width 0.5 Da).

### Liposome preparation and characterization

#### Lipid cake preparation

Lipid films were prepared in glass tubes by dissolving an appropriate amount of phospholipid in chloroform to reach a concentration of 2 mg $mL^{-1}$. To prepare one sample of vesicles, 100 µL of phospholipid solution was used per glass tube (i.e., 0.2 mg of lipid per tube). Depending on the experimental goal, the composition of the lipid cake varied. Typically, 16:0 biotinyl PE was added at 1:100% w/w (i.e., 1 µL of 1 mg $mL^{-1}$ solution to the 100 µL lipid solution of 2 mg $mL^{-1}$), labeled phospholipid was added at ~1:400% w/w (i.e., 0.25 µL of 1 mg $mL^{-1}$) to SMPC (i.e., 9.875 ul of a of 1 mg $mL^{-1}$ solution of SMPC in 38.876 µL of chloroform). The lipid solution was prepared in a batch for 5-10 lipid cakes, and 100 µL per lipid cake was distributed into clean glass tubes. Subsequently, the solution in the tubes was dried under a nitrogen flow overnight to form a thin lipid film. Tubes were sealed with Parafilm and stored at -20 °C.

**Vesicle preparation**

The lipid cake was wetted with 10 mM PB buffer, pH 7.4, or a solution of the protein of interest (POI) at 2 µM (HaloTag and $SNAP_f$-tag) or 4 µM (human scGrx1 and scGrx1-K20Ac) in the same buffer (for 0.2 mg of the lipid per tube). The wetted lipid cake was shaken in the thermoshaker (ThermoFisher Scientific) for 30 min at 1000 rpm and 4 °C above the transition temperature of the lipid (34 °C). An additional 500 µL of 10 mM PB was added to the vesicle solution. The speed of shaking was decreased to 800 rpm and the sample was incubated for an additional 30 min. To uniformize the vesicle size distribution, the mixture was extruded through a membrane with a pore size of 100 nm. The extruder (Avanti Lipids, Inc.) was cleaned, assembled, and equilibrated at 34 °C on the heating block. The vesicle mixture was taken up in a Hamilton glass syringe. The syringe was equilibrated for 10 min to reach 34 °C, then extruded 31 times (the sample was collected in the initially empty syringe).

**SEC of liposomes**

The SEC column (height of the bed was 21 cm, d = 1.5 cm) was prepared according to the manufacturer's instructions (Sepharose CL 6B, Cytiva). The column was equilibrated with 10 mM PB buffer, pH 7.4. Samples were loaded directly onto the column and passed through by gravity flow. Fractions of 1 mL were collected. For the specified column, vesicle, protein, and dye fractions are usually contained in 10-11 mL, 25-27 mL, and 34-36 mL, respectively. To confirm in which fraction the vesicles are present after purification with the SEC column, we ran thin-layer chromatography (TLC) in the solvent system 65:25:4 (v/v/v) chloroform:methanol:water. The lipids were visualized by potassium permanganate staining and irradiation with 356 nm if fluorescent lipids were added. To test their stability, the vesicle size was measured over 24 hours with timepoints at 1 h, 2 h, 4 h, 6 h, 21 h and 24 h by dynamic light scattering (DLS, Figure S1). The vesicles were sampled from the SEC column fraction kept at 25 °C or 30 °C in an Eppendorf tube in a thermoshaker. The data were analyzed using Kalliope (Anton Paar) and plotted using Python.

### Vesicle fusion and lipid exchange

We tested whether liposomes fused or exchange material between them. Vesicles were labeled with 2-dipalmitoyl-sn-glycero-3-phosphoethanolamine (DPPE) modified with dyes Atto520 ($\lambda_{ex}$ = 488 nm) or Atto647N ($\lambda_{ex}$ = 640 nm). Separate batches of liposomes were prepared, each labeled with one type of dye, and the batches were mixed before analysis using a flow nanoanalyzer (NanoFCM). Vesicles were measured separately at 1 h, 3 h, 6 h, and 24 h after mixing the two singly colored vesicles and storing them at room temperature (20 °C) in the dark. For measurement by flow nanoanalyzer, the fractions were diluted 1000 to 2000-fold. The samples were acquired for 1-2 min to record 4000-8000 events. The 488 nm laser was used at 20 mW, and the 640 nm laser at 40 mW. The data were analyzed using NF Profession (NanoFCM) and FlowJo (Becton, Dickinson and Company). Only minimal scrambling of dyes occurred upon mixing, with only modest further mixing over the course of 24 h (Figure S2).

### Cargo exchange between liposomes

Biotinylated liposomes were labeled with DPPE-Atto425 ($\lambda_{ex}$ = 405 nm) and loaded with unlabeled HaloTag. Separately, biotinylated liposomes were labeled with DPPE-Atto520 and loaded with HaloTag labeled with SiR-Br ($\lambda_{ex}$ = 640 nm), a far-red emitting rhodamine derivative (Figure S3). After SEC, the fractions containing the vesicles were combined in diluted form directly before the measurement on the microscopy slide in a 1:1 ratio (volume). To observe potential cargo exchange, the column fractions were combined 1:1 and kept in the dark at room temperature (20 °C) before sampling and diluting for the time point measurements (1 h, 6 h, 24 h). TIRF microscopy was used to analyze the colocalization HaloTag-SiR-Br with either DPPE-Atto520 or DPPE-Atto425 liposomes (Figure S3). The colocalization procedure is described in the computational section. The red-emitting HaloTag-SiR-Br complex colocalized mostly with DPPE-Atto520 liposomes but it also colocalized with spots that contained signals from both DPPE-Atto520 and DPPE-Atto425. This population can arise from either vesicles that exchanged lipids, which could be as high as ~10% (Figure S2), or DPPE-Atto520 and DPPE-Atto425 liposomes that coincide within a diffraction-limited area. A small population (<5%) of HaloTag-SiR-Br colocalized exclusively with DPPE-Atto425 liposomes, suggesting minimal cargo exchange between pre-formed vesicles (Figure S3). Of note, upon excitation at 640 nm, we observed several single-molecule-like signals outside of liposomes, which could be explained as impurities on the glass surface or proteins that escaped liposomes that might have ruptured during sample preparation. Overall, proteins remain mostly within their original liposome and by colocalizing the protein and vesicle signals, the provenance of the protein can be tracked.

#### Vesicle occupancy

Vesicles containing the fluorescent lipid DPPE-Atto520 were prepared using increasing protein concentrations of 2, 4, 8, and 16 μM of HaloTag labeled with SiR-Br. The samples were measured directly after preparation by fluorescence microscopy. Randomly selected fields of view were measured in the 488 nm (10 frames, 10% laser power, 30 ms) and in the 640 nm channel (1000 frames, 50% laser power, 30 ms) in TIRF mode. Vesicles and proteins were localized as described below in the computational section. Examples of vesicle localizations and distributions of proteins per vesicle at varying concentrations are displayed in Figure S3.

### Coverslip preparation

#### Cleaning

Round cover glasses (25 mm, 1.5 H, ThorLabs) were transferred to a glass staining dish that allowed spacing between slides. The coverslips were initially cleaned with MiliQ water to remove dust and water-soluble contaminants. The water was exchanged with 10% Deconex (Borer Chemie), and the slides were sonicated for 20 min, followed by thorough rinsing with MiliQ water until all traces of surfactant were removed, and sonicated in MiliQ water for 5 min. The slides were washed with acetone and sonicated in fresh acetone for 20 min. The coverslips were dried under a nitrogen flow and placed in the UV-ozone cleaner (PSDP, Novascan) on a piece of aluminum foil ~2 cm away from the light source, the upward-facing side was marked with a vertical line on the bottom right. They were irradiated for 10 min at 25 °C and then rested for 30 min in the closed chamber.

#### Coverslips modification and vesicle deposition

After the UV-ozone treatment, clean slides were transferred to a light-protected pipette tip box and 150 µL of 0.1 mg $mL^{-1}$ biotinyl PLL-g-PEG and PLL-g-PEG solutions mixed in a ratio of 1:50 were deposited onto the etched side of the coverslip. The solution was incubated at room temperature for 30 min. After incubation, the sample was removed and rinsed with MiliQ water. The coverslips were then covered with 0.1 mg $mL^{-1}$ Neutravidin solution in 10 mM PB buffer, pH 7.4, for at least 30 min. After washing with MiliQ water, the coverslip was fixed in a cell chamber (Aireka Scientific), the collected vesicles were deposited directly on the slides, and they were incubated for at least 15 min. Finally, we washed the slide with the same PB buffer by removing the solution (not to dryness) and diluting them at least five times. In multicolor experiments, the vesicles were mixed in the given ratio before the application to the modified coverslips and washed analogously.

### Fluorescence imaging

#### General TIRF method

A Nikon N-STORM microscope equipped with an EMCCD camera iXon 888 (Andor) was employed. TIRF images were collected with an SR HP Apochromat TIRF 100x 1.49 N. A. oil immersion objective lens. A piezo-electronic focus-lock system (perfect focus system) was used to prevent axial drift during data acquisition. Laser lines for fluorescence imaging were 405 nm, 488 nm, 561 nm, and 638 nm. Appropriate filter cubes were configured within the light path. The microscope was operated using the NIS Elements software.

#### Single-molecule imaging in coverslips

Coverslips fixed in the cell chamber were imaged in TIRF mode with a 638 nm laser, 30 ms exposure time for 8000 or 6000 frames followed by an acquisition with a 488 nm laser, 30 ms exposure time for 10 frames. The conditions were kept constant for all acquisitions. For all proteins, single-molecule measurements were carried out on different days, all these signals were mixed to create the datasets for further analysis.

### Small-molecule synthesis

#### General methods

All reagents were purchased from commercial sources and used as received. Anhydrous solvents were procured from Acros Organics and used as received. All solvents used in the preparation of the coverslips were HPLC grade. Flash column chromatography was carried out using prepacked Buchi Reveleris $SiO_2$ on a a Biotage® Selekt System. Reverse-phase HPLC was conducted on an Agilent 1290 Infinity II System with a $C_{18}$-column (Phenomenex). TLCs were run on $SiO_2$-60-$F_{254}$ plates (Sigma-Aldrich) and visualized using UV light at 254 nm and 366 nm or permanganate stain. Nuclear magnetic resonance (NMR) spectra were acquired on a Bruker Ultrashield 400 instrument. $^1$H NMR chemical shifts are reported in ppm relative to $SiMe_4$ (δ = 0) and were referenced internally with respect to residual protons in the solvent (δ = 7.26 for $CDCl_3$, δ = 1.94 for $CD_3CN$, δ = 3.31 for $CD_3OD$ and δ = 3.58 for THF). Coupling constants are reported in Hz. $^{13}$C NMR chemical shifts are reported in ppm relative to $SiMe_4$ (δ = 0) and were referenced internally with respect to solvent signal (δ = 77.16 for $CDCl_3$ and δ = 1.32 for $CD_3CN$). Preliminary peak assignments are based on calculated chemical shifts. Multiplicities were abbreviated as follows: singlet (s), doublet (d), triplet (t), quartet (q), multiplet (m). High-resolution MS (HRMS) was conducted by the staff at the ISIC Mass Spectrometry facility (at EPFL) employing a Waters Xevo G2-2 quadrupole time-of-flight (QTOF) or Agilent Technologies 6530 Accurate-Mass QTOF LC-MS or by staff at Mass Spectrometry Laboratory of the Department of Chemistry at the UZH employing a Dionex Ultimate 3000 ultra-HPLC system (ThermoFischer Scientific) connected to a QExactive MS with a heated ESI source (ThermoFisher Scientific). Agilent 1290 Infinity II LC System with an Agilent InfinityLab LC/MSD. IUPAC names of all compounds are provided and were determined using CS ChemBioDrawUltra 22.2.

*3,3'-(Dimethylsilanediyl)bis(N,N-dimethylaniline) (**S1**)*

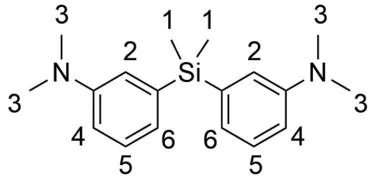

Compound **S1** (5 g, 25 mmol) was dissolved in dry THF (60 mL) and the solution was cooled to –78 °C. *n*-BuLi (17.2 mL, 27.5 mmol, 1.6 M) was added dropwise and the mixture was stirred at –78 °C for 2 h. Dichlorodimethylsilane (1.52 mL, 12.5 mmol) was added dropwise and the mixture was stirred for 2 h at 25 °C. Brine (40 mL) and $H_2O$ (10 mL) were added and the mixture was extracted three times with EtOAc. The combined organic phases were dried over $MgSO_4$ and concentrated onto Celite. The crude was purified by flash column chromatography ($SiO_2$; hexane to hexane/EtOAc 9:1) to give the product **S1** as a light-yellow oil (3.6 g, 50%). $^1$H NMR (400 MHz, $CDCl_3$) δ = 7.31–7.19 (m, 2H, H5), 6.97–6.90 (m, 4H, H4, H2), 6.78 (dd, *J* = 8.3, 2.8, 2H, H6), 2.94 (s, 12H, H3), 0.55 (s, 6H, H1) ppm. $^{13}$C NMR (101 MHz, $CDCl_3$) δ = 154.75, 141.11, 131.49, 125.30, 120.50, 116.20, 42.84, 0.84 ppm.

*3,3'-(Dimethylsilanediyl)bis(4-bromo-N,N-dimethylaniline) (**S2**)*

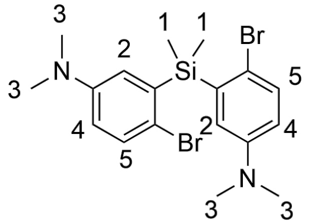

Compound **S1** (3.00 g, 0.01 mol) was dissolved in $CH_3CN$ (75 mL) in a flame-dried flask. The solution was cooled to 0 °C and NBS (3.75 g, 0.021 mmol) was added in small portions. After complete addition, the solution was stirred for 1.5 h at 25 °C then sat. $NaHCO_3$ was added to neutralize the solution. The organic phase was washed with water. The combined aqueous phases were extracted twice with $CH_2Cl_2$. All $CH_2Cl_2$ fractions were combined, dried over $MgSO_4$, and concentrated under reduced pressure. The residue was purified by flash column chromatography. ($SiO_2$; hexane: EtOAc 95:2) yielding the white product **S2** (2.69 g, 59%). $^1$H NMR (400 MHz, $CDCl_3$) δ = 7.36 (d, *J* = 8.8, 2H, H5), 6.85 (d, *J* = 3.2, 2H, H2), 6.61 (dd, *J* = 8.7, 3.2, 2H, H4), 2.89 (s, 12H, H3), 0.77 (s, 6H, H1) ppm. $^{13}$C NMR (101 MHz, $CDCl_3$) δ = 149.02, 138.86, 133.10, 121.92, 116.94, 115.40, 40.70, -0.78 ppm. HRMS (ESI/QTOF) $[M+H]^+$ calcd. for $[C_{18}H_{25}Br_2N_2Si]^+$: 455.0148; found 455.0146.

*3,7-bis(Dimethylamino)-5,5-dimethyl-3'H,5H-spiro[dibenzo[b,e]siline-10,1'-isobenzofuran]-5'-carbonitrile (**S3**)*

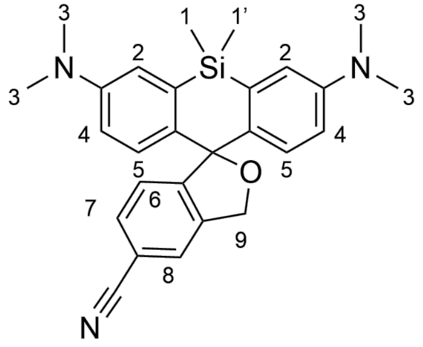

Compound **S2** (450 mg, 0.986 mmol) and 5-cyanophthalide (345 mg, 2.17 mmol) were dried under reduced pressure in two flame-dried flasks and dissolved in dry THF (11.25 mL and 22.5 respectively). The solution of 3,3'-(dimethylsilanediyl)bis(4-bromo-*N,N*-dimethyl aniline) was cooled to –78 °C using an acetone/dry ice bath. *t*-BuLi solution (2.55 mL, 4.34 mmol, 1.7 M) was added, and the reaction mixture was stirred at –78 °C for 30 min. The phthalide solution was added dropwise over 60 min. The reaction mixture was allowed to warm to 20 °C and was stirred for 16 h. Saturated $NH_4Cl$ was added to the mixture and was stirred for 30 min. The aqueous phase was extracted three times with $CH_2Cl_2$. The organic phase was dried over $Na_2SO_4$, concentrated under reduced pressure, loaded onto Celite, and purified by flash chromatography ($SiO_2$; hexane/EtOAc 95:5 to hexane/EtOAc 8:2). Product **S3** was obtained as a white/blue powder (130 mg, 30%). $^1$H NMR (400 MHz, $CDCl_3$) δ = 7.61 (s, 1H, H8), 7.51 (d, *J* = 7.9, 1H, H7), 7.11 (s, 1H, H6), 6.95 (d, *J* = 2.9, 2H, H2), 6.92 (dd, *J* = 8.9, 1.2, 2H, H5), 6.63 (dd, *J* = 8.9, 2.9, 2H, H4), 5.32 (s, 2H, H9), 2.96 (d, J=0.8, 12H, H3), 0.62 (d, *J* = 0.8, 3H, H1 or H1'), 0.54 (d, *J* = 1.2, 3H, H1 or H1') ppm. $^{13}$C NMR (101 MHz, $CDCl_3$) δ 152.29, 149.01, 140.48, 137.14, 135.28, 131.72, 128.55, 125.49, 125.37, 119.16, 116.69, 113.97, 111.11, 92.85, 72.22, 40.53, 0.56, -0.76. HRMS (ESI-MS) $[M+H]^+$ calcd. for $[C_{27}H_{30}N_3OSi]^+$: 440.21527, found 440.21522.

*5'-(Aminomethyl)-N3,N3,N7,N7,5,5-hexamethyl-3'H,5H-spiro[dibenzo[b,e]siline-10,1'-isobenzofuran]-3,7-diamine (**S4**)*

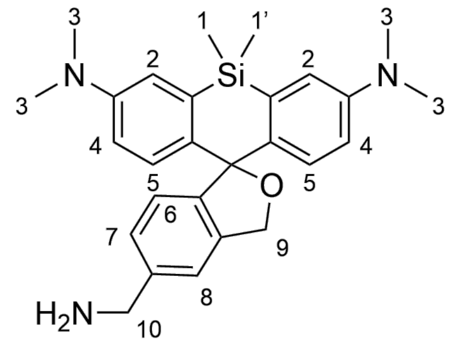

Compound **S3** (130 mg, 0.3 mmol) was transferred to a flame-dried reaction flask with 2.4 mL of THF and cooled to 0 °C. The flask was flushed with nitrogen. The solution was stirred, and a $LiAlH_4$ solution (0.65 mL, 0.65 mmol, 1 M) at 0 °C was added dropwise. The color changed from green-blue to yellow to dark orange to brownish green. After 2 h the reaction mixture was left warm to 20 °C. The reaction mixture was neutralized with a few drops of saturated $NH_4Cl$. It was extracted three times with $CH_2Cl_2$. The organic layer was dried with brine and then over $Na_2SO_4$, concentrated under reduced pressure, and loaded onto Celite. The crude was purified by flash chromatography ($SiO_2$; $CH_2Cl_2/CH_3OH$ 1:0 to $CH_2Cl_2/CH_3OH$ 92:8), yielding compound **S4** as a white powder (80 mg, 66%). $^1H$ NMR (400 MHz, $CDCl_3$) δ = 7.26 (s, 1H, H8, covered by $CHCl_3$ peak), 7.18 (d, $J$ = 7.9, 1H, H6), 7.03 (d, $J$ = 7.7, 1H, H7), 6.98 (d, $J$ = 5.5, 2H, H5), 6.96 (s, 2H, H2), 6.61 (dd, $J$ = 8.9, 2.9, 2H, H4), 5.21 (s, 2H, H9), 3.93 (s, 2H, H10), 2.94 (s, 12H, H3), 0.61 (s, 3H, H1 or H1'), 0.54 (s, 3H, H1or H1') ppm. $^{13}C$ NMR (101 MHz, $CDCl_3$): $\delta$ = 148.12, 144.68, 139.80, 137.76, 134.92, 127.87, 126.10, 124.11, 119.37, 116.18, 113.18, 91.75, 71.53, 45.23, 39.93, 0.00, -1.84 ppm. HRMS (ESI-QTOF) $[M+H]^+$ calcd. for $[C_{27}H_{34}N_3OSi]^+$ 444.2471, found 444.2474.

*N-((3,7-bis(Dimethylamino)-5,5-dimethyl-3'H,5H-spiro[dibenzo[b,e]siline-10,1'-isobenzofuran]-5'-yl)methyl)-2-iodoacetamide (**HMSiR-IA**)*

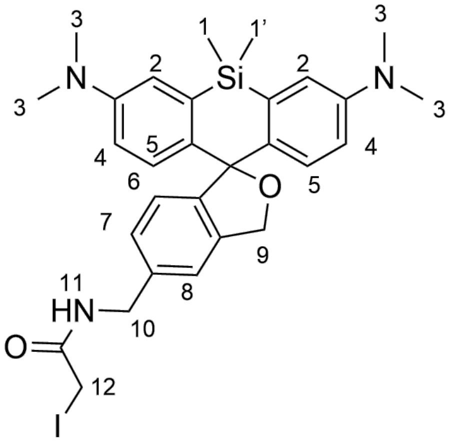

TSTU (33.9 mg, 0.113 mmol) and DIPEA (14.6 mg, 0.113 mmol) were added to a solution of iodoacetic acid (20.9 mg, 0.113 mmol) in $CH_3CN$ (1.2 mL). After the reaction mixture was stirred at 25 °C for 1 h, a solution of compound **S4** (25 mg, 0.056 mmol) in $Na_2CO_3$ saturated $CH_3CN$ was added. After 1 h, 1 M HCl (1 mL) was added to the mixture, before removing the $CH_3CN$ under reduced pressure. The residue was dissolved in $CH_2Cl_2$, more water was added and extracted three times in total with $CH_2Cl_2$. The organic phase was washed with brine and dried over $Na_2SO_4$, concentrated under reduced pressure. The crude was dissolved in $CH_3CN$ /water 50:50 with 0.1% TFA and purified by HPLC ($SiO_2$-$C_{18}$; $CH_3CN$ + 0.1% TFA/$H_2O$ + 0.1% 2:8 to $CH_3CN$ + 0.1% TFA/$H_2O$ + 0.1% 9:1) resulting in 12 mg of material **HMSiR-IA** (17%). $^1H$ NMR (400 MHz, $CD_3OH$) δ = 7.67 (s, 1H, H8), 7.41 (d, $J$ = 7.9, 1H, H7), 7.35 (d, $J$ = 2.8, 2H, H2), 7.11 (d, $J$ = 7.7, 1H, H6), 7.09 (d, $J$ = 9.6, 2H, H5), 6.74 (dd, $J$ = 9.7, 2.8, 2H, H4), 4.52 (s, 2H, H9), 4.31 (s, 2H, H10), 3.80 (s, 2H, H12), 3.34 (s, 12H, H3, H3'), 0.60 (d, $J$ = 4.2, 6H, H1, H1') ppm. $^{13}C$ NMR (101 MHz, $CD_3OH$) δ = 171.49, 169.59, 155.78, 149.42, 142.55, 141.02, 140.90, 137.34, 130.55, 128.75, 127.24, 126.93, 122.11, 115.06, 62.37, 44.31, 40.88, -1.09, -1.35, -2.18. HRMS (ESI-MS) $[M+H]^+$ calcd. for $[C_{29}H_{35}IN_3O_2Si]^{+:}$ 612.15377, found 612.15482.

*4-Bromo-3-(dibromomethyl)benzoic acid (**S5**)*

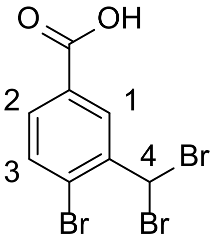

4-Bromo-3-methylbenzoic acid (10 g, 46.5 mmol), NBS (8.28 g, 46.5 mmol), and AIBN (99.3 mg, 605 µmol) were suspended in α, α, α-trifluorotoluene (149 mL) and the solution was heated to 110 °C. After it was heated more AIBN (50 mg) and 1 additional equivalent of NBS was added. AIBN (50 mg) was further added every 3 to 4 h for a total of 24 h. The mixture was cooled to 25 °C and 50 mL of heptane was added dropwise. This mixture was stirred in a salt ice bath at –10 °C for 3 h. The solid was concentrated by filtration, washed with heptane at –20 °C, and dried under reduced pressure to give compound **S5** as a white solid (15.5 g, 89%). $^{1}$H NMR (400 MHz, DMSO-$d_6$) δ = 8.47 (s, 1H, H3), 7.81 (d, $J$ = 1.2 Hz, 2H, H1, H2), 7.39 (s, 1H, H4). $^{13}$C NMR (101 MHz, DMSO-$d_6$) δ = 165.86, 140.22, 133.84, 131.88, 131.30, 131.04, 40.57. HRMS (ESI/QTOF) [M-H]$^-$ calcd. for $[C_8H_4Br_3O_2]^-$: 368.7767, found 368.7765.

*4-Bromo-3-(hydroxymethyl)benzoic acid (**S6**)*

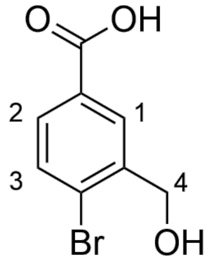

Compound **S5** (1 g, 4.37 mmol) was dissolved in THF (20 mL) and then cooled to 0 °C. The mixture was stirred between 0 and 5 °C for 5 h. Then 1 M HCl was added, and the mixture was fully concentrated under reduced pressure. The residue was dissolved in EtOAc and 1 M HCl. The aqueous acidic phase was removed, and the organic phase was washed with water and brine, dried over $Na_2SO_4$, and concentrated under reduced pressure. Little product was obtained (clean by NMR, 70 mg) therefore the aqueous phase was further acidified by the addition of 1 M HCl. The aqueous phase was extracted twice with $CH_2Cl_2$, dried over $Na_2SO_4$, and concentrated under reduced pressure. The combined product **S6** was a white solid (0.94 g, 93%). $^{1}$H NMR (400 MHz, $CD_3OD$) δ = 8.21 (dt, $J$ = 2.0, 0.9, 1H, H1), 7.81 (ddt, $J$ = 8.4, 2.2, 0.7, 1H, H2), 7.66 (d, $J$ = 8.3, 1H, H3), 4.69 (t, $J$ = 0.8, 2H, H4) ppm. $^{13}$C NMR (101 MHz, $CD_3OD$) δ = 169.10, 142.40, 133.68, 131.52, 130.71, 130.29, 128.00, 64.34 ppm. HRMS (ESI/QTOF) [M-H]$^-$ calcd. for $[C_8H_6BrO_3]^-$: 228.9506, found 228.9504.

*4-Bromo-3-formylbenzoic acid (**S7**)*

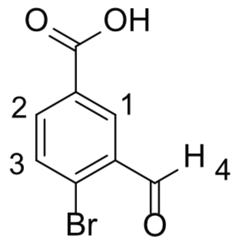

Compound **S6** (4.77 g, 12.8 mmol) was added to a stirred solution of 10% aqueous $Na_2CO_3$ (84 mL). The solution was heated to 70 °C and stirring was continued for 5 h. The resulting precipitate was removed by filtration while the solution was still hot. The filtrate was carefully acidified with HCl until pH 1 was reached and extracted twice with EtOAc. The combined organic phase was washed with brine, dried over $Na_2SO_4$, filtered, and evaporated to give **S7** as a white solid (9.5 g, 96%). $^1H$ NMR (400 MHz, $(CD_3)_2CO$) δ = 10.37 (s, 1H, H4), 8.47 (d, *J* = 2.2 Hz, 1H, H1), 8.17 (d, *J* = 8.3 Hz, 1H, H2), 7.95 (d, *J* = 8.3 Hz, 1H, H3). $^{13}C$ NMR (101 MHz, $(CD_3)_2CO$) δ = 190.37, 165.17, 135.62, 134.62, 133.88, 130.80, 130.59. HRMS (ESI/QTOF) $[M-H]^-$ calcd. for $[C_8H_4BrO_3]^-$: 226.9349, found 226.9356.

*tert-Butyl 4-bromo-3-(tert-butoxymethyl)benzoate (**S8**)*

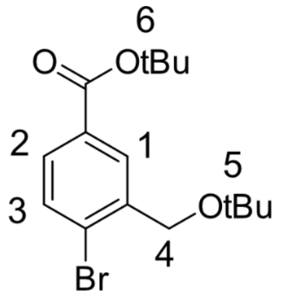

Concentrate $H_2SO_4$ (378 µL, 7.1 mmol) was added to a vigorously stirred suspension of anhydrous Epsom salt (3.42 g, 28.4 mmol) in $CH_2Cl_2$ (60 mL). The mixture was stirred for 15 min at 25 °C then compound **S7** (820 mg, 3.55 mmol) and *tert*-butanol (3.37 mL) were added. The flask was closed, and the mixture was stirred for 5 d at 25 °C. After the addition of saturated $NaHCO_3$ the reaction mixture was extracted twice with EtOAc, washed with brine, dried over $Na_2SO_4$, filtered, and concentrated under reduced pressure. The crude was purified by flash column chromatography ($SiO_2$; hexane to hexanes/EtOAc 91:9) to give the product **S8** as a white oil, that crystallized when left unmoved (0.972 g, 80%).$^1H$ NMR (400 MHz, $CDCl_3$) δ = 8.14 (dd, *J* = 2.1, 1.0, 1H, H1), 7.71 (dd, *J* = 8.3, 2.3, 1H, H2), 7.55 (d, *J* = 8.2, 1H, H3), 4.50 (s, 2H, H4), 1.59 (s, 9H, H6), 1.32 (s, 9H, H5) ppm. $^{13}C$ NMR (101 MHz, $CDCl_3$) δ = 165.40, 139.48, 132.34, 131.42, 130.11, 129.33, 127.26, 81.40, 74.12, 63.53, 28.30, 27.77 ppm. HRMS (ESI/QTOF) $[M+Na]^+$ calcd. for $[C_{16}H_{23}BrNaO_3]^+$: 365.0723, found 365.0721.

*3,7-bis(Dimethylamino)-5,5-dimethyldibenzo[b,e]silin-10(5H)-one (**S9**)*

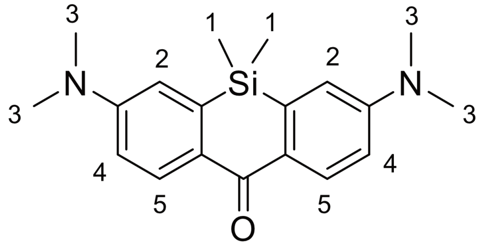

*t*-BuLi solution (5.15 mL, 8.76 mmol, 1.7 M) was added dropwise to a solution of compound **S2** (0.999 g, 2.19 mmol) in anhydrous THF (40 mL) at –78 °C. The resulting bright yellow solution was stirred at –78 °C for 1.5 h. Neat dimethyl carbamoyl chloride (0.259 g, 2.41 mmol) was then added dropwise. The resulting mixture was stirred at –78 °C for 50 min, then allowed to warm up to 25 °C and left stirring for 16 h. The reaction was stopped with saturated $NH_4Cl$ solution (25 mL), water was added to dissolve solids, and the mixture was extracted with EtOAc three times. The combined extracts were dried over $Na_2SO_4$, filtered, and the product was isolated by flash column chromatography ($SiO_2$; hexane to hexane/EtOAC 9:1) to give 650 mg (92%) of the ketone **S9** as a bright green crystalline solid. $^1$H NMR (400 MHz, $CDCl_3$) δ = 8.40 (d, *J*=8.9, 2H, H2), 6.84 (dd, *J*=8.9, 2.7, 2H, H4), 6.79 (d, *J*=2.8, 2H, H5), 3.10 (s, 12H, H3), 0.47 (s, 6H, H1). $^{13}$C NMR (101 MHz, $CDCl_3$) δ = 185.41, 151.59, 140.61, 131.76, 129.83, 114.38, 113.30, 40.18, -0.85. HRMS (ESI/QTOF) $[M+H]^+$ calcd. for $[C_{19}H_{25}N_2OSi]^+$: 325.1731, found 325.1763.

*3,7-bis(Dimethylamino)-5,5-dimethyl-3'H,5H-spiro[dibenzo[b,e]siline-10,1'-isobenzofuran]-5'-carboxylic acid (**S10**)*

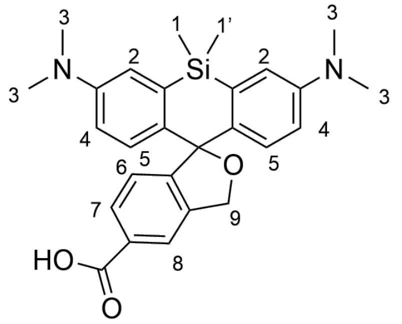

Compound **S8** (291 mg, 0.847 mmol) was dissolved in THF (5 mL) under $N_2$ atmosphere in a pre-dried flask and cooled to –78 °C. Then, *t*-BuLi solution (0.45 mL, 0.77 mmol, 1.7 M) was added and the reaction was stirred at the same temperature for 10 min. Compound **S9** (50 mg, 0.154 mmol) dissolved in anhydrous THF (2 mL) and was added. The resulting mixture was warmed to ambient temperature and stirred for 16 h. Saturated $NH_4Cl$ was added to the reaction mixture and extracted three times with EtOAc and once with $CH_2Cl_2$. The combined organic phases were dried over $Na_2SO_4$, and the solvent was removed under reduced pressure. The resulting intermediate was dissolved in TFA (3.5 g, 30.8 mmol) and the solution was stirred at 20 °C for 3.5 h. After removal of all volatiles under reduced pressure the crude was purified by reverse phase HPLC ($SiO_2$-$C_{18}$; $CH_3CN$ + 0.1% TFA/$H_2O$ + 0.1% TFA 1:9 to $CH_3CN$ + 0.1% TFA/$H_2O$ + 0.1% TFA 95:5) yielding 56 mg of the product **S10** as a blue solid (56%). $^1$H NMR (400 MHz, $CD_3OD$) δ = 8.41 (s, 1H, H8), 8.11 (dd, *J* = 7.8, 1.7, 1H, H7), 7.37 (s, 2H, H2), 7.28 (d, *J* = 7.9, 1H, H6), 7.04 (d, *J* = 9.6, 2H, H5), 6.78 (dd, *J* = 9.6, 2.9, 2H, H4), 4.36 (s, 2H, H9), 3.35 (s, 12H, H3), 0.61 (d, *J* = 3.7, 6H, H1) ppm. $^{13}$C NMR (101 MHz, $CD_3OD$) δ = 169.14, 167.95, 155.83, 149.40, 142.97, 142.12, 141.52, 132.78,

130.66, 129.46, 129.22, 128.15, 122.33, 115.32, 62.02, 40.93, -1.09, -1.33 ppm. HRMS (ESI/QTOF) $[M+H]^+$ calcd. for $[C_{27}H_{31}N_2O_3Si]^+$: 459.2099; found 459.2097.

*N-(2-(2-((6-Chlorohexyl)oxy)ethoxy)ethyl)-3,7-bis(dimethylamino)-5,5-dimethyl-3'H,5H-spiro[dibenzo[b,e]siline-10,1'-isobenzofuran]-5'-carboxamide (HMSiR-CA)*

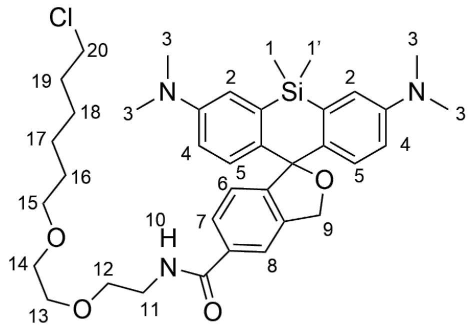

The protected chloroalkane ligand (16.5 mg, 0.051 mmol) was dissolved in 4 M HCl in dioxane (0.35 mL) and was stirred at 20 °C for 1 h. A saturated $NaHCO_3$ solution was added to the reaction, and the mixture was extracted with $CH_2Cl_2$. The combined organic phases were dried over $Na_2SO_4$, the solvent was evaporated, and the product was directly used in the next step. In a flame-dried Schlenk flask under $N_2$-atmosphere, compound **S10** (10 mg, 0.017 mmol) was dissolved in dry DMF (2 mL), then DIPEA (65.9 mg, 0.51 mmol) was added and the reaction was stirred at 20 °C for 10 min, then HATU (12.9 mg, 0.034 mmol) and the crude from the previous step was added and the reaction was stirred at 20 °C for 1.5 h. The solvent was evaporated, and the crude was stored in the freezer overnight. The crude was purified by reverse phase HPLC ($SiO_2$-$C_{18}$; $CH_3CN$ + 0.1% TFA/$H_2O$ + 0.1% TFA 5:95 to $CH_3CN$ + 0.1% TFA/$H_2O$ + 0.1% TFA 65:35). The $CH_3CN$ was evaporated as well as parts of the $H_2O$, then sat. $Na_2CO_3$ was added until no bubbles were observed. The aqueous phase was extracted with $CH_2Cl_2$, the organic phases were dried over $NaSO_4$ and concentrated under reduced pressure. The oily solid (10 mg) was redissolved in $CH_3CN/H_2O$ 1:1 and lyophilized giving 9 mg of compound **HMSiR-CA** (80%). $^1H$ NMR (400 MHz, $CD_3OD$) δ = 7.81 (s, 1H, H8), 7.64 (d, *J* = 8.2, 1H, H7), 7.00 (dd, *J* = 2309.9, 12.6, 2H, H2), 6.99 (d, *J* = 45.5, 2H, H5), 6.90 (d, *J* = 8.0, 1H, H6), 6.70 (dd, *J* = 8.9, 2.9, 2H, H4), 5.39 (s, 2H, H9), 3.68–3.61 (m, 4H, H19, H11), 3.60–3.54 (m, 4H, H10, H14), 3.45 (t, *J* = 6.6, 4H, H12, H13), 2.92 (s, 12H, H3), 1.73–1.58 (m, 2H, H18), 1.57–1.46 (m, 2H, H15), 1.42–1.30 (m, 4H, H16, H17), 0.59 (s, 3H, H1 or H1'), 0.49 (s, 3H, H1 or H1') ppm. $^{13}C$ NMR (101 MHz, $CD_3OD$) δ = 170.03, 152.59, 150.53, 139.73, 139.44, 135.44, 135.03, 129.92, 127.97, 124.72, 121.76, 117.55, 115.66, 94.17, 73.83, 72.18, 71.29, 71.13, 70.47, 45.67, 41.02, 40.81, 33.68, 30.49, 27.67, 26.41, 0.22, -0.20 ppm. HRMS (ESI-QTOF) $[M+H]^+$ calcd. for $[C_{37}H_{51}ClN_3O_4Si]^+$ 664.3332, found 664.3329.

### Computational methods

#### Analysis of cargo exchange between liposomes

The script reads raw ND2 movies directly and runs Picasso[10] maximum likelihood estimator (MLE) localization on each frame of each relevant channel (405, 515, 640 nm) before colocalization. Multi-position ND2 files (containing several FOVs) are handled transparently: each position is extracted and localized independently. Localization uses the parameters recorded in the original Picasso YAML sidecar files: box size 7 px, minimum net gradient 20 000, MLE fitting with convergence criterion ε = 0.001 and maximum 1000 iterations.

For the colocalization step, all protein localizations and all vesicle localizations from a given FOV (pooled across all frames) are treated as spatial point clouds. For each protein localization, the nearest vesicle localization in each channel is found via a KD-tree query[6]. A protein localization is classified as localizing with a DPPE-Atto520 vesicle if it is within `max_dist` pixels (default 3.0) of any Atto520 (C3) localization. A protein is assigned to DPPE-Atto425 if it is within `max_dist` pixels of any Atto425 (C4) localization. A protein is assigned as free if it is not within `max_dist` of either vesicle type. A protein localization may be simultaneously associated to both DPPE-Atto45 and -Atto520 if both vesicle types are present within `max_dist`. Such cases are counted in a third category (both). The free fraction is the complement of the union: protein localizations that are within `max_dist` of neither vesicle type. Classification counts are summed across all FOVs within a slide and expressed as percentages of the total protein localization count for that slide.

Note that this analysis operates on raw Picasso[10] localizations, not on linked or tracked molecules. The same physical molecule contributes one localization per frame in which it was detected, so the counts are weighted by the number of frames each molecule was detected in. For the purpose of quantifying population-level transfer fractions, this weighting is acceptable because it is applied uniformly across all slides and timepoints. Results are aggregated per slide (summing counts across all FOVs) and then sorted by timepoint. The analysis script writes two comma-separated-value (CSV) files: one per replicate and a combined file with a replicate column for joint analysis.

#### Analysis of liposome occupation

To characterize how many protein molecules are encapsulated per vesicle as a function of protein loading concentration, a photobleaching step-counting approach was used. Vesicles were immobilized on a glass surface and imaged under continuous 640 nm illumination. Each fluorescently labeled protein molecule undergoes irreversible single-step photobleaching, producing a discrete downward step in the integrated vesicle fluorescence trace. The number of steps observed in the trace is therefore equal to the number of labeled protein molecules present in that vesicle at the start of imaging[7].

Before step detection, the algorithm preprocesses traces per vesicle to perform background subtraction. The mean of the last 50 frames of the raw pixel-sum trace is subtracted from every frame (corresponding to the noise floor after all fluorophores have bleached). Then, the background-subtracted trace is divided by the ROI area (boxsize2 = 49 pixels) to obtain per-pixel signal normalization. Photobleaching steps are then detected using the `quickpbsa` package (v2021.0.1), which implements the Kalafut–Visscher (KV) algorithm[8]. The preliminary KV pass fits a piecewise-constant model to the trace using a maximum-likelihood criterion, with the number of steps determined by minimizing the Schwarz information criterion (SIC). A refinement pass then resolves simultaneous multi-molecule bleaching events using a Bayesian posterior criterion.

Each analyzed vesicle is assigned a quality flag, with flag = 1 meaning that KV detected one or more clearly resolved photobleaching steps. The molecule count (occupation) is stored in the `fluors_kv` field of the results CSV. Only these vesicles are used for the occupation histogram. Flag = −1 means that KV found zero steps, indicating either that all proteins were already bleached before imaging began, that no protein was encapsulated, or that the signal was too weak to resolve individual steps. These vesicles are excluded from the step-counting result. Other negative flags are additional quality-control rejections (e.g., flag = −3 for incomplete bleaching, flag = −7 when the refinement search space exceeds the combination cutoff).

Vesicles that contain no protein are not detected in the 640 nm channel and therefore do not appear in the KV result CSV at all. To correctly compute the full occupation distribution (including the zero-protein fraction), these empty vesicles must be counted separately. Their number is inferred as the difference between the total vesicle cluster count from Picasso and the number of vesicles with flag = 1. Vesicle clusters are detected from the 488 nm (DPPE-Atto520) channel by density-based clustering[9] applied to all localizations across the full movie; each cluster represents one vesicle. The resulting empty-vesicle rows (occupation = 0) are appended to the DataFrame before computing distributions. The clustering results file occasionally contains duplicate entries (if the pipeline was re-run on the same data), so the script reads only the first $N_{FOV}$ entries, where $N_{FOV}$ is determined by counting the per-FOV trace pickle files in the directory.

### Trace extraction

*Step 1. Gradient threshold optimization*

Movies were acquired as described in the Fluorescence Imaging section. To construct single-molecule traces of fluorescence vs time, we employed Picasso[10] for single-molecule localization per frame. Critically, whether Picasso finds a molecule (or liposome) depends on the gradient used for localization. Too low a threshold floods the field of view with noise-driven false spots; too high a threshold misses genuine single-molecule signals. Because the optimal value is different for molecules or liposomes, and it depends on the specific camera settings, excitation intensity, and dye brightness, our algorithm determines it independently for each channel and each experiment using Bayesian optimization. We maintained the size of the localization box fixed at 5 × 5 pixels across all conditions to ensure that pixel-summed intensities are directly comparable between proteins and experiments. For each candidate gradient value, the pipeline runs Picasso localization on the first 1,000 frames of one randomly selected test movie, links the resulting spot positions into traces (see Step 3), and evaluates a composite objective score:

$$\mathcal{O} = N_{\text{single}} \cdot \left(1 + \frac{N_{\text{IN}}}{N_{\text{single}}}\right) \cdot (1 - r_{\text{overlap}})^2$$

where $N_{\text{single}}$ is the number of non-overlapping single-molecule regions of interest (ROIs), $N_{\text{IN}}$ is the subset of single ROIs that colocalize with a vesicle in the 488 nm channel (labeled IN, see Step 3), and $r_{\text{overlap}}$ is the fraction of all ROIs that share pixels with another ROI. Therefore, this objective function maximizes the number of non-overlapping single-molecule localizations that are associated with a liposome. The code can be run in single-channel mode (no vesicle reference), in which $N_{\text{IN}}$ is omitted and the objective reduces to $N_{\text{single}} \bullet (1\text{-}r_{\text{overlap}})^2$.

The optimization uses the Tree-structured Parzen Estimator (TPE) sampler implemented in Optuna[11]. TPE models the distribution of good and bad parameter settings separately, and samples new candidates preferentially from the region where the ratio p(good)/p(bad) is highest. This is substantially more efficient than random or grid search and we found that 20 trials are sufficient to identify a near-optimal gradient value over a discrete search space (default: 10,000–80,000 in steps of 10,000 for the protein channel; 1,000–10,000 in steps of 1,000 for the vesicle channel). By default, the optimizer runs in per-experiment mode: one gradient set is found for all proteins within the same experimental session (same day, same optical path, same illumination intensity). One test movie per protein is selected at random, and the objective score for each trial is the mean (or optionally minimum) over all proteins in that session. This shared optimization guarantees that the relative intensity scale is consistent across proteins in downstream comparisons. Figure S7 displays composite maximum projections for a test movie with ROI boxes and objective score for a few parameters.

*Step 2. Single-molecule localization*

Single-molecule positions were determined by the Picasso software package using its MLE algorithm[10]. For each candidate spot (detected via the net gradient filter described above), Picasso fits a 2D symmetric Gaussian point-spread function (PSF) to the raw pixel intensities within the 5 × 5 pixel box, using a camera-noise model that accounts for baseline (detector dark current), sensitivity, gain, and quantum efficiency. The MLE approach maximizes the likelihood of observing the measured pixel values given the PSF model plus Poisson-distributed shot noise and Gaussian readout noise[12]. The fit yields a sub-pixel (x, y) centroid position, a photon count, a PSF width σ, and a background estimate for every detected molecule in every frame. Localization files are stored in HDF5 format (one file per ND2 movie) with an accompanying YAML metadata file, following the Picasso file convention. Protein and vesicle channel files are processed independently with their respective gradient thresholds (both from the optimization in Step 1).

*Step 3. ROI definition, trace extraction, and colocalization labeling*

Each Picasso HDF5 file contains one independent localization per detected molecule per frame. To identify unique molecules, localizations in successive frames are linked using a nearest-neighbor algorithm (Picasso's `postprocess.link`, based on the Crocker–Grier algorithm[13]): a localization in frame *t* is assigned to the same molecule as the nearest localization in frame *t*-1, provided the displacement is below `max_distance` (default 2 pixels). A molecule is allowed to disappear for up to `max_darktime` frames (default 4,000) before its track is terminated. This accommodates physically plausible dark states without fragmenting a single molecule's history into multiple tracks. The linked result yields one entry per unique molecule, with sub-pixel centroid coordinates averaged over all detected frames.

Lipid-dye-labeled vesicles emit continuously and are therefore detected in virtually every collected frame. The resulting localizations are collapsed into single vesicle positions by density-based clustering (Picasso's `clusterer.cluster`). Localizations within `max_distance_ground_truth` (default 2.5 pixels) of each other are grouped, requiring a minimum of `min_on_ground_truth` (default 3) localizations per cluster. Cluster centers are then computed as the intensity-weighted centroid over all member localizations. This procedure is equivalent to Density-Based Spatial Clustering of Applications with Noise (DBSCAN)[9] applied to spatial coordinates and is robust to the variable localization density across vesicles of different brightness.

For each linked protein localization, a 5 × 5 pixel ROI box is placed at the rounded sub-pixel centroid. The pipeline detects spatial overlaps by sequential mask painting: each box is rendered onto an integer label image, and the existing labels beneath a newly placed box are inspected. If any previously placed ROI contributes more than `overlap_threshold` (default 3) overlapping pixels, both ROIs are flagged as overlapping. Overlapping ROIs are extracted and stored in the trace DataFrame with overlap = "overlapping" but are excluded from all downstream analysis, because the summed intensity in an overlapping box contains contributions from multiple molecules and cannot be interpreted as a single-molecule signal. Only single (non-overlapping) ROIs are used for training.

For each single ROI, the intensity trace is computed as the pixel-sum over the 5 × 5 box at every frame:

$$I_{\text{trace}}(t) = \sum_{i,j \in \text{box}} \text{movie}(t, i, j)$$

This integrated box intensity captures the total fluorescence from the molecule including photon shot noise. The trace is 6,000 values long, one per frame. Each protein trace is assigned a ground-truth label based on its proximity to the nearest vesicle cluster center, using a KD-tree nearest-neighbor query[6]. Traces whose centroid falls within `max_dist_closest_ground_truth` (default 4 pixels) of a vesicle center are labeled IN, representing proteins that were successfully encapsulated inside a vesicle. All other single-ROI traces are labeled OUT and are assumed to originate from random noise or protein molecules that escaped their vesicle and remained adsorbed on the glass surface. The IN label is the positive class for machine-learning classification.

In addition to protein traces, the pipeline samples `n_background_traces` (default 50) background traces per movie from randomly selected pixels that do not overlap with any protein ROI. These traces capture the camera offset, read noise, and any diffuse autofluorescence or out-of-focus protein, providing a noise reference for quality control and classifier calibration. A spatial buffer of `background_buffer` pixels (default 5) around all protein ROIs is enforced to avoid partial contamination.

*Step 4. Trace aggregation, background correction, and normalization*

Per-movie trace pickles from all experiments are concatenated by protein identity. Each trace is assigned a globally unique integer ID (uniqueID) that traces back to its experiment and original movie for subsequent audit. Overlapping traces are discarded at this stage; only IN-single and OUT-single traces are retained. Traces are stored in matrix format (rows = frames, columns = traces).

The raw trace for each molecule contains a constant camera offset plus any diffuse (unfocused) fluorescence background. These contributions are estimated and removed by fitting a two-component Gaussian Mixture Model (GMM) independently to the raw intensity values of each trace:

$$p(I) = \pi_1 \mathcal{N}(I;\ \mu_1, \sigma_1^2) + \pi_2 \mathcal{N}(I;\ \mu_2, \sigma_2^2)$$

where the lower-mean component ($\mu_1$, $\pi_1$) represents frames in which the molecule is in the off (dark) state or permanently bleached, and the higher-mean component ($\mu_2$, $\pi_2$) represents frames in the on (emitting) state. The background estimate is the mean of the lower component, $\hat{\mu}_{\text{bg}} = \mu_1$. The background-removed (bg_rm) trace is:

$$I_{\text{bg\_rm}}(t) = I_{\text{raw}}(t) - \hat{\mu}_{\text{bg}}$$

GMM fitting is performed with five random initializations to avoid local minima, with a fallback to the mean of the last 500 frames for traces where fitting fails.

Two normalized versions of each background-removed trace are computed for downstream machine learning: 1) Z-score normalization: subtract the trace mean and divide by the trace standard deviation (`sklearn.preprocessing.StandardScaler`). This centers each trace at zero and scales it to unit variance, making classifier training insensitive to absolute brightness; and 2) Minmax normalization: linearly rescale each trace to the [0, 1] interval (`sklearn.preprocessing.MinMaxScaler` applied to the bg_rm trace). This preserves the relative step heights between on and off events. Both normalizations are computed per trace.

*Step 5. Quality filtering*

Traces are filtered using a cascade of five independent criteria applied to the background-corrected trace. The criteria are designed to retain traces that exhibit genuine, repeatable on/off switching behavior while removing camera noise, adsorbed aggregates, and imaging artifacts. All five criteria must be satisfied; any failure causes the trace to be discarded.

A GMM-based signal-to-noise (SNR) gate is defined by fitting a two-component GMM to each background-removed trace (same model as step 4), and the separation between components is computed:

$$\mathrm{SNR} = \frac{\mu_{\mathrm{signal}} - \mu_{\mathrm{noise}}}{\sigma_{\mathrm{noise}}}$$

Where $\mu_{noise}$ and $\sigma_{noise}$ are the mean and standard deviation of the lower GMM component, and $\mu_{signal}$ is the mean of the upper component. Traces with SNR < `snr_min_separation` (default 2.0) are rejected as pure noise: the two components overlap too heavily to distinguish a real signal from fluctuating background. This gate precedes all peak-based criteria because fitting a noise trace with a GMM always produces two artificial components with low separation regardless of their actual information content.

The fraction of frames classified as on (signal state) by the GMM is computed for each trace. Traces where this fraction exceeds `max_duty_cycle` (default 0.3) are rejected because for HMSiR, this would be an unnaturally large duty cycle and we observed that most traces with such large duty cycles did not show blinking behavior but were instead large drifts in baseline during acquisition.

Each frame is classified into one of two states using the posterior probability of the GMM signal component:

$$\mathrm{state}(t) = \begin{cases} \text{noise} & P(\text{signal} \mid I_t) < \theta \\ \text{signal} & P(\text{signal} \mid I_t) \geq \theta \end{cases}$$

where $\theta$ = `gmm_proba_threshold` (default 0.9) is the posterior probability threshold. Contiguous runs of signal frames constitute emission peaks. Peaks shorter than `min_peak_width` (default 1) frames are discarded as noise spikes. Traces with `n_peaks` ≤ `min_peak_number` (default 3) are rejected. This ensures that only traces displaying a few distinct on–off emission cycles are retained. Multiple cycles provide a statistically meaningful fingerprint of the photophysical environment, which is the quantity the machine learning model is trained to classify.

Three additional criteria on peak timing eliminate traces with artifactual temporal patterns. The first peak must appear before frame `first_peak_time` (default 1,000), ensuring the molecule was present at the start of acquisition rather than adsorbing during imaging. The last peak must appear after frame `last_peak_time` (default 100), confirming activity extends beyond the very first frames. The gap between the first and second peaks must be shorter than `delta_first_second` (default 1,000 frames).

Together, the five criteria impose the expected temporal signature of a stochastically blinking single molecule that is present throughout the entire 3-minute acquisition: early onset, repeated bursts, not too many frames above GMM threshold (artifactual step-on), and high statistical confidence that the on-state signal is genuinely distinct from noise. Examples of accepted and rejected traces are displayed in Figure S8.

**Machine Learning**

*General workflow*

The entire machine learning code was implemented in PyTorch. For classification, we used only accepted traces IN vesicles. In a first phase we performed 4-fold cross-validation for model architecture comparison and data-augmentation tuning, running on shorter traces with Monte Carlo (MC) Dropout uncertainty quantification and a Wasserstein distance (WD) sweep to calibrate a confidence filter. In the second phase, we trained a final model on the full-length traces with a single stratified split and MC Dropout evaluation was linked to source trace identifiers. The two phases share the same training loop, optimizer, loss function, augmentation strategy, early stopping logic, and MC Dropout infrastructure.

*Trace loading and pre-processing*

Single-molecule traces are loaded from filtered pickle trace matrices. Within each pickle file the trace matrix has shape ($T$, $M$), frames × traces, and is loaded as a set of $M$ individual time series. Each pickle file also stores the two pre-computed normalizations (minmax, zscored) as separate columns. The channels list in the config (["minmax", "zscored"]) selects which channels to load and stack along the channel axis, producing the $(N, 2, T)$ input tensor.

The function assembles the full dataset array $X$ of shape $(N, C, T)$ and a label vector $y \in \{0, \dots, K-1\}^N$, and assigns a globally unique integer identifier (uniqueID) to every trace. The uniqueID traces back to the source experiment and movie for later audit.

Two optional preprocessing operations are applied before training: Trace trimming (`trim_end`) removes the last `trim_end` frames from every trace. For K-fold cross-validation, this is set to 2,000 frames (reducing 6,000-frame traces to 4,000 frames), which substantially decreases training time without discarding the most informative early blinking events. For final model training we used `trim_end` = 0 and to train on the full 6,000-frame traces. Optionally, fewer than the total amount of available traces can be used for cross-validation or training by using the `max_traces_per_class` option. This is useful for debugging runs, but for this publication all traces available were used.

*Class balancing*

The number of traces per protein can differ substantially across experimental conditions. Two complementary strategies ensure the classifier does not simply learn to predict the most frequent class. For the training set, we used `WeightedRandomSampler`. `DataLoader` samples training indices with replacement, where the sampling probability of sample $i$ with class $c_i$ is proportional to $w_{c_i} = 1/n_{c_i}$ (inverse class frequency). Each epoch draws exactly $N_{\text{train}}$ indices, so the effective class distribution seen by the model is uniform regardless of the raw imbalance. This avoids discarding any majority-class traces: all traces contribute to training, just not at equal frequency. For the validation and test sets, we performed majority-class subsampling: `balance_val = balance_test = True`, so the majority class is randomly downsampled to match the minority class size before evaluation.

*Data augmentation*

Augmentation is applied only to the training set. The augmented dataset is constructed by repeating each real trace `aug_factor` times with independently sampled transformations, so the effective training set size grows by a factor of (`aug_factor + 1`). Four operations are available:

Time warping: the time axis is deformed by a smooth random displacement field. A displacement signal $\delta(t)$ is drawn from a zero-mean Gaussian process with standard deviation `time_warp_sigma` (default: 0.5), smoothed by a Gaussian kernel to ensure continuity, and integrated to produce a monotonically increasing warped time grid $\tilde{t}$. The trace is then resampled onto the original time grid by cubic spline interpolation. This simulates realistic variability in blinking kinetics without introducing artificial discontinuities.

Gaussian noise: independent Gaussian noise is added to every frame:

$$\tilde{I}(t) = I(t) + \varepsilon(t), \varepsilon(t) \sim \mathcal{N}(0, \sigma_{\text{noise}}^2)$$

with `noise_sigma` (default: 0.5) in the units of the normalized trace. This simulates photon shot noise variability and prevents the classifier from overfitting to specific noise realizations.

Magnitude jitter: a multiplicative scaling factor is drawn uniformly:

$$\tilde{I}(t) = I(t) \cdot (1 + u), u \sim \text{Uniform}(-\delta, +\delta)$$

with `magnitude_jitter` $= \delta$ (default: 0.5). This simulates run-to-run variability in excitation intensity and dye brightness, making the classifier invariant to small global amplitude rescalings.

Time reversal (mirroring): the trace is reflected in time, $\tilde{I}(t) = I(T - 1 - t)$. This is a valid augmentation because the photophysical blinking statistics of a single fluorophore are approximately time-reversible. Only the fraction of the trace between the first and last detected peaks is time-reversed, so that time reversal does not change the timings of first and last blinking events. For cross-validation, distinct augmentation seeds (random_seed = seed + fold) are used, so the synthetic samples differ between folds. An example of a portion of a single-molecule trace augmented in these four ways is displayed in Figure S9.

*Model zoo*

Three models were explored (CNN-GRU, ResNet, TCN) and they all accept tensors of shape $(N, C, T)$ and produce logit vectors of length $K$ (number of classes).

CNN-GRU (`orig_conv_gru`): A two-stage encoder that first applies a stack of 1D convolutional layers (with batch normalization and ReLU activations) to extract local temporal features at multiple scales, then passes the resulting feature sequence into a gated recurrent unit (GRU) to integrate information across the full trace length. The final hidden state of the GRU is projected to class logits by a linear head. The convolutional stage provides translation invariance within local windows; the GRU captures long-range temporal dependencies such as the timing and ordering of bleaching steps. This model is a PyTorch implementation of our previously published (Keras-based) model[14].

1D ResNet (`resnet1d`): a stack of one-dimensional residual blocks. Each block applies two 1D convolutions (kernel size 3, same padding) with batch normalization and ReLU and adds the input via a skip connection (with a 1×1 projection if channel dimensions change). Global average pooling collapses the time dimension after the last block, and a linear layer maps the pooled features to logits. Residual connections enable gradient flow through deep networks, allowing more layers to be trained without degradation.

TCN (tcn): a stack of dilated causal 1D convolutions with exponentially increasing dilation rates $d_k = 2^k$. Each layer at depth $k$ has a receptive field that extends $2^k$ steps into the past, so the total receptive field grows exponentially with depth, covering the entire trace length with far fewer parameters than a fully convolutional network. Residual connections are applied with 1×1 projection if the number of channels changes between blocks.

All three models include dropout layers (rate dropout, default 0.2) after the non-linearities. Dropout serves two roles: regularization during training, and stochastic sampling for MC Dropout uncertainty quantification at inference time (see below).

*Optimizer and loss function*

`AdamW` optimizer: parameters are updated by the Adam rule with decoupled weight decay[15]:

$$\theta_{t+1} = \theta_t - \alpha \frac{\widehat{m}_t}{\sqrt{\hat{v}_t} + \epsilon} - \alpha\lambda\theta_t$$

where $\widehat{m}_t$ and $\hat{v}_t$ are bias-corrected first and second moment estimates, $\alpha$ is the learning rate ($4 \times 10^{-4}$), and $\lambda$ is the weight decay coefficient ($4 \times 10^{-4}$). Decoupled weight decay regularizes all parameters uniformly without interfering with the adaptive learning rate.

`CrossEntropyLoss` with label smoothing: the training objective is

$$\mathcal{L} = -\sum_{c=1}^{K} \tilde{y}_c \log\, p_c$$

where $p_c = \text{softmax}(\text{logits})_c$ and the smoothed target is

$$\tilde{y}_c = (1-\varepsilon)\cdot \mathbb{1}[c=y] + \frac{\varepsilon}{K}$$

with smoothing factor $\varepsilon = 0.001$. Label smoothing prevents the model from becoming overconfident on the training set by distributing a small fraction of the target probability mass uniformly across all classes.

Gradient clipping: after each backward pass, the global $\ell_2$ norm of all parameter gradients is clipped to 1.0 (`torch.nn.utils.clip_grad_norm_`). Clipping prevents exploding gradients, which are particularly common in recurrent architectures (CNN-GRU) and in networks with very large receptive fields (TCN).

*Combined early stopping*

The model checkpoint condition is a conjunction of two criteria evaluated at the end of each epoch:

Primary condition: Area under the curve (AUC) improvement: $\text{AUC}_{\text{val}} \geq \text{AUC}_{\text{best}} + \delta_{\text{AUC}}$ with $\delta_{\text{AUC}} = 0.005$, and loss constraint: $\mathcal{L}_{\text{val}} \leq \mathcal{L}_{\text{best}} \cdot 1.10$.

Both criteria must be satisfied simultaneously. The AUC criterion prevents checkpointing when the model has only marginal improvements in AUC. The loss constraint prevents accepting a checkpoint where the validation loss has risen substantially above its minimum, even if the AUC improved slightly.

Alternative condition: a checkpoint is also saved when the AUC is within 0.003 of the best AUC and the validation loss has decreased by at least 0.005 below the loss at the last checkpoint. This allows the optimizer to continue refining the model after the AUC plateaus, as long as the loss is still improving substantially. If neither condition is met, a patience counter is incremented. Training terminates when the counter reaches `patience_limit` (10 epochs without a checkpoint). The maximum number of epochs is 500 as default. The best model weights are saved as a CPU-side copy of the state dictionary at each checkpoint, so the best-ever checkpoint is available regardless of when training stops.

*Monte Carlo Dropout uncertainty quantification*

Uncertainty quantification was implemented using Monte Carlo (MC) Dropout at inference time analogously to our previous work[14]. MC Dropout[16] repurposes the dropout layers to generate a distribution over predictions for each input. The model is set to training mode (`model.train()`) to activate dropout, and $n_{\text{MC}} = 100$ independent forward passes are run for the same input batch.

Each pass samples a different dropout mask, producing a slightly different output probability vector. The result is a three-dimensional array of shape $(n_{\text{MC}}, N, K)$.

The mean probability vector $\bar{p}_i = \frac{1}{n_{\text{MC}}} \sum_{m=1}^{n_{\text{MC}}} p_i^{(m)}$ is used for the final class prediction ($\hat{y}_i = \arg\max_c \bar{p}_i^c$). The spread of the individual-pass probabilities across the $n_{\text{MC}}$ samples characterizes the model's epistemic uncertainty for each trace. Traces for which the dropout masks consistently agree produce tight, well-separated class-probability distributions, while uncertain traces produce diffuse, overlapping distributions.

The per-sample uncertainty is quantified by the Wasserstein distance (WD) between the MC Dropout probability distribution for the predicted class and the distribution for the nearest competing class:

$$W_i = \min_{c \neq \hat{y}_i} W_1 \left( \{p_i^{(m),\hat{y}_i}\}_{m=1}^{n_{\text{MC}}}, \{p_i^{(m),c}\}_{m=1}^{n_{\text{MC}}} \right)$$

where $W_1$ is the one-dimensional Wasserstein-1 distance (Earth Mover's Distance), computed via quantile interpolation on a common grid of $n_{\text{MC}}$ mid-quantile points:

$$W_1(X, Y) = \frac{1}{m} \sum_{k=1}^{m} \left| Q_X \left( \frac{k - 0.5}{m} \right) - Q_Y \left( \frac{k - 0.5}{m} \right) \right|$$

A large $W_i$ means the predicted-class probability mass is well-separated from all competitor-class probability masses across MC samples. In such case, the model is confidently and consistently predicting the same protein identity. A small $W_i$ means the MC samples fluctuate between competing proteins. In this case, the model is uncertain, typically because the trace is ambiguous or poorly representative of the training distribution.

For all test sets, we report a WD sweep: 50 linearly spaced thresholds $\tau$ from 0 to $\max_i W_i$ are evaluated. At each $\tau$, only the subset of samples with $W_i > \tau$ is retained, and the balanced accuracy on this subset is recorded alongside the fraction of retained traces. This produces a trade-off curve: as $\tau$ increases, fewer but more confident traces are retained, typically with increasing accuracy.

Threshold selection: for cross-validation runs, a fixed `wasserstein_threshold` = 0.7 was applied (bypassing the auto-sweep), chosen to correspond to a well-separated region of the WD distribution based on preliminary runs. In full-length trace model training a default auto-selection is used: the threshold is chosen to maximize balanced accuracy subject to removing no more than 50% of samples.

*Cross-validation for model selection*

We used stratified K-fold cross validation to evaluate and compare multiple model architectures and augmentation factors. Because K-fold cross validation re-trains the model K times per combination, it provides a robust estimate of generalization performance with confidence intervals, at the cost of K times the compute.

For each protein comparison, three sets of cross validation runs were performed using traces with time reversal (mirrored), without time reversal, and a background control to verify that classification performance degrades to chance when only background noise is used for training and classification. Each set was run for all three protein pairs (Halo-D106 vs SNAP-C148, Halo-D106 vs Halo-K117, and scGrx1 vs scGrx1-AcK20) and for all three normalization strategies (both channels, minmax only, zscored only).

*K-fold setup*

`StratifiedKFold(n_splits=4, shuffle=True, random_state=840410)` partitions the dataset into 4 folds, maintaining the protein-class ratio in each fold to within one sample. For each fold $k \in \{1, \ldots, 4\}$ there is a held-out fold (25% of data), which serves as evaluation set for this fold. The remaining 75% of the data serve as training set, to which augmentation is applied. Then, a fresh model is instantiated (weights re-initialized), trained from scratch, and evaluated on the validation fold. MC Dropout is run on the validation set and the WD threshold is applied.

At the end of all 4 folds, the confusion matrices are aggregated and mean and standard deviation of the K-folds are reported.

*Augmentation sweep*

To test the effect of data augmentation on training, a list of augmentation factors is provided (factors: [0, 3, 5]). The Cartesian product of models × augmentation factors defines a set of tasks, each being a full 4-fold cross validation run for one (`model, aug_factor`) combination. The aggregate results (AUC mean ± std across folds, balanced accuracy mean ± std, training time per epoch) are collected and visualized as a heatmap with models on the rows and augmentation factors on the columns. This plot directly answers which architecture benefits most from augmentation and which augmentation factor produces the best performance/cost trade-off.

Cross-validation and augmentation sweeps are also applied to the training of background noise using traces from the experiments of each protein. These experiments help identify which combinations of models and augmentation strategies learn the most from background noise. The selection of models for final training takes into account the best performance for true protein signals paired with the least learning of the corresponding background traces.

*Final training*

The final model and classified traces are obtained by training on the full length of all available traces. The dataset is partitioned into three non-overlapping sets using two successive stratified random splits:

$$\text{train} + \text{temp} \leftarrow \text{train\_test\_split}(p_{\text{temp}} = 0.30, \text{stratify} = y, \text{seed})$$

$$\text{val} + \text{test} \leftarrow \text{train\_test\_split}(\text{temp}, p_{\text{test}} = 0.50, \text{stratify} = y_{\text{temp}}, \text{seed})$$

The result is a 70 / 15 / 15 split. `stratify=y` ensures the protein-class ratio is preserved in each set. The model is trained on the training set, early stopping is evaluated on the validation set, and the test set is used only for the final evaluation.

*MC Dropout with trace-level linkage*

`evaluate_uncertainty_filtered` receives two additional arguments in final training compared to cross validation: 1) `traces`: the raw test set intensity traces (N, C, T), for saving alongside WD scores; and 2) `unique_ids`: per-trace integer identifiers. The `unique_ids` are assigned during dataset loading and are threaded directly through the stratified split: `create_dataloaders` performs all splits by integer index (`np.arange(len(X))`), applies `train_test_split` to those indices, and indexes into `unique_ids` with the resulting test-set index array. Any subsequent majority-class subsampling (`balance_test`) is applied consistently to both the data and the `unique_ids` array using the same subsample indices. This guarantees an unambiguous one-to-one correspondence between every test trace and its uniqueID. The uniqueID linkage makes it possible to retrieve the original ND2 movie, frame range, and localization position for any trace that the MC Dropout filter flags as uncertain, enabling manual verification of borderline predictions directly in the raw microscopy data, as well as downstream extraction of interpretable features.

*Extraction of interpretable features*

After final training, traces are filtered by classification uncertainty (Wasserstein distance, `mcd_filter.wasserstein_min`) and all passing traces are analyzed together. Peaks are selected by running GMM on channel 0 (minmax-normalized) using the threshold `gmm_proba_threshold`. Peak intensities reported in intermediate outputs use channel 1 (z-scored). The `min_peak_width` parameter sets the minimum number of consecutive "on" frames required for a run to be counted as a peak (identical to the Extraction pipeline setting). An active window is defined as the span from the first frame of the first peak to the last frame of the last peak. Three features are extracted per trace and arranged in a 1×3 panel: mean off-time (mean dark-interval duration per trace), mean on-time (mean peak duration per trace), and duty cycle (on-frames/active-window frames per trace).

For binary comparisons (exactly two proteins), Mann-Whitney U tests are run on each of the three features and p-values are corrected for multiple testing using Benjamini-Hochberg false-discovery rate correction. Corrected p-values and effect sizes (rank-biserial correlation coefficients r) are displayed as significance brackets on the violin plots.

### Further control experiments for the TCN classifier of scGrx1 acetylation

*Label scrambling*

This experiment involved the training of traces with scrambled protein labels. This was implemented in the following way: immediately after the full dataset ($X$, $y$) is assembled by `build_dataset_from_keys`, the label vector $y$ is shuffled in place with `np.random.default_rng(seed).shuffle(y)` before the train/val/test split. This ensures all three splits receive randomly relabelled traces, so no label information can be exploited. Everything else was kept identical to the main training pipeline: TCN architecture, mirroring augmentation (`aug_factor=1`, `include_mirror=True`, no noise or warp), AdamW optimization, MC Dropout (100 passes).

*Noise classification with pre-trained model*

To test whether during training on real protein signals the models learns features from the background noise, we used the pre-trained TCN model of the scGrx1 vs AcK20-scGrx1 to classify background traces. The background traces were given the label of the protein that was present in the same FOV. This was an inference-only run, no training occurs, and class probabilities were obtained from a single deterministic forward pass.

#### Random forest classification with interpretable features

A random forest (RF) trained on six hand-engineered blinking statistics tests how much discriminative information is already captured by simple summary statistics, without any learned representation by a deep learning model (e. g., TCN). This serves two purposes: 1) Classical machine learning baseline: if the RF achieves comparable performance, the TCN's added complexity may not be necessary. If the TCN substantially outperforms the RF, it implies the raw time-series contains information that summary statistics cannot capture; and 2) Internal consistency: the RF pipeline includes its own label-scrambling control, providing a reference point for that validation at the feature level.

Traces were loaded from the minmax channel (single channel, same traces used in training). For each trace, a 2-component GMM classifies every frame as ON or OFF using a posterior probability threshold of 0.9 and a minimum peak width of 1 frame (30 ms per frame). Traces with fewer than 2 detected ON events are discarded. Six features are computed per trace: 1) mean duration of ON-state dwell times (ms); 2) mean duration of OFF-state dwell times (ms); 3) standard deviation of ON dwell times (ms); 4) standard deviation of OFF dwell times (ms); 5) sum of all ON dwell times in the trace (ms); and 6) sum of all OFF dwell times in the trace (ms).

The pipeline runs first a classifier using a stratified 80/20 train/test split with both halves subsampled to the minority class. A RF classifier with 500 trees is trained and, to simulate MC Dropout, a vote-margin filter ($|p(class0) - p(class1)|$) sweeps 50 thresholds to find the value that maximizes balanced accuracy subject to $\leq$50% trace removal. Outputs include a normalized confusion matrix, a feature importance bar chart, and a margin sweep scatter plot.

After this real classification run, a label scrambling control is run: The full label vector is globally shuffled before the train/test split, making class information unavailable to the RF. From there, the training procedure is identical to the one described above.

#### Pose modeling

To determine the possible poses that the covalently bound HMSiR could adopt in all proteins and conjugation sites, we employed Boltz-2[17] to generate large structural ensembles (500 poses per condition) of covalent protein–ligand complexes. Four protein contexts (Halo-D106, Halo-K117, scGrx1, and AcK20-scGrx1) were modeled across three ligand forms (closed, open deprotonated, open protonated) each, totaling 12 independent ensemble runs. All poses were subsequently clustered by ligand root-mean square deviation (RMSD), and PyMOL sessions were generated for the top clusters. Boltz-2 was run using the BioPipelines platform[18].

*Pose analysis*

For every predicted model, the distance between the modified protein atom and ligand atom was measured. Poses where this distance exceeded `cov_bond_cutoff` (default 3.0 Å) were discarded as malformed predictions in which the covalent constraint was not satisfied. All retained poses were superimposed onto the first valid pose (model 0) by aligning protein Cα atoms using MDAnalysis alignto. This removed rigid-body rotation and translation, leaving only differences in ligand pose and protein conformation. The all-heavy-atom RMSD matrix was computed across the aligned ligand coordinates (hydrogen atoms excluded). RMSD was normalized by the square root of the number of ligand heavy atoms: $RMSD(i,j) = || coords_i - coords_j || F / (n_{atoms})^2$ to give a symmetric N × N distance matrix where N is the number of retained poses.

The condensed RMSD matrix was clustered using average linkage (`scipy.cluster.hierarchy.linkage, method=average`). The dendrogram was cut at `rmsd_threshold` (5.0 Å), producing flat cluster labels. Average linkage was chosen over Ward linkage because the cut threshold maps directly to a physically interpretable quantity: two clusters merge when their average pairwise cross-cluster RMSD falls below the threshold.

For each cluster, the medoid (most central pose) was identified as the pose with the minimum mean intra-cluster RMSD to all other members of the same cluster. The medoid was used as the structural representative of the cluster for display in Figures S18, S18, S27 and S28. Clusters were sorted by population (number of poses) in descending order. The top `top_n` clusters (default 10) were selected for further analysis.

### Supporting Figures

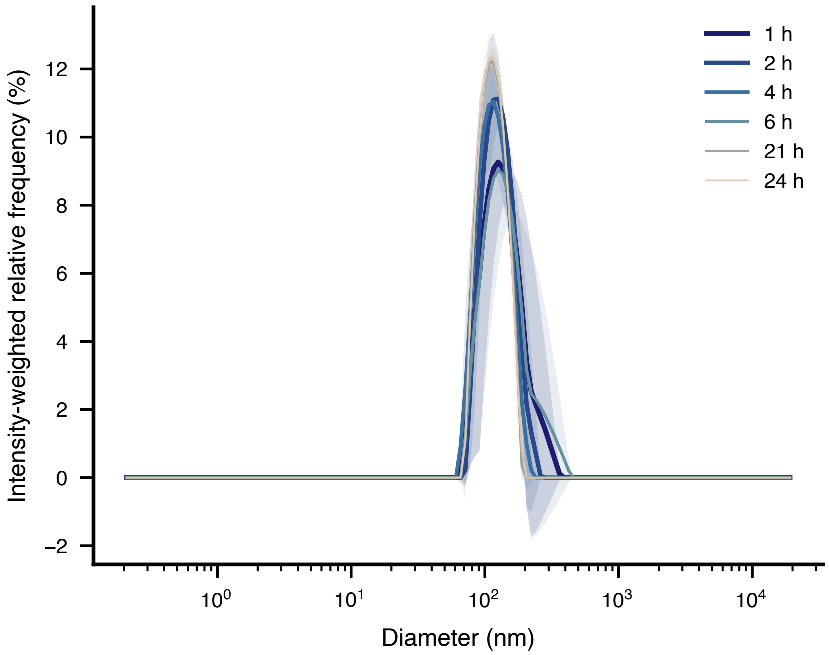

**Figure S1**. Distribution of liposomes sizes measured by DLS at varying time points after preparation. Solid lines are means and shaded areas represent standard deviation of three preparative replicates.

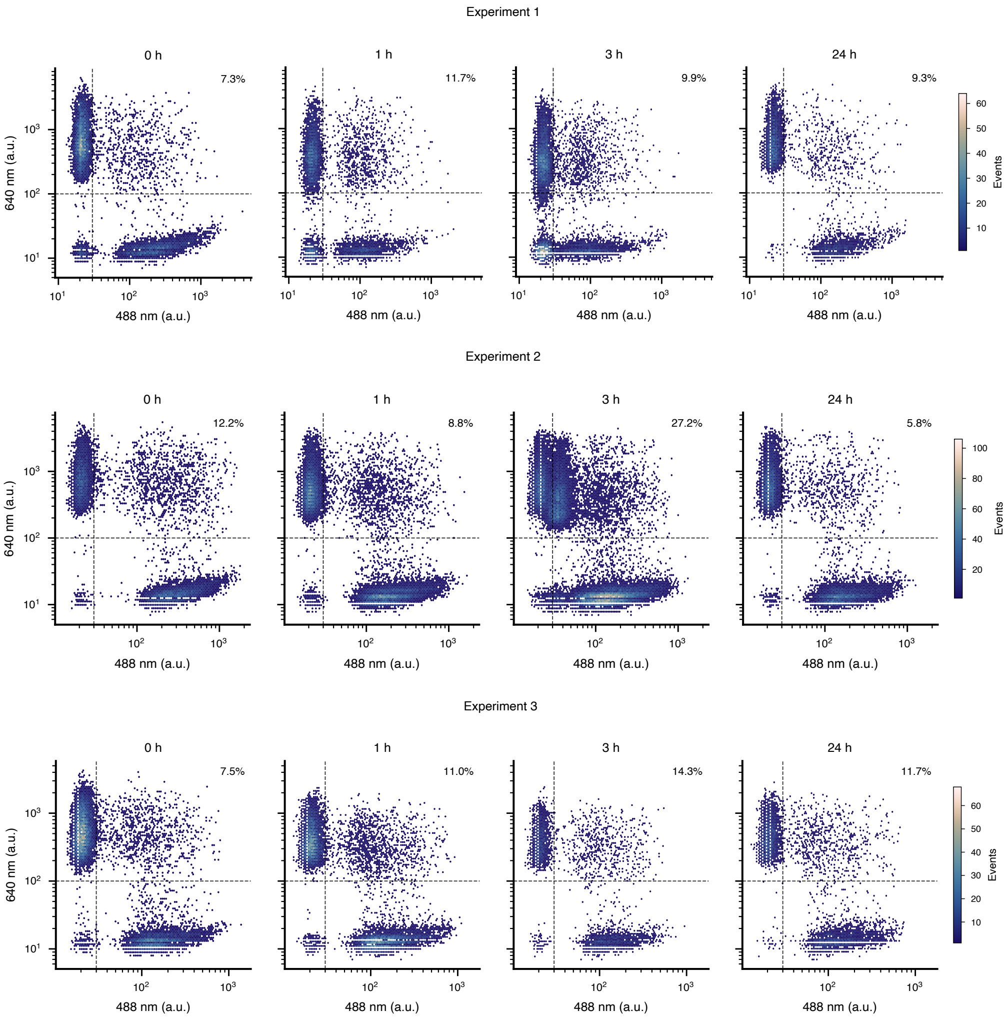

**Figure S2**. NanoFCM fluorescence scatter of mixed DPPE-Atto520 and DPPE-Atto647N-labeled vesicle populations over time. Three independent preparative replicates were measured by nano-flow cytometry (NanoFCM). Each panel shows a hexbin density scatter plot of the 488 nm (FITC-H, Atto520) versus 640 nm (PC5-H, Atto647N) fluorescence signals, both axes on a log scale. Color encodes local event density on a shared scale per replicate (lapaz colormap, low to high). Before plotting, events were gated on the side-scatter channel (SS-H) by retaining those between the 5th and 99.5th percentile of each acquisition, to select the vesicle population and exclude instrument noise and aggregates. The number of gated events per panel ranged from 3,204 to 8,081. Where multiple acquisitions were available for a given timepoint, the file acquired at a sample concentration of approximately $2\times10^3$ particles $\mu L^{-1}$ was preferred. Vesicles carrying only DPPE-Atto520 appear along the horizontal axis; vesicles carrying only DPPE-Atto647N appear along the vertical axis; double-positive events in the upper-right region indicate membrane mixing between the two populations.

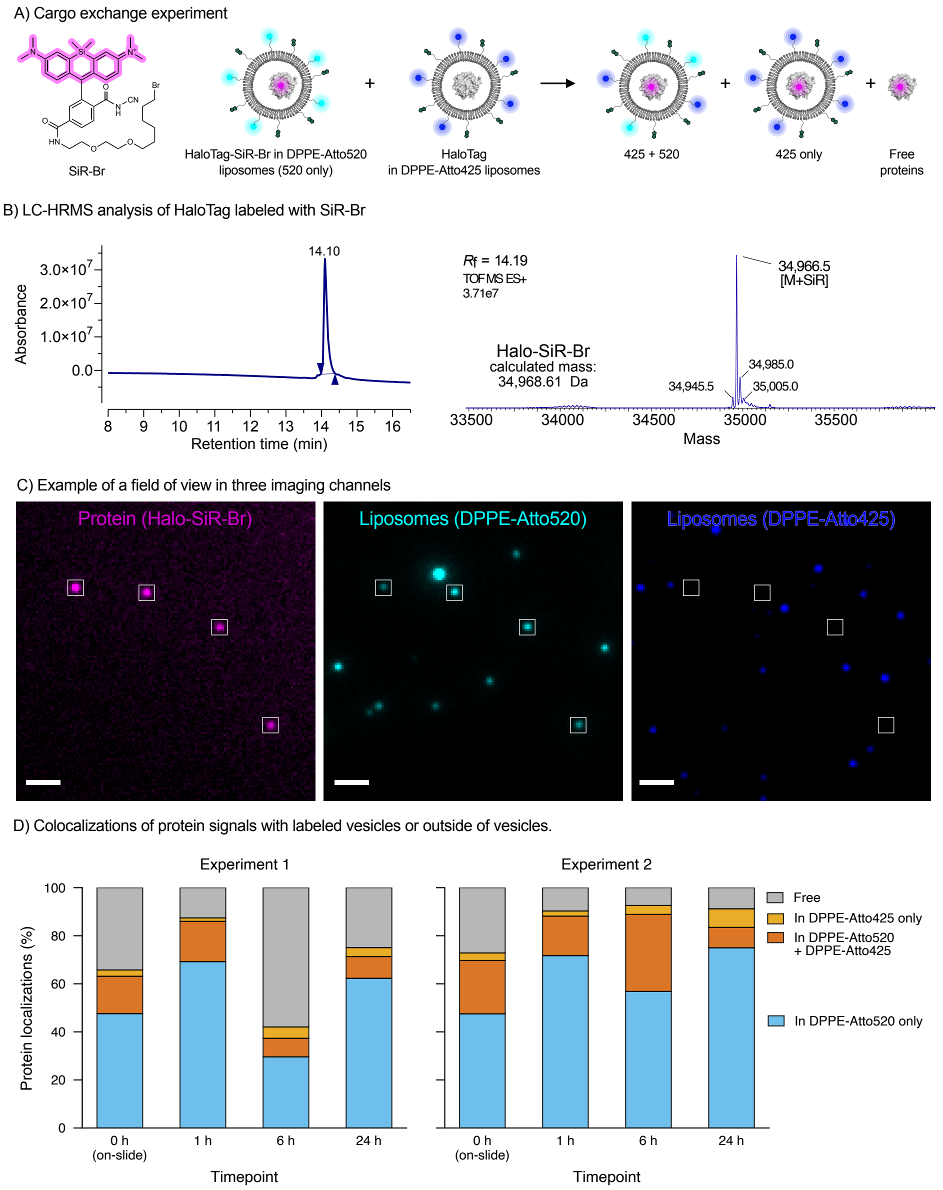

**Figure S3**. Cargo exchange between pre-formed vesicles. A) Scheme of potential exchange of carbo and lipids between pre-formed vesicles. B) LC-HRMS analysis of HaloTag labeled with SiR-Br. C) Example field of view. Left to right: 640 nm channel (magenta, protein HaloTag labeled with SiR-Br), 515 nm channel (cyan, DPPE-Atto520-labeled vesicles), 405 nm channel (blue, DPPE-Atto425-labeled vesicles). White squares indicate protein localizations detected by Picasso[10]. Scale bar = 2 μM. D) Quantification of protein localization across timepoints for two independent preparative replicates. At each independent timepoint (0 h–mixing on-slide,1 h, 6 h, or 24 h), protein localizations from 80 fields of view per slide were classified by their nearest-neighbor colocalization with either vesicle population (radius threshold: 3 pixels).

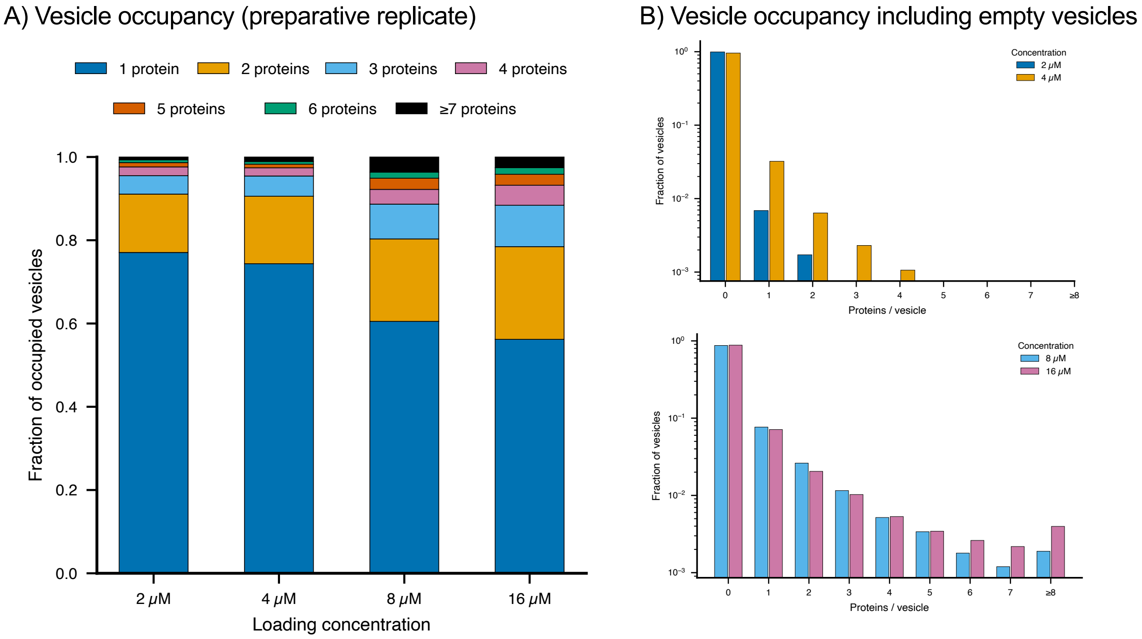

**Figure S4**. Vesicle protein occupancy distributions. A) Stacked bar chart for the second independent preparative replicate, in the same format as Figure 2C. Bars show the fraction of occupied vesicles (those carrying at least one detectable protein) with 1, 2, 3, 4, 5, 6, or ≥7 proteins, as determined by Kalafut-Visscher step detection applied to TIRF photobleaching traces. Data shown for four protein loading concentrations (2, 4, 8, and 16 μM Halo-SiR-Br). B) Full occupancy distributions at low (2 and 4 μM) and high (8 and 16 μM) loading concentrations. Bars show the fraction of all vesicles, including empty ones, with a given number of proteins per vesicle (log scale y-axis). The total vesicle count per condition was obtained by Picasso[10] cluster detection on the 488 nm channel. Empty vesicles (0 proteins) represent the dominant population at all concentrations.

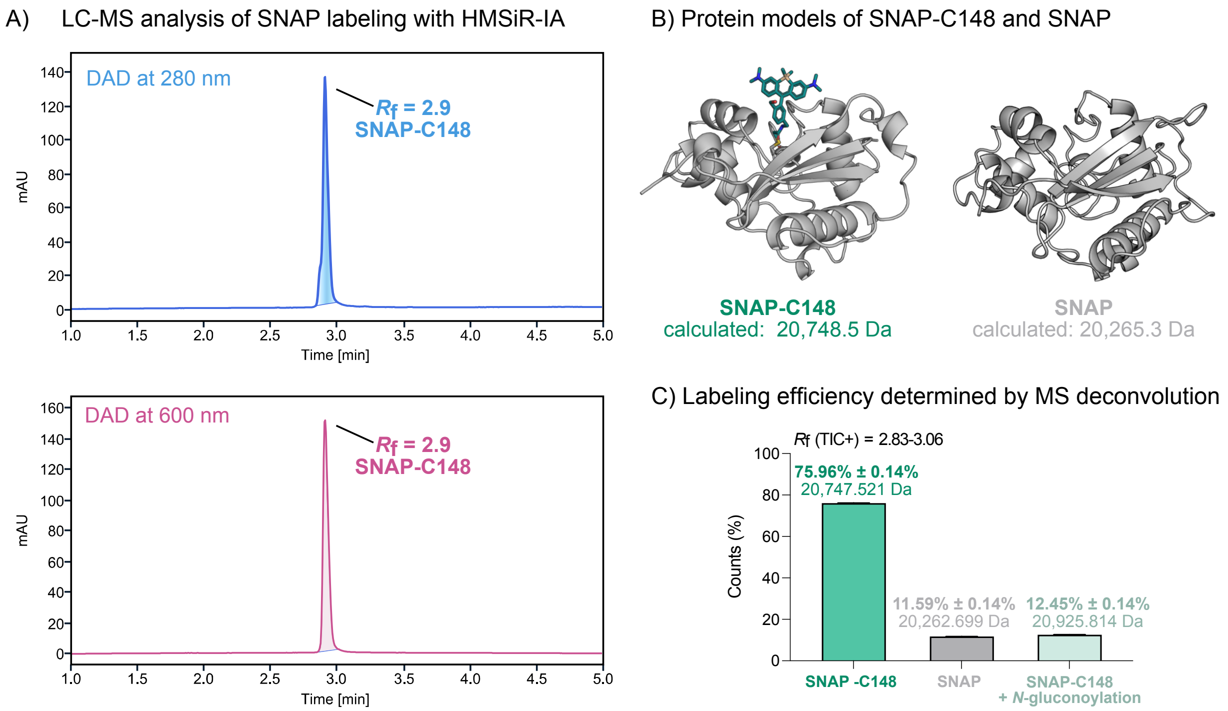

**Figure S5**. LC-MS analysis of SNAP-C148 labeled with HMSiR-IA. A) Integrated protein peak ($R_f$ = 2.9) in the diode array detector (DAD) chromatogram at 280 nm (top) and 600 nm (bottom). B) Protein structure models with corresponding calculated mass for SNAP-C148 (20,748.5 Da) and unlabeled SNAP$_f$ (20,265.3 Da). Models were generated with Boltz-2[17]. C) Protein masses and labeling efficiency were determined by deconvolution of the TIC+ peak ($R_f$ = 2.83-3.06) corresponding to both the labeled and unlabeled protein peaks with SAMMI (Cerno Biosciences)[4,5]. The labeling reaction was performed with 20 μM protein and 5 equiv. HMSiR-IA (2 h at 37 °C). The reaction was purified by centrifugal membrane filters. The LC-MS analysis was performed after the purification.

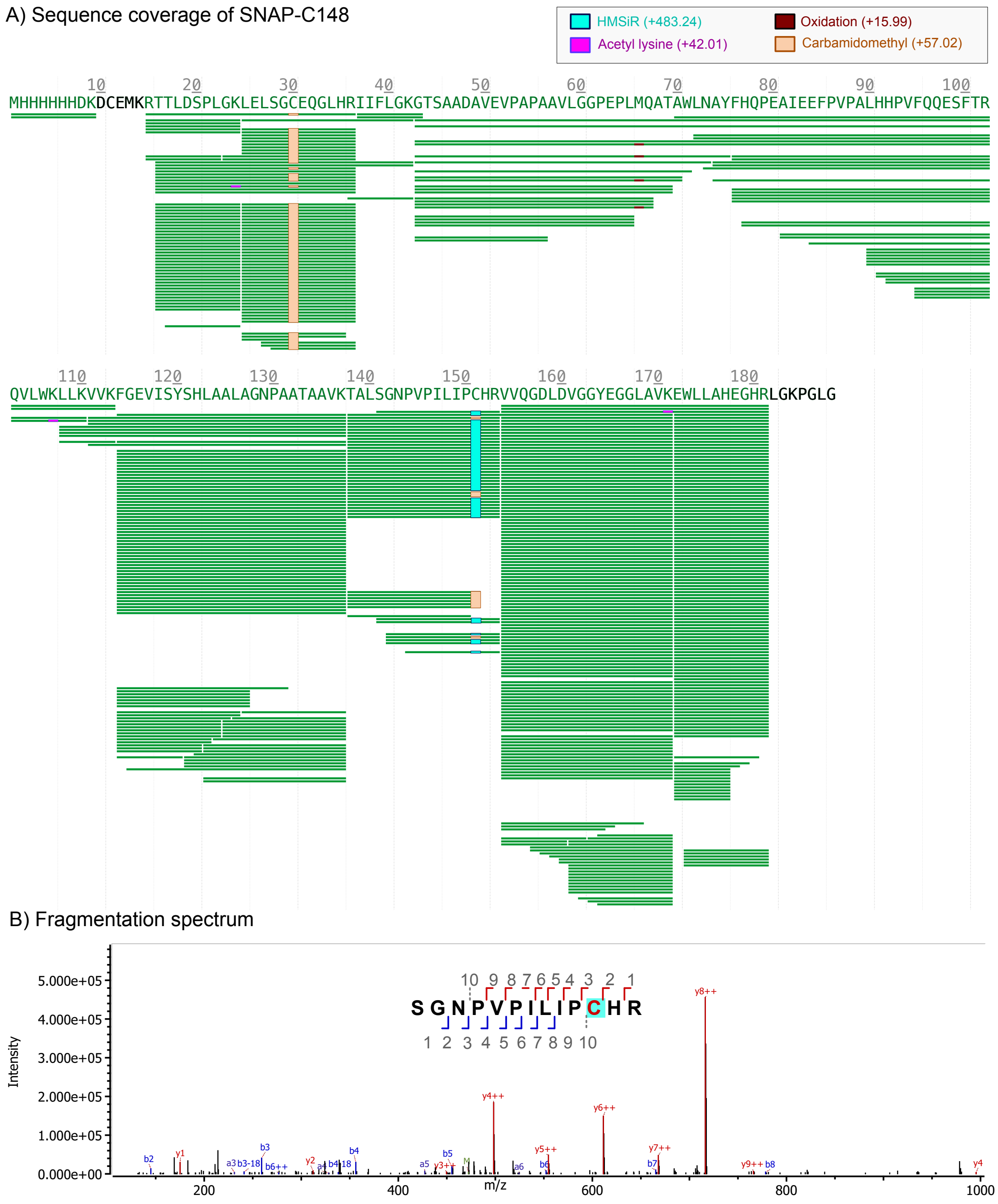

**Figure S6**. LC-MS/MS determination of the bioconjugation site of SNAP-tag labeled with HMSiR-IA. A) Protein sequence coverage map of SNAP-C148 (position C151 in sequence with *N*-terminal His6-tag). The green bars represent all identified peptide fragments after filtering for score >200 and ppm error <5. The observed modifications are highlighted in different colors: HMSiR (turquoise; +483.24), acetyl lysine (pink; +42.01), carbamidomethyl (orange; +57.02), and oxidation (dark red; +15.99). B) Fragmentation spectrum of the HMSiR-IA-labeled peptide (SGNPVPILPCHR; peptide position 141, score = 540.5, ppm error = –5.70).

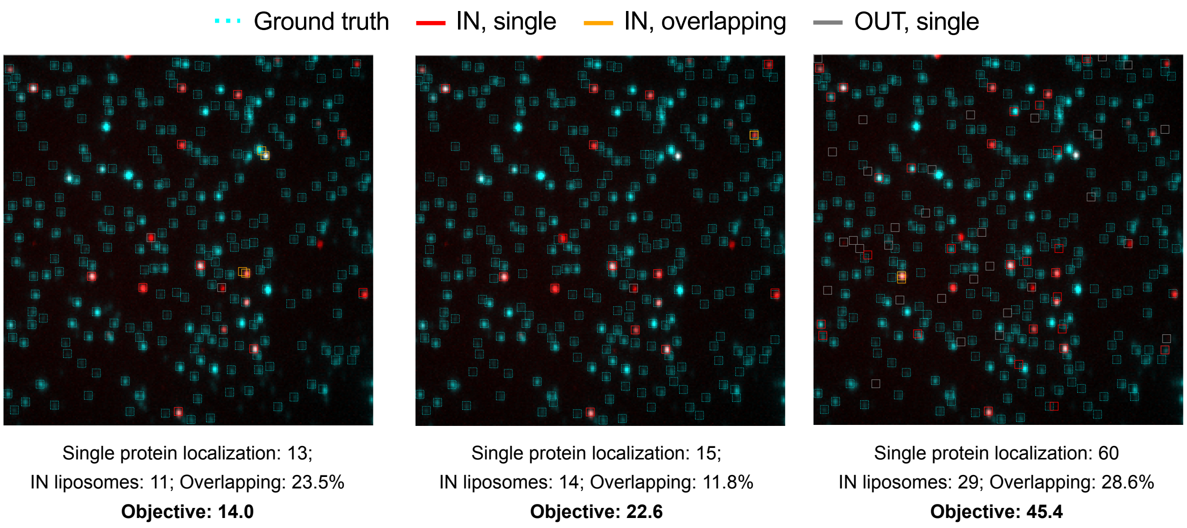

**Figure S7**. Example of Bayesian optimization of Picasso gradient parameters for single protein molecules (Halo-D106 in this example) and liposomes (DPPE-Atto520). The example displays low (left), medium (center) and high (right) values for the objective function, which maximizes the number of non-overlapping, single-molecule localizations associated with vesicles. In this case the ground truth is the vesicle signal (cyan) and the protein signal is displayed in red. Plots like this one are produced for all proteins as standard output of our extraction pipeline (https://github.com/locbp-uzh/blinkognition/tree/main/Extraction).

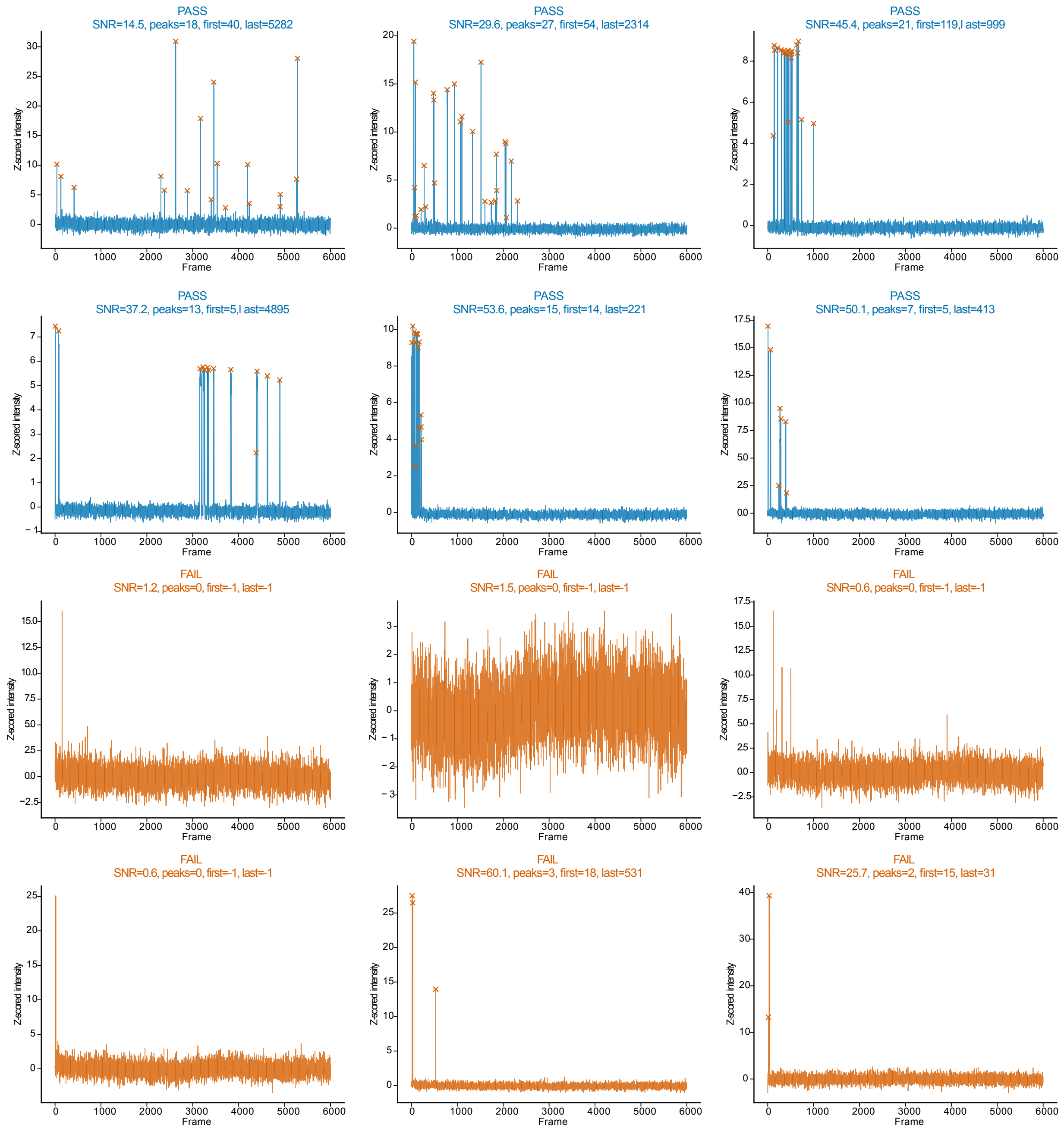

**Figure S8**. Randomly selected examples of single-molecule traces of Halo-D106 that either passed or failed our filtering criteria. Selected peaks are marked with a cross. Plots like this one are produced for all proteins as standard output of our extraction pipeline (https://github.com/locbp-uzh/blinkognition/tree/main/Extraction).

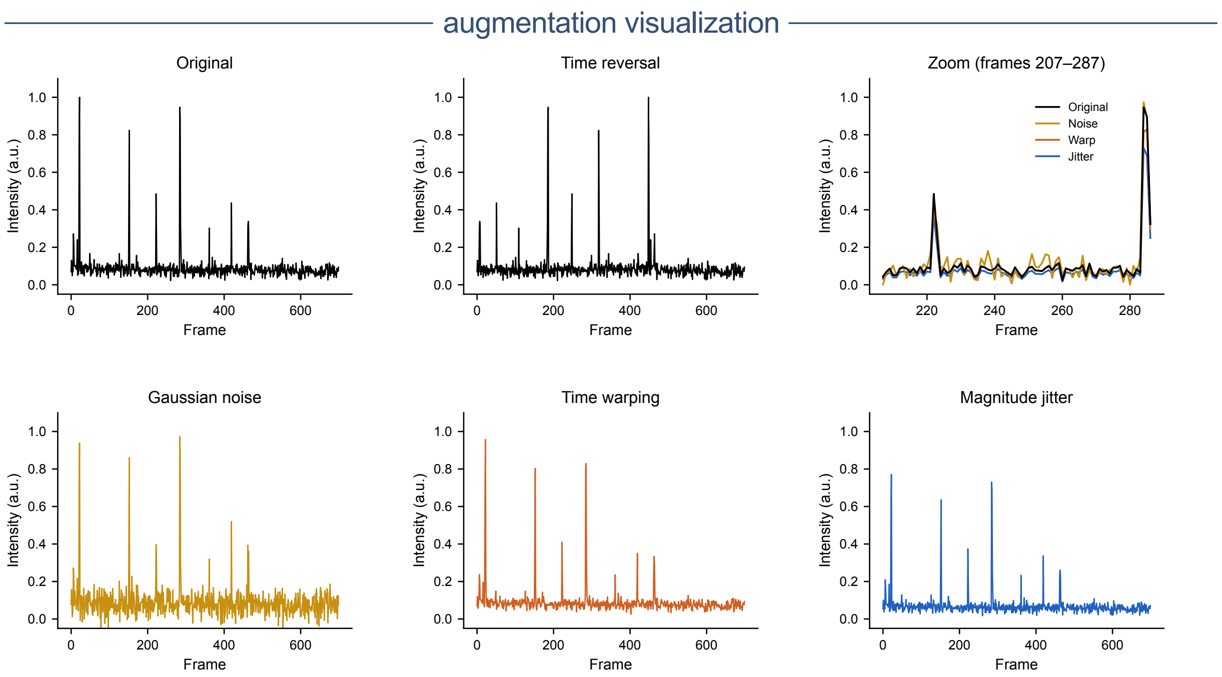

**Figure S9**. Representative example of data augmentation strategies applied to single-molecule fluorescence intensity traces. Each panel shows a single representative minmax-normalized protein trace (700 frames) together with one augmented version generated by the four strategies used during model training. Time reversal is applied only to the active window, defined as the interval from the first frame of the first detected peak to the last frame of the last detected peak. Peaks are identified by a two-component GMM (posterior probability threshold 0.9). Baseline frames before the first peak and after the last peak are left unchanged.

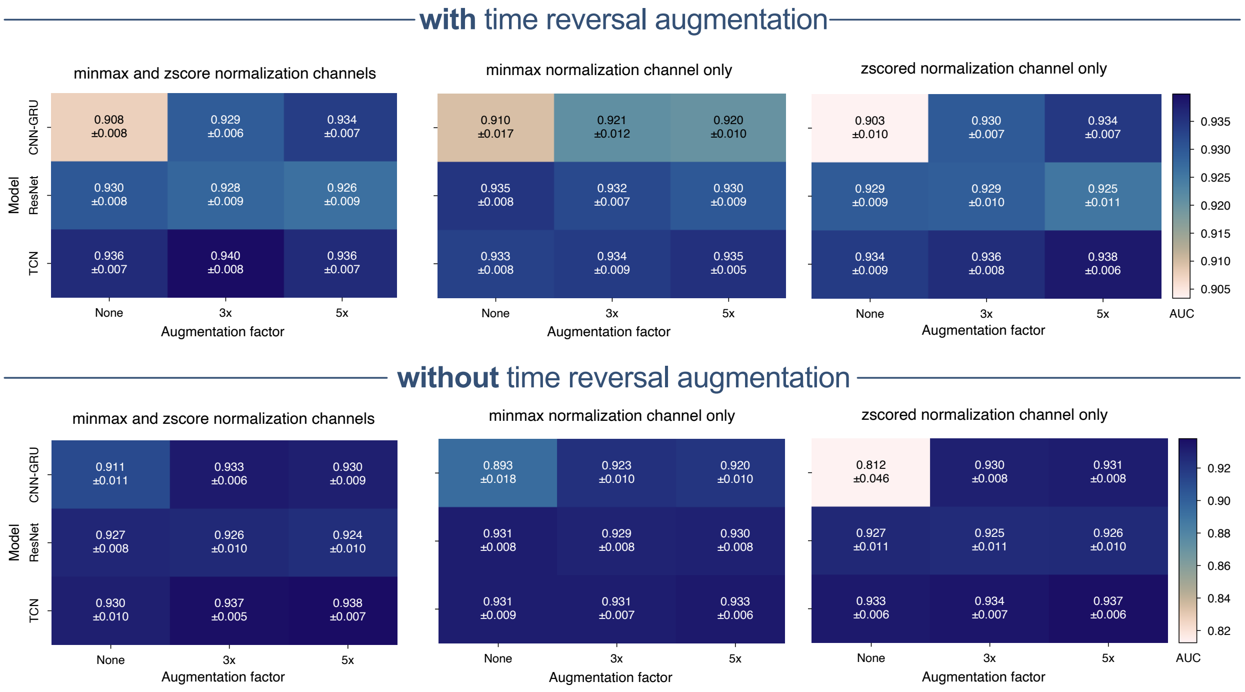

**Figure S10**. Four-fold cross validation of CNN-GRU, ResNet and TCN models for various augmentation conditions, including inclusion of time-reversed traces and injection of Gaussian noise, time warping, and magnitude jitter (sigma=0.5 in all cases) for the binary classification of Halo-D106 vs SNAP-C148.

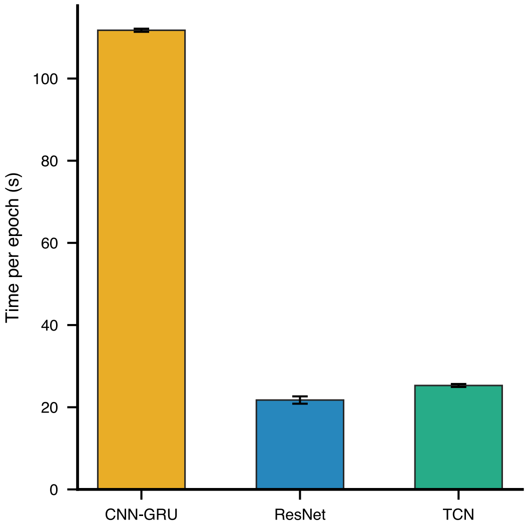

**Figure S11**. Comparison of training times of models CNN-GRU, ResNet, and TCN using time traces of 4000 frames (30 ms per frame = 120 s) in the binary classification of Halo-D106 vs SNAP-C148. All models were trained using an Nvidia A100 GPU.

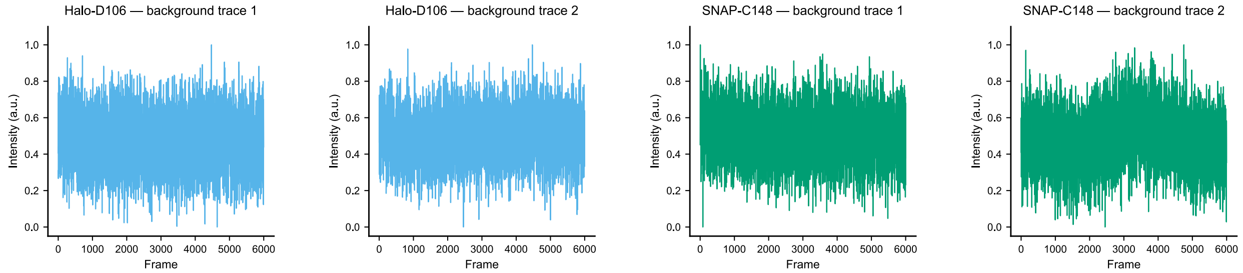

**Figure S12**. Examples of background traces from Halo-D106 or SNAP-C148 acquisitions.

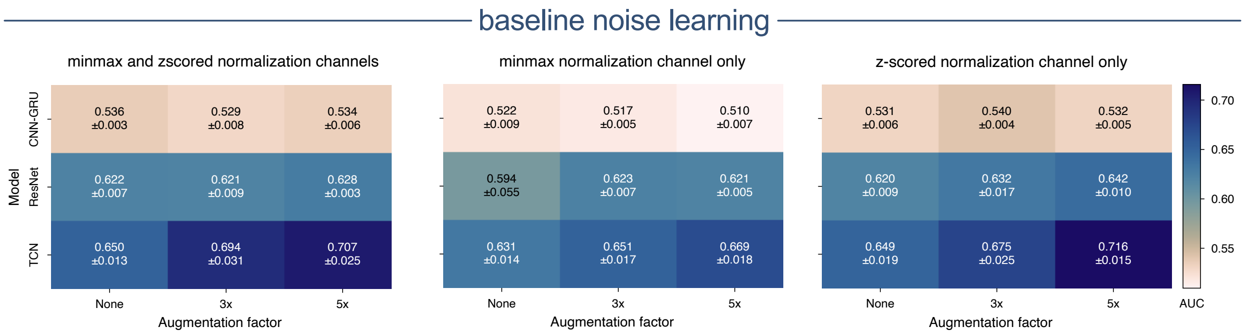

**Figure S13**. Four-fold cross validation of baseline noise learning with CNN-GRU, ResNet, and TCN models under various augmentation conditions, including injection of Gaussian noise, time warping, and magnitude jitter (sigma = 0.5 in all cases), and using one or two normalization channels.

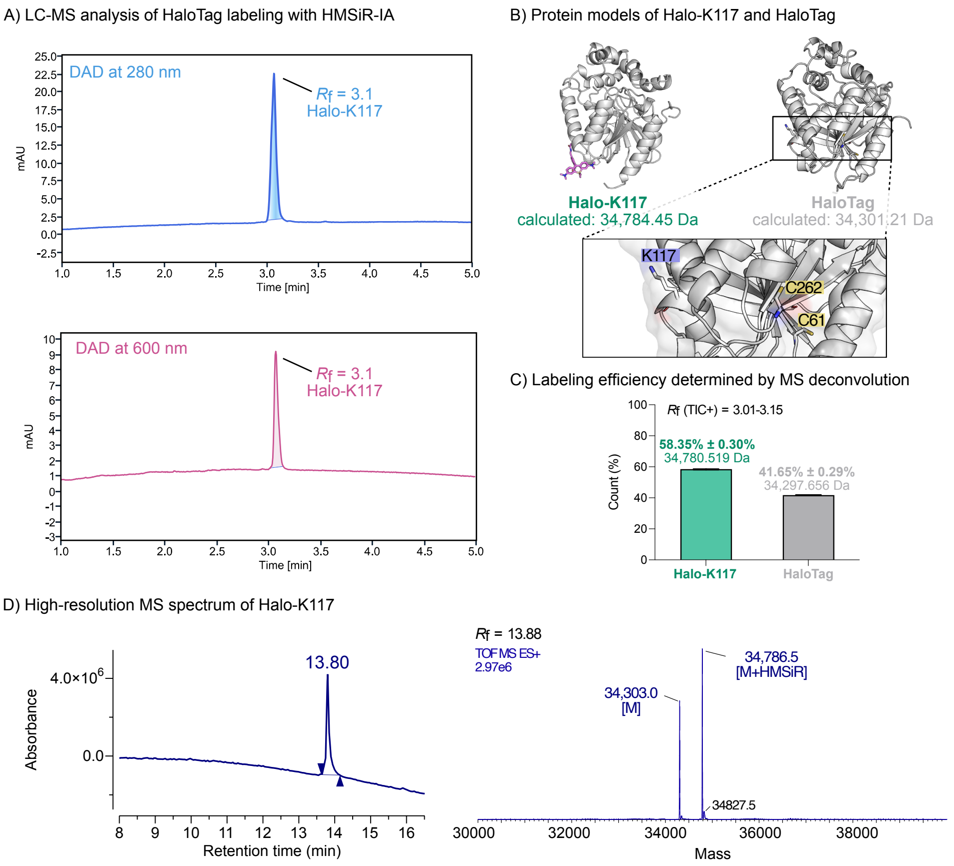

**Figure S14**. LC-MS analysis of Halo-K117 labeled with HMSiR-IA. A) Integrated protein peak ($R_f$ = 3.1) in the diode array detector (DAD) chromatogram at 280 nm (top) and 600 nm (bottom). B) Protein structure models with corresponding calculated mass for Halo-K117 (34,784.45 Da) and unlabeled HaloTag (34,301.21 Da). The labeled K117 is solvent-exposed, unlike C61 and C262, which are buried in the protein structure. Models were generated with Boltz-2[17]. C) Protein masses and labeling efficiency were determined by deconvolution of the positive total ion count (TIC+) peak ($R_f$ = 3.01-3.15) corresponding to both the labeled and unlabeled protein peaks with SAMMI (Cerno Biosciences)[4,5]. The labeling reaction was performed with 19 µM protein and 10 equiv. HMSiR-IA (4 h at 37 °C). The reaction was purified by centrifugal membrane filters. The LC-MS analysis was performed after the purification. D) Absorbance chromatogram (left) with integrated protein peak ($R_f$ = 13.80) and deconvoluted mass spectra (right) of HaloTag labeled with HMSiR-IA (Halo-K117, [M+HMSiR] = 34,786.5 Da) and unlabeled HaloTag ([M] = 34,786.5 Da). The samples were analyzed by UPLC coupled to a Synapt G2-Si mass spectrometer (30-minute run with an increasing gradient of 0.1% DFA in acetonitrile/75% 2-propanol in 0.1% DFA in water). The $R_f$ value assigned to the mass spectrum refers to the TIC chromatogram and may differ by a few digits due to the delay time between the absorbance detector and the mass spectrometer.

A) Sequence coverage of Halo-K117

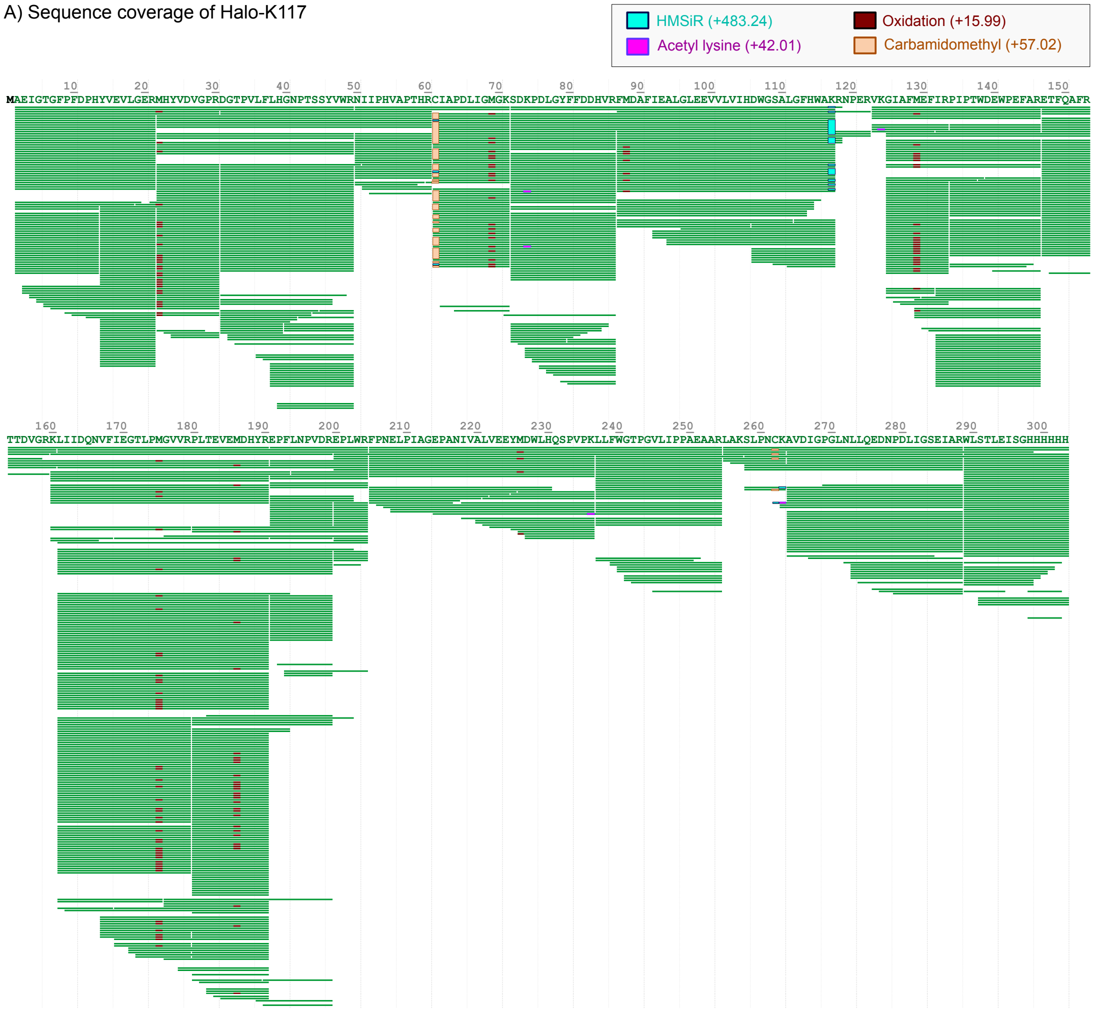

B) Fragmentation spectrum

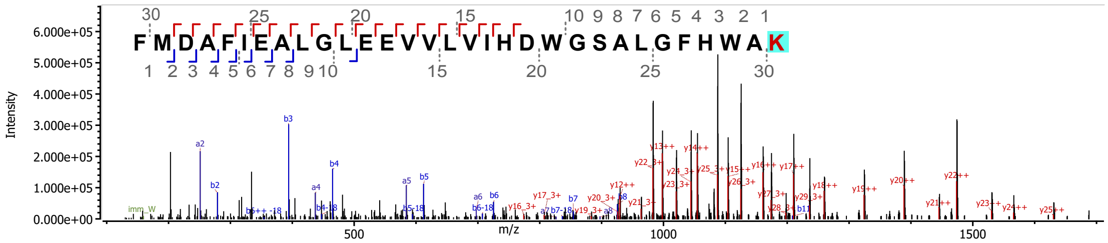

**Figure S15**. LC-MS/MS analysis of Halo-K117. A) Protein sequence coverage map of Halo-K117. The green bars represent all identified peptide fragments after filtering for score > 200 and ppm error < 5. The observed modifications are highlighted in different colors: HMSiR (turquoise; +483.24), acetyl lysine (pink; +42.01), carbamidomethyl (orange; +57.02), and oxidation (dark red; +15.99). B) Fragmentation spectrum of the HMSiR-labeled peptide (FMDAFIEALGLEEVVLVIHDWGSALGFHWAK; peptide position 87, score = 783.4, ppm error = –1.27).

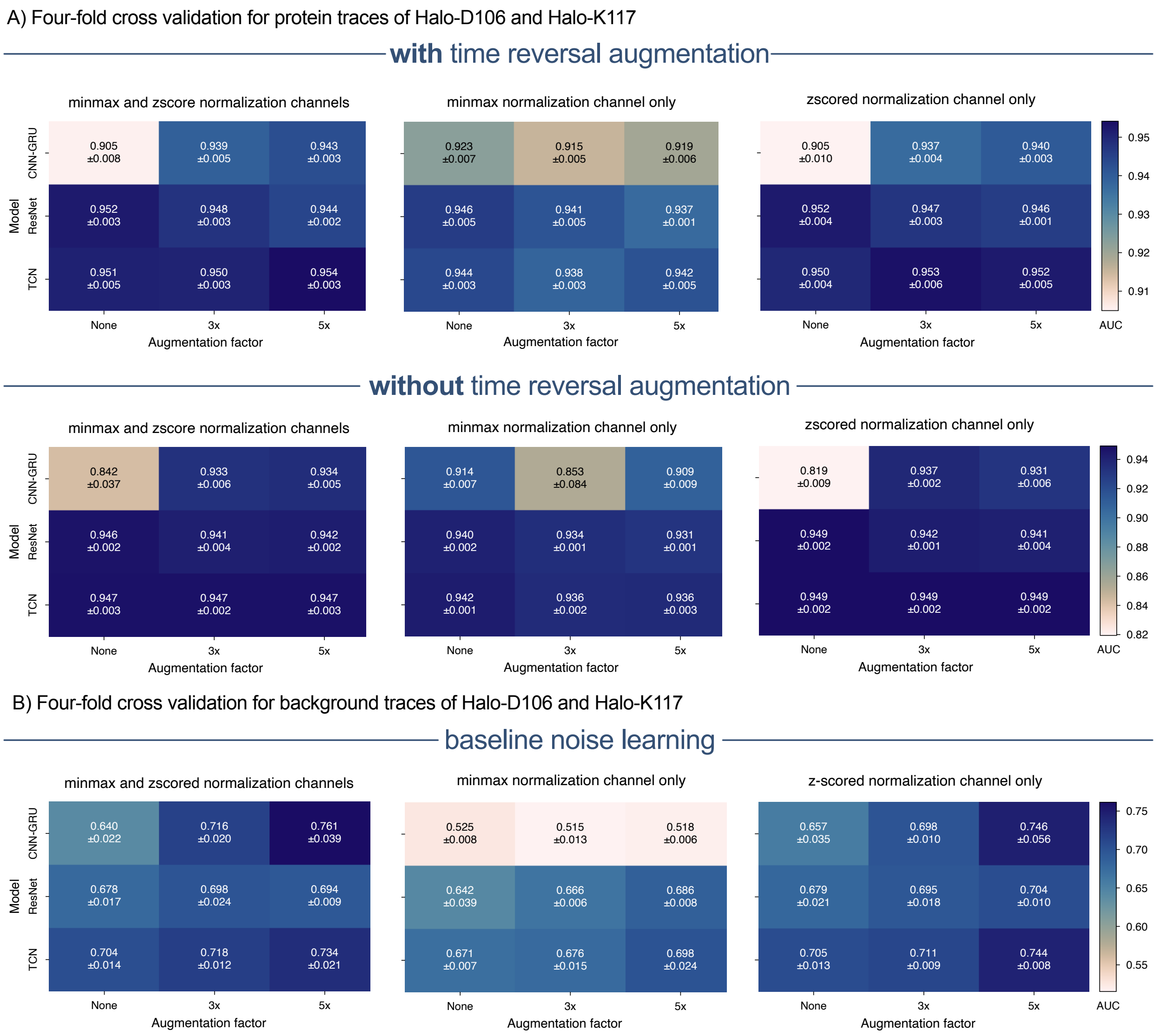

**Figure S16**. Four-fold cross validation of CNN-GRU, ResNet and TCN models for various augmentation conditions, including inclusion of time-reversed traces and injection of Gaussian noise, time warping, and magnitude jitter (sigma=0.5 in all cases) for the binary classification of Halo-D106 vs Halo-K117 and their corresponding background-only traces.

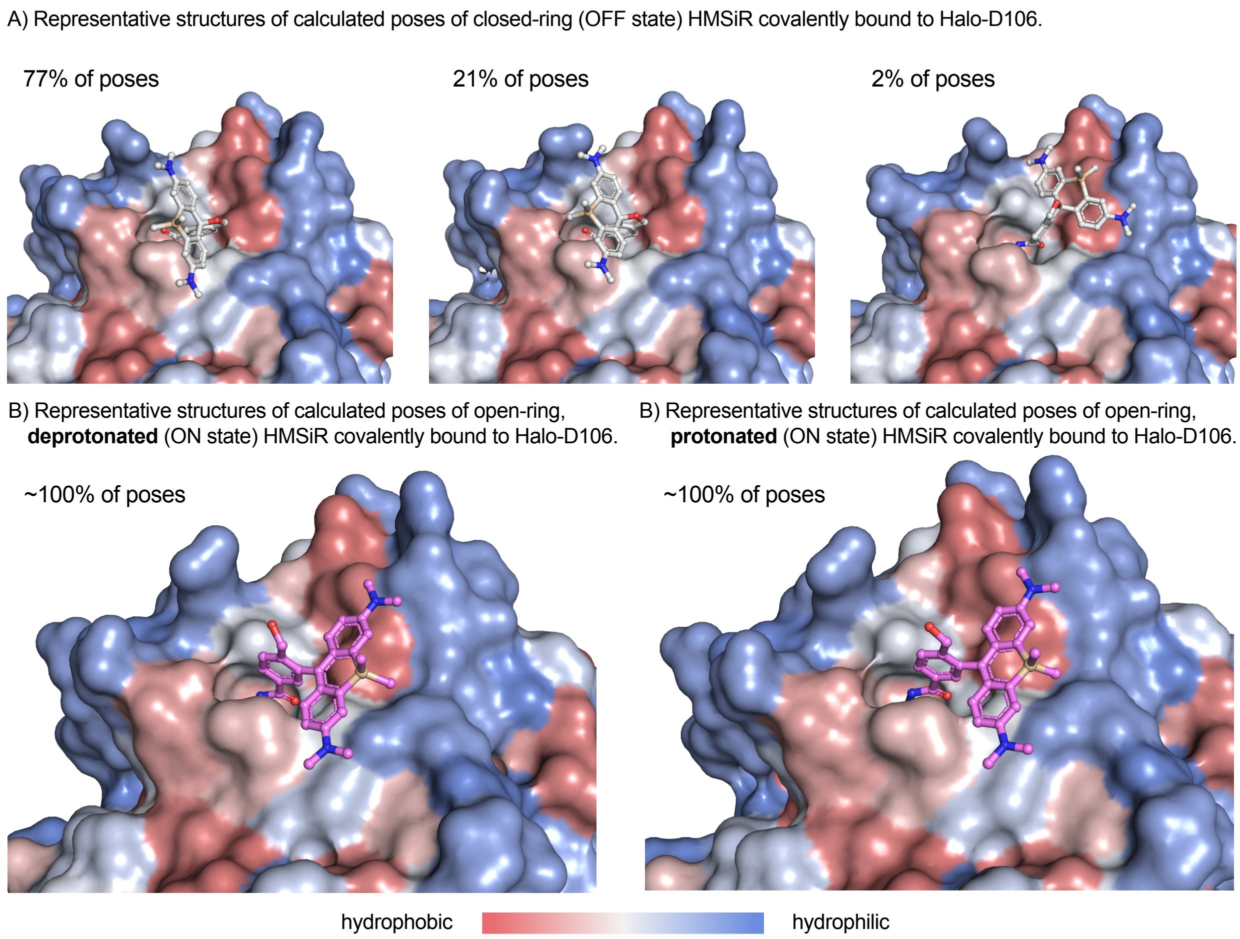

**Figure S17**. Boltz-2[17] generated poses of HMSiR in its OFF and ON states in Halo-D106. Structures displayed are medoid (most central) members of clusters obtained by average linkage hierarchical clustering with an average RMSD threshold of 5 Å on all ligand heavy atoms. Percentages represent the number of calculated poses that clustered around the displayed medoid from a total of 500 calculated poses. Protein surface colored according to the Kyte-Doolittle[19] hydrophobicity scale.

A) Representative structures of calculated poses of closed-ring (OFF state) HMSiR covalently bound to Halo-K117.

83% of poses | 15% of poses | 1% of poses

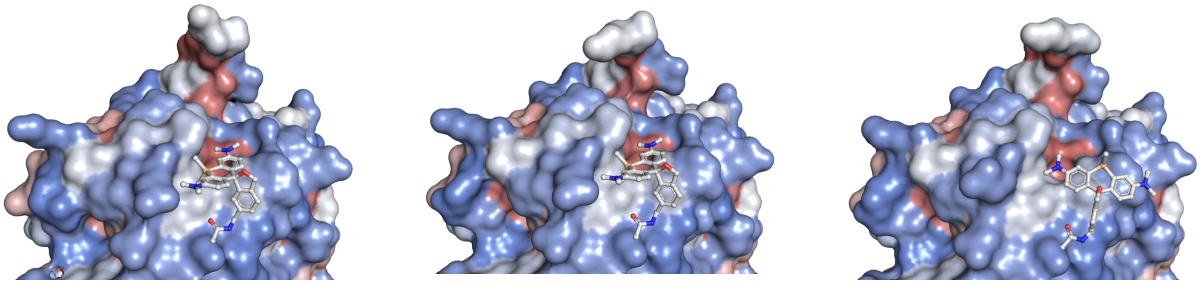

B) Representative structures of calculated poses of open-ring, **deprotonated** (ON state) HMSiR covalently bound to Halo-K117.

78% of poses | 11% of poses | 10% of poses

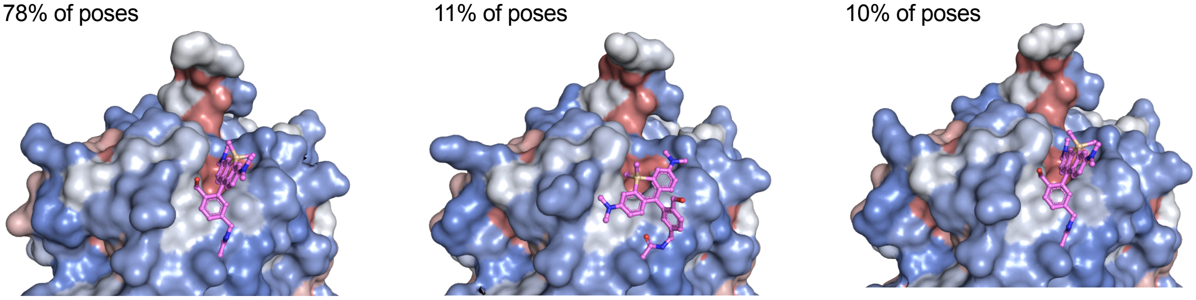

C) Representative structures of calculated poses of open-ring, **protonated** (ON state) HMSiR covalently bound to Halo-K117.

63% of poses | 22% of poses | 11% of poses

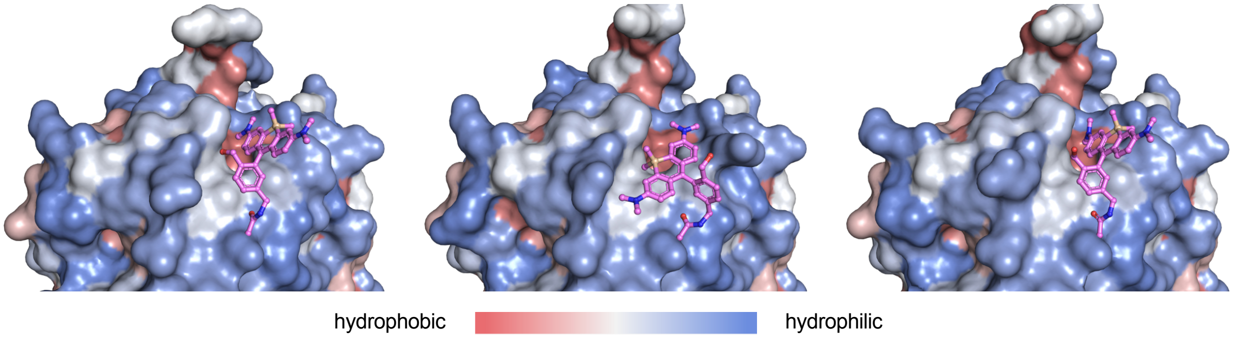

**Figure S18**. Boltz-2[17] generated poses of HMSiR in its OFF and ON states in Halo-K117. Structures displayed are medoid (most central) members of clusters obtained by average linkage hierarchical clustering with an average RMSD threshold of 5 Å on all ligand heavy atoms. Percentages represent the number of calculated poses that clustered around the displayed medoid from a total of 500 calculated poses. Protein surface colored according to the Kyte-Doolittle[19] hydrophobicity scale.

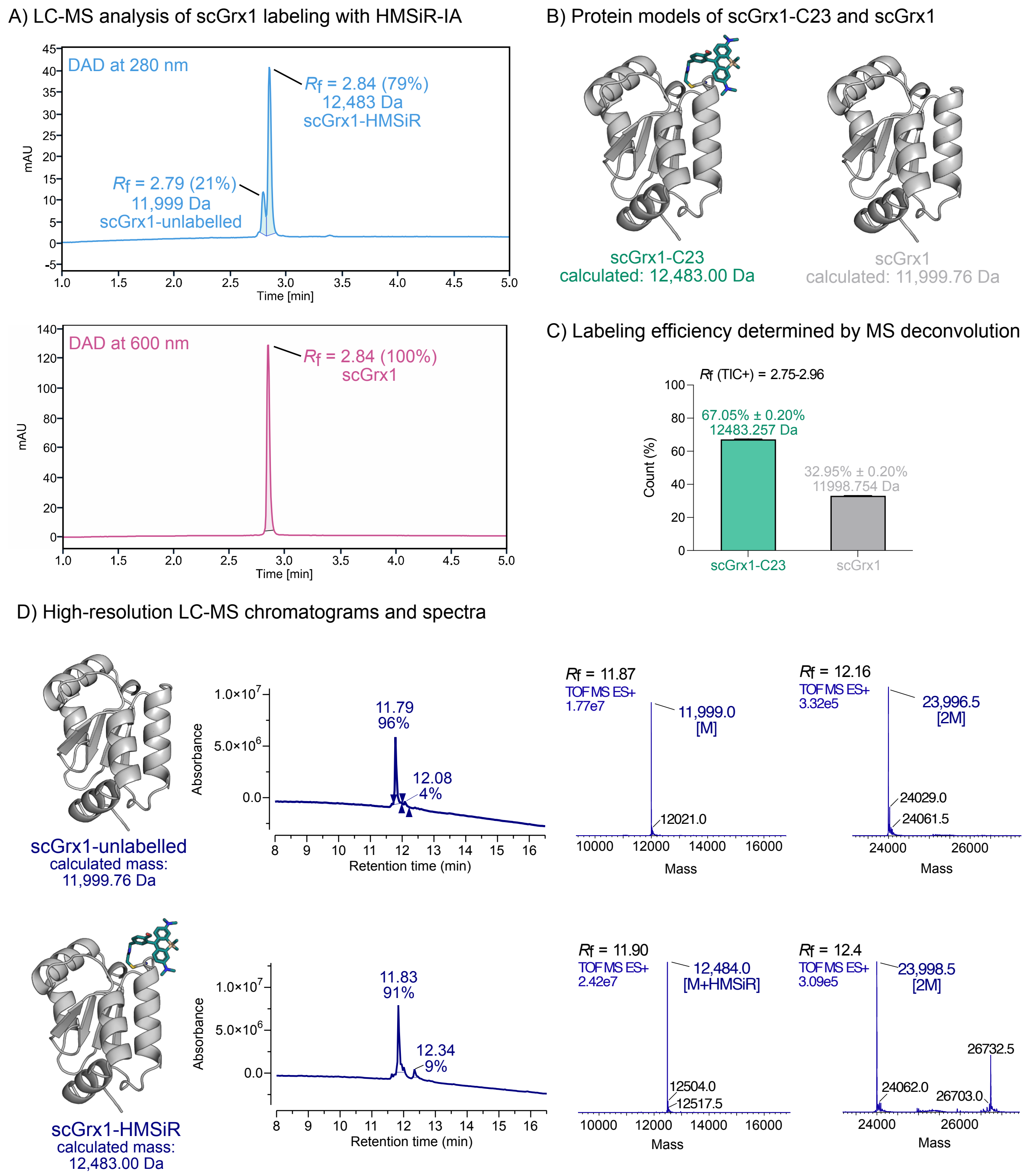

**Figure S19**. LC-MS analysis of scGrx1-C23 labeled with HMSiR-IA. A) Unlabeled ($R_f$ = 2.79, 21%, [M] = 11,999 Da) and labeled ($R_f$ = 2.84, 79%, [M+HMSiR] = 12,483 Da) protein peaks are integrated in the diode array detector (DAD) chromatogram at 280 nm (top) and 600 nm (bottom). B) Protein structure models with corresponding calculated mass for scGrx1-C23 (12,483.00) and scGrx1 (11,999.76). Models were generated with Boltz-2[17]. C) Protein masses and labeling efficiency were determined by deconvolution of the TIC+ peak ($R_f$ = 2.75-2.96) corresponding to both the labeled and unlabeled protein peaks with SAMMI (Cerno Biosciences)[4,5]. The labeling reaction was performed with 83 µM protein and 2.5 equiv. HMSiR-IA (1 h at 37 °C). The reaction was purified by SEC and concentrated with centrifugal membrane filters. The LC-MS analysis was performed after the purification. D) UHPLC-HR-ES-MS analysis of HMSiR labeled *human* scGrx1 proteins. Absorbance chromatogram (left) with integrated protein peak ($R_f$ = 11.79 and 12.08) and deconvoluted mass spectra (right) of scGrx1 ([M] = 11,999.79 Da) and its dimer ([2M] = 23,996.5 Da). Absorbance chromatogram (left) with integrated protein peak ($R_f$ = 11.83 and 12.34) and deconvoluted mass spectra (right) of scGrx1 labeled with HMSiR-IA (scGrx1-C23, [M+HMSiR] = 12,484.0 Da) and its unlabeled dimer ([2M] = 23,998.5 Da).

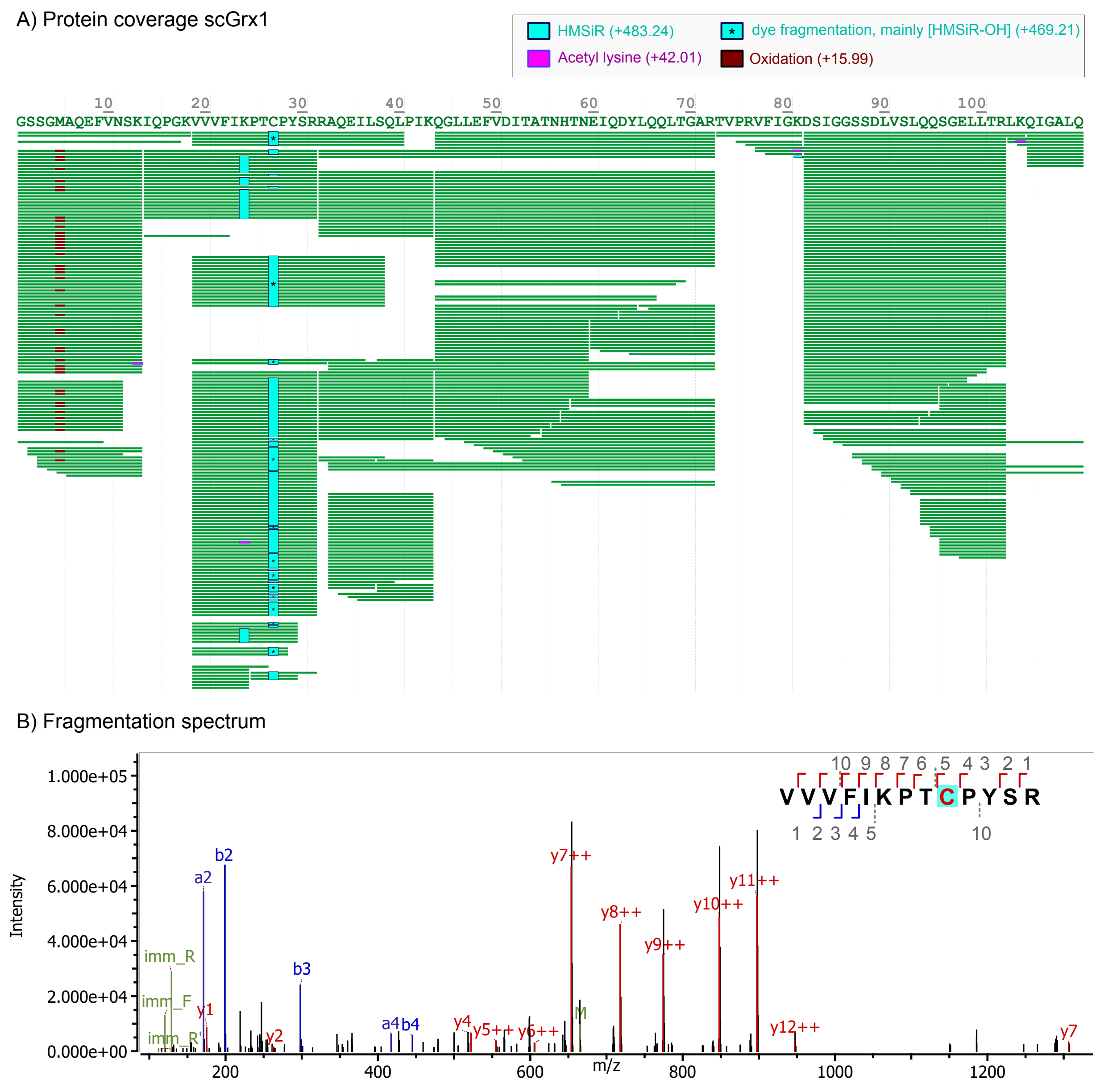

**Figure S20**. LC-MS/MS analysis of scGrx1-C23. A) Protein sequence coverage map of scGrx1-C23 (position C27 in sequence with N-terminal GSSG-linker). The green bars represent all identified peptide fragments after applying the filtering for score > 200 and ppm error < 5. The observed modifications are highlighted in different colors: HMSiR (turquoise; +483.24), HMSiR-fragmentation patterns (turquoise with *; mainly +469.21 [HMSiR-OH]), acetyl lysine (pink; +42.01), and oxidation (dark red; +15.99). B) Fragmentation spectrum of the HMSiR-labeled peptide (VVVFIKPTCPYSR; peptide position 19, score = 449.5, ppm error = –4.02).

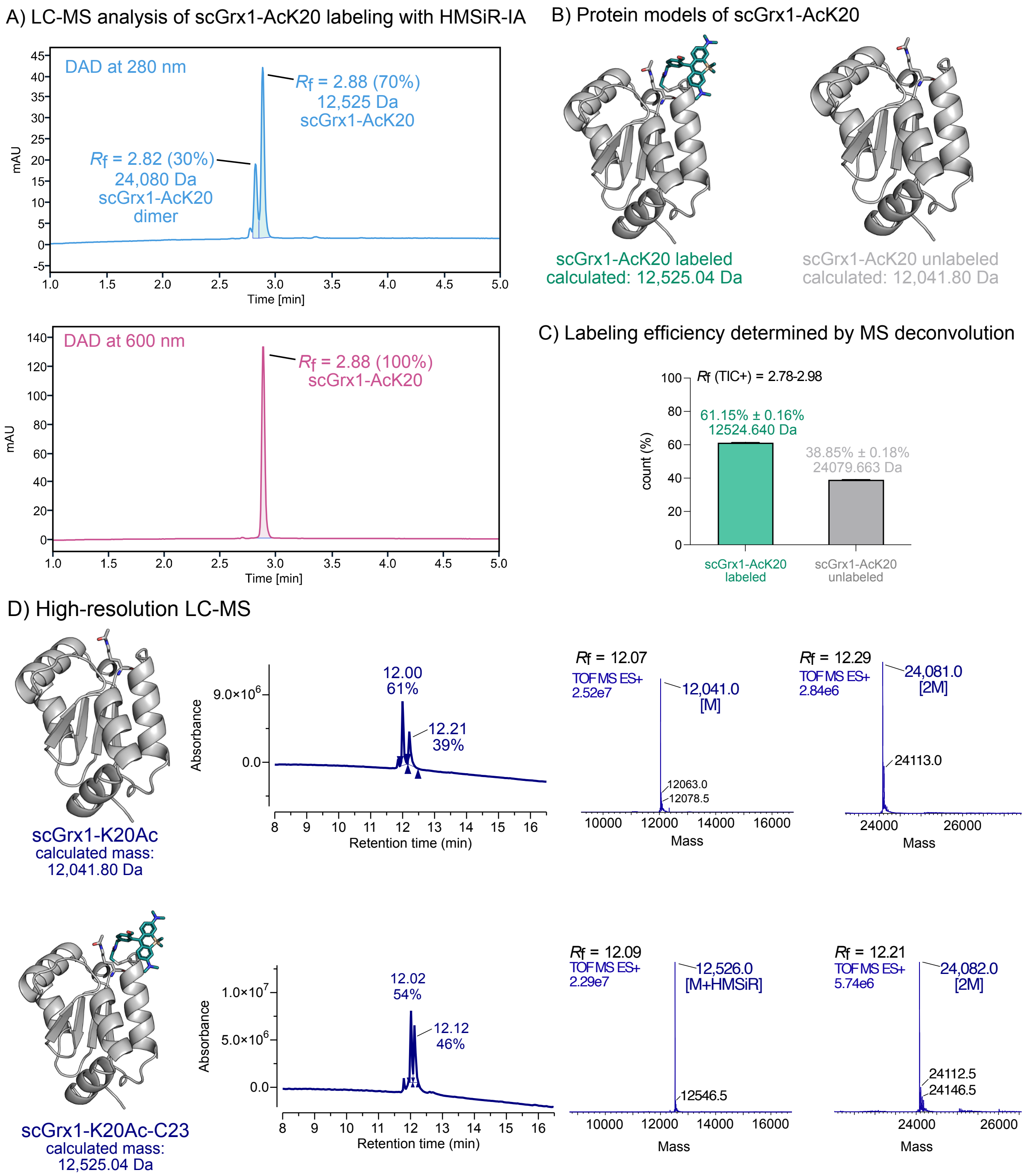

**Figure S21**. LC-MS analysis of scGrx1-AcK20 labeled with HMSiR-IA. A) Unlabeled ($R_f$ = 2.82, 30%, [2M] = 24,080 Da) and labeled ($R_f$ = 2.88, 70%, [M+HMSiR] = 12,525 Da) protein peaks are DAD chromatogram at 280 nm (top) and 600 nm (bottom). B) Protein structure models with corresponding calculated mass for unlabeled scGrx1-AcK20 (12,041.80) and HMSiR-labeled scGrx1-K20Ac (12,525.00). Models were generated with Boltz-2.[17] C) The protein masses and labeling efficiency were determined by deconvolution of the TIC+ peak ($R_f$ = 2.78-2.98) corresponding to both the labeled and unlabeled protein peaks with SAMMI (Cerno Biosciences)[4,5]. The labeling reaction was performed with 32 µM protein and 3.5 equiv. HMSiR-IA (2 h at 37 °C). The reaction was purified by SEC and concentrated with centrifugal membrane filters. The LC-MS analysis was performed after the purification. D) UHPLC-HR-ES-MS analysis of HMSiR labeled human scGrx1-AcK20. Absorbance chromatogram (left) with integrated protein peak ($R_f$ = 12.00 and 12.21) and deconvoluted mass spectra (right) of scGrx1-AcK20 ([M] = 12,041.0 Da) and its unlabeled dimer ([2M] = 24,081.0 Da). Absorbance chromatogram (left) with integrated protein peak ($R_f$ = 12.02 and 12.12) and deconvoluted mass spectra (right) of scGrx1-AcK20 labeled with HMSiR-IA (scGrx1-AcK20, [M+HMSiR] = 12,526.0 Da) and its unlabeled dimer ([2M] = 24,082.0 Da). All samples were analyzed by UPLC coupled to a Synapt G2-Si mass spectrometer (30-minute run with an increasing gradient of 0.1% DFA in acetonitrile/75% 2-propanol in 0.1% DFA in water). The $R_f$ value assigned to the mass spectrum refers to the TIC chromatogram and may differ by a few digits due to the delay time between the absorbance detector and the mass spectrometer. The protein structures shown were modeled using Boltz-2[17].

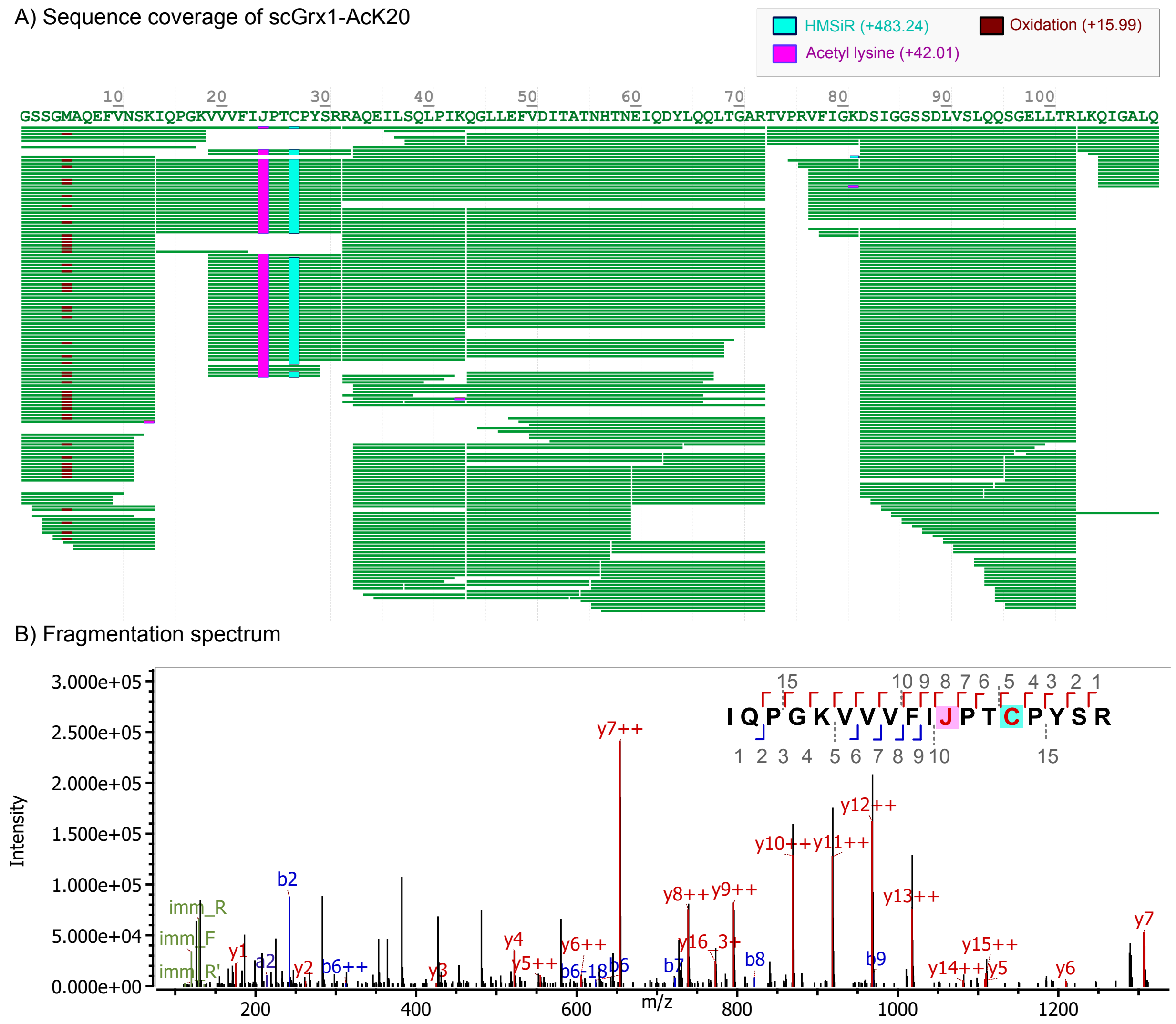

**Figure S22**. LC-MS/MS analysis of scGrx1-AcK20 labeled with HMSiR-IA. A) Protein sequence coverage map of scGrx1-AcK20 (position C27 in sequence with *N*-terminal GSSG-linker). The green bars represent all identified peptide fragments after applying the filtering for score > 200 and ppm error < 5. The observed modifications are highlighted in different colors: HMSiR (turquoise; +483.24), acetyl lysine (pink; +42.01), and oxidation (dark red; +15.99). B) Fragmentation spectrum of the HMSiR-labeled peptide (IQPGKVVVFIJPTCPYSR, with J being the post-translational modification position AcK20; peptide position 14, score = 385.1, ppm error = –6.42).

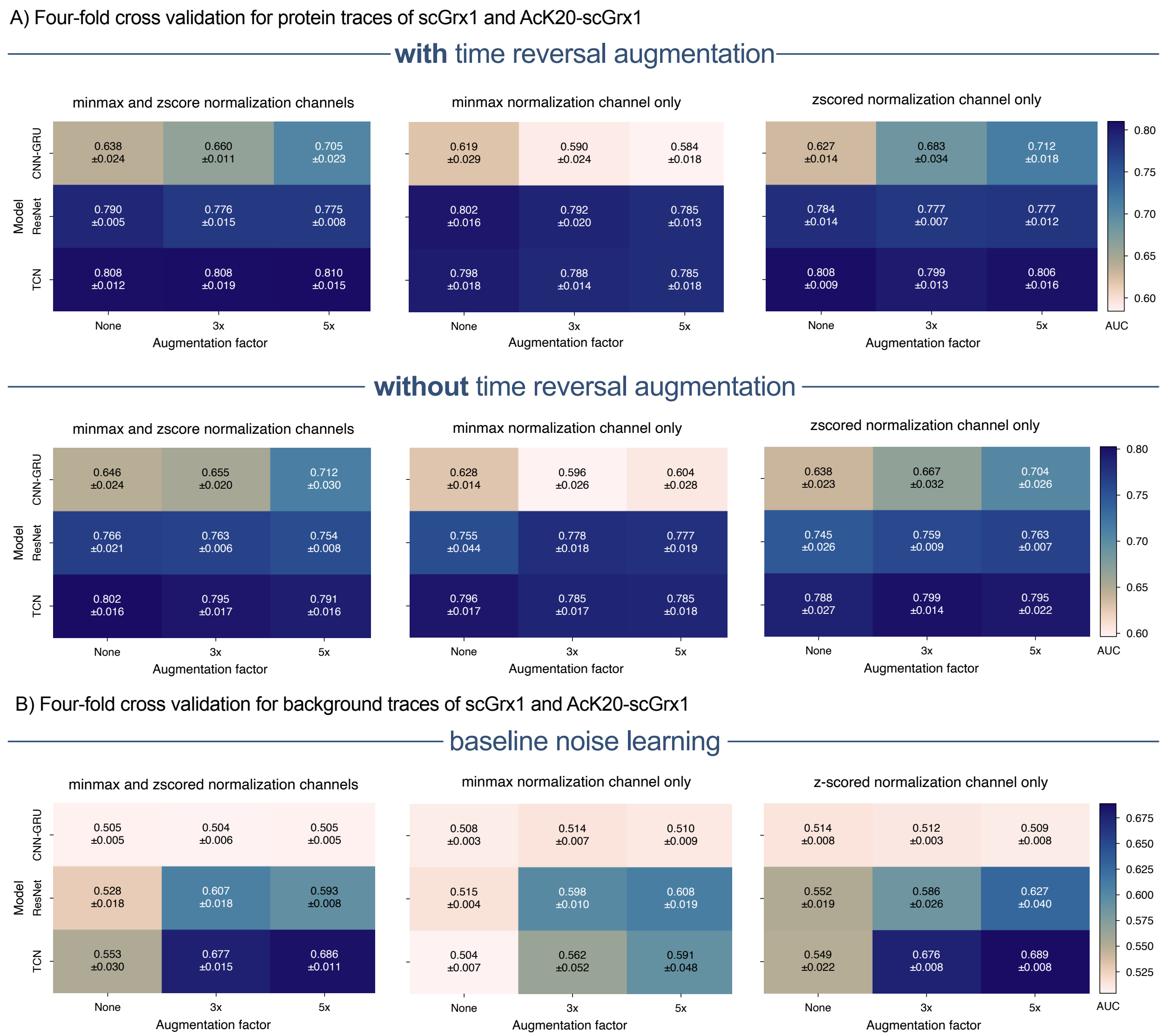

**Figure S23**. Four-fold cross validation of CNN-GRU, ResNet and TCN models for various augmentation conditions, including inclusion of time-reversed traces and injection of Gaussian noise, time warping, and magnitude jitter (sigma=0.5 in all cases) for the binary classification of scGrx1 vs AcK20-scGrx1 and their corresponding background-only traces.

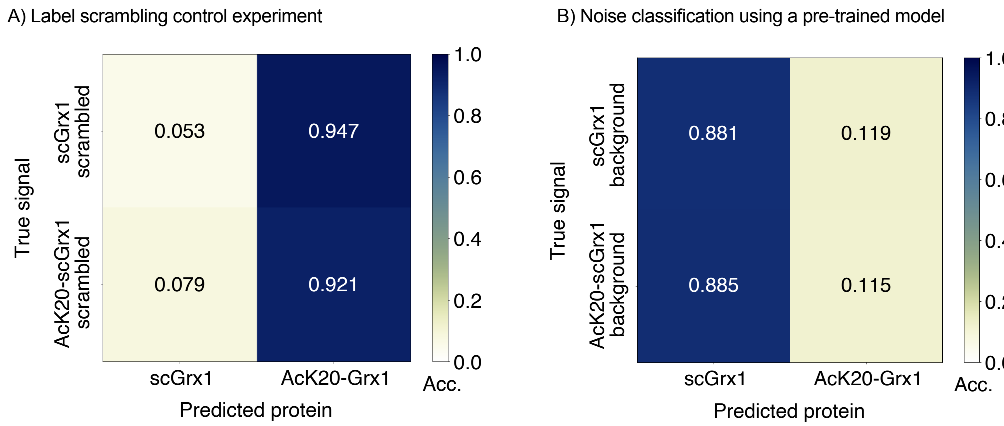

**Figure S24**. Control experiments to justify the use of the TCN model for the identification of acetylation at K20 of scGrx1. A) Label scrambling experiment: Labels of real protein traces were scrambled prior to training and classification. B) Noise classification: A TCN model was trained on real protein data and then used to classify background traces coming from the same FOVs than the corresponding protein traces. In both cases, classification collapses into a single class, indicating that the model fails to learn useful information without protein identity information or is unable to classify the background signals based on a model train on real protein data.

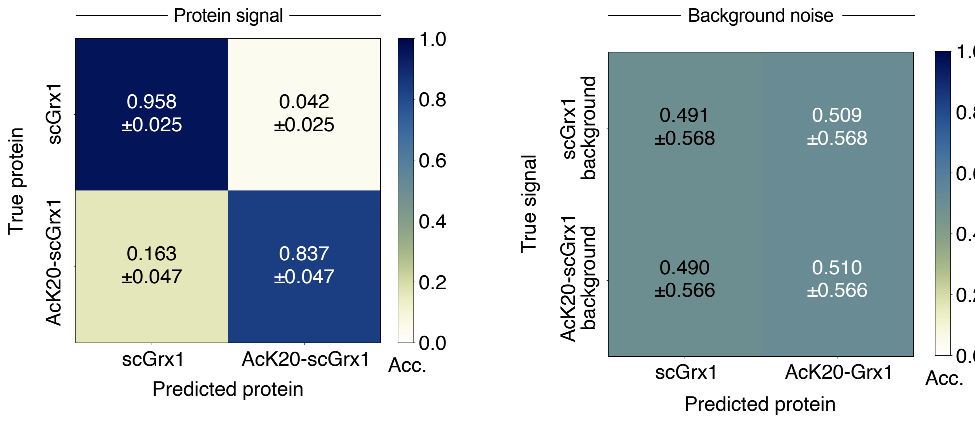

**Figure S25**. Binary classification of scGrx1 vs AcK20-scGrx1, and background training control, using the ResNet model with time reversal augmentation, minmax normalization and no further augmentation. The confusion matrices show filtered predictions and their corresponding background traces using the ResNet model with time-reversal but no further augmentation. Accuracy (Acc.) is reported as mean and standard deviation of four-fold cross validation. This result is similar to that obtained with the TCN model, suggesting that the high-certainty misclassification of Ac20K-scGrx1 as scGrx1 might be related to the deacetylation of the protein during expression and purification (Figure S23).

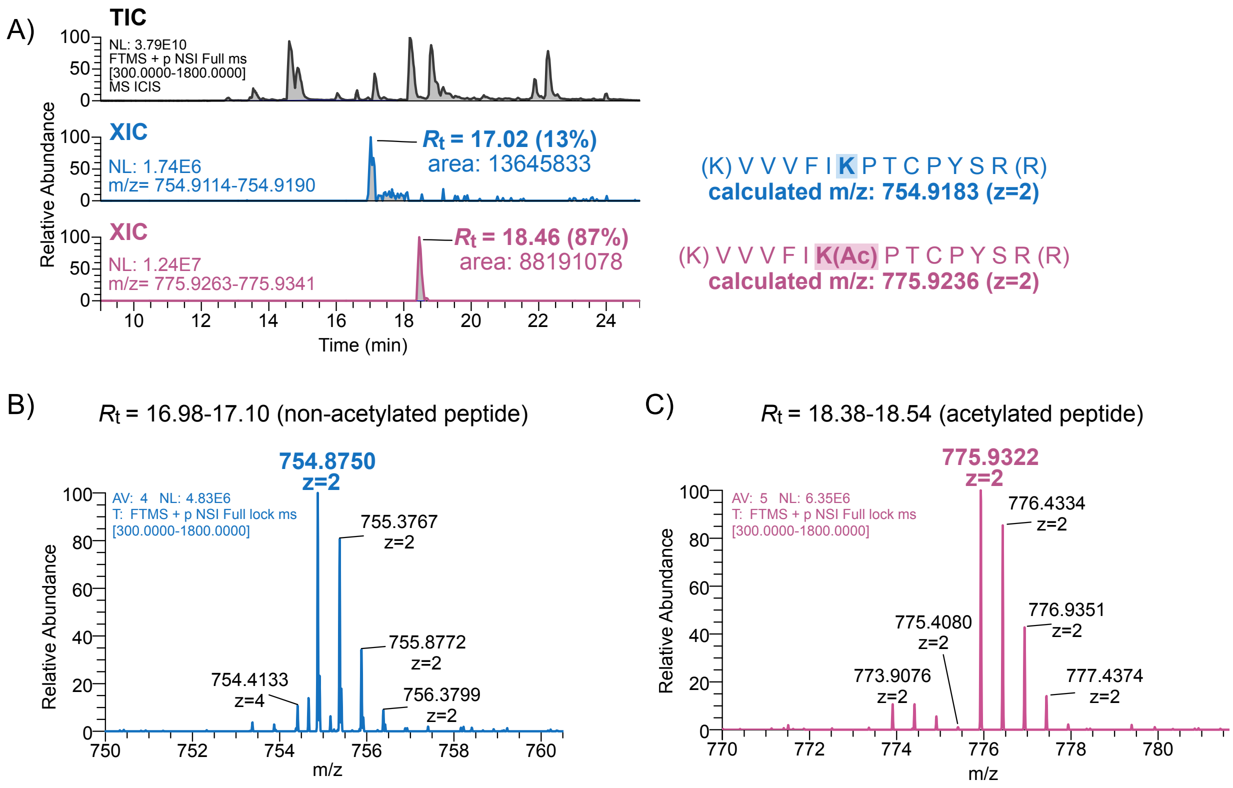

**Figure S26**. Quantification of lysine acetylation of scGrx1-K20Ac by extracted ion chromatography analysis. A) Stacked chromatograms showing TIC (black) and extracted ion chromatogram (XIC) for the non-acetylated peptide VVVFIKPTCPYSR (blue; $R_t$ = 17.02 min, 13%, calculated $[M+2H]^{2+}$=754.9183, extracted at 754.9114-754.9190) and the acetylated peptide VVVFIK(Ac)PTCPYSR (pink; $R_t$ = 18.46 min, 87%, calculated $[M+2H]^{2+}$=775.9236, extracted at 775.9263-775.9341). The peptide abundance was calculated from the relative peak areas of the XIC traces. B) Mass spectra of the non-acetylated peptide ($R_t$ = 16.98-17.10 min) with a doubly charged species centered at *m/z* 754.8750. C, Mass spectra of the acetylated peptide ($R_t$ = 18.38-18.54 min) with a doubly charged species centered at *m/z* 775.9322. All data were acquired in positive mode (FTMS, *m/z* range 300-1800), and XICs were extracted with a 5 ppm error tolerance.

A) Representative structures of calculated poses of closed-ring (OFF state) HMSiR covalently bound to scGrx1.

95% of poses | 3% of poses | 1% of poses

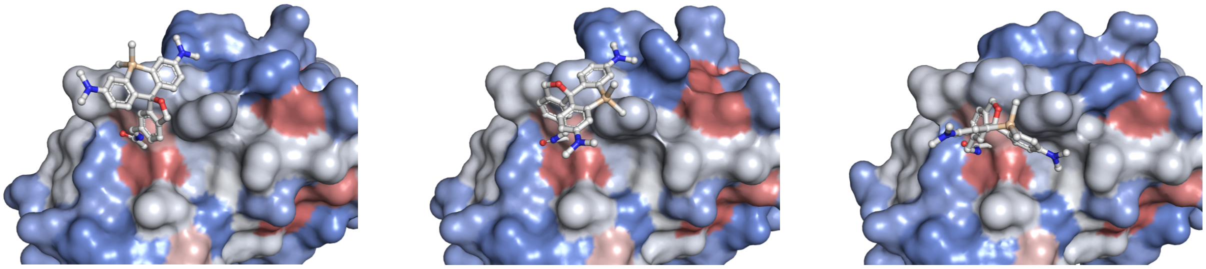

B) Representative structures of calculated poses of open-ring, **deprotonated** (ON state) HMSiR covalently bound to scGrx1.

41% of poses | 35% of poses | 14% of poses

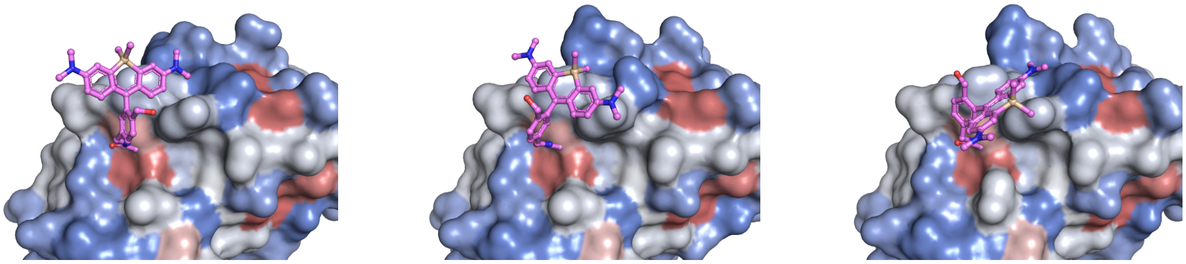

C) Representative structures of calculated poses of open-ring, **protonated** (ON state) HMSiR covalently bound to scGrx1.

85% of poses | 12% of poses | 1% of poses

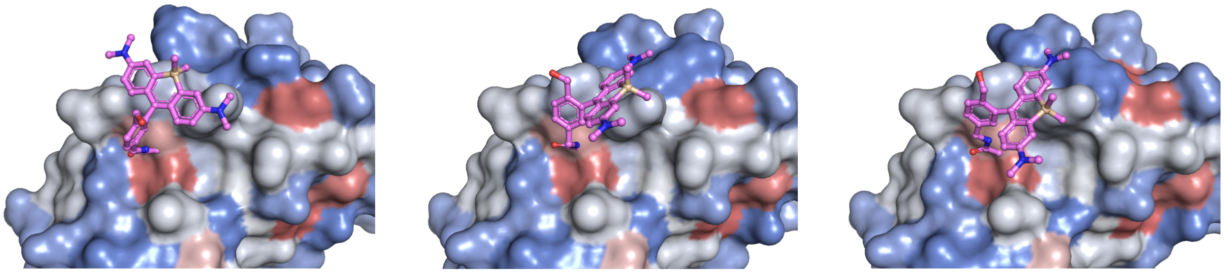

hydrophobic 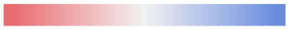 hydrophilic

**Figure S27**. Boltz-2[17] generated poses of HMSiR in its OFF and ON states in scGrx1. Structures displayed are medoid (most central) members of clusters obtained by average linkage hierarchical clustering with an average RMSD threshold of 5 Å on all ligand heavy atoms. Percentages represent the number of calculated poses that clustered around the displayed medoid from a total of 500 calculated poses. Protein surface colored according to the Kyte-Doolittle[19] hydrophobicity scale.

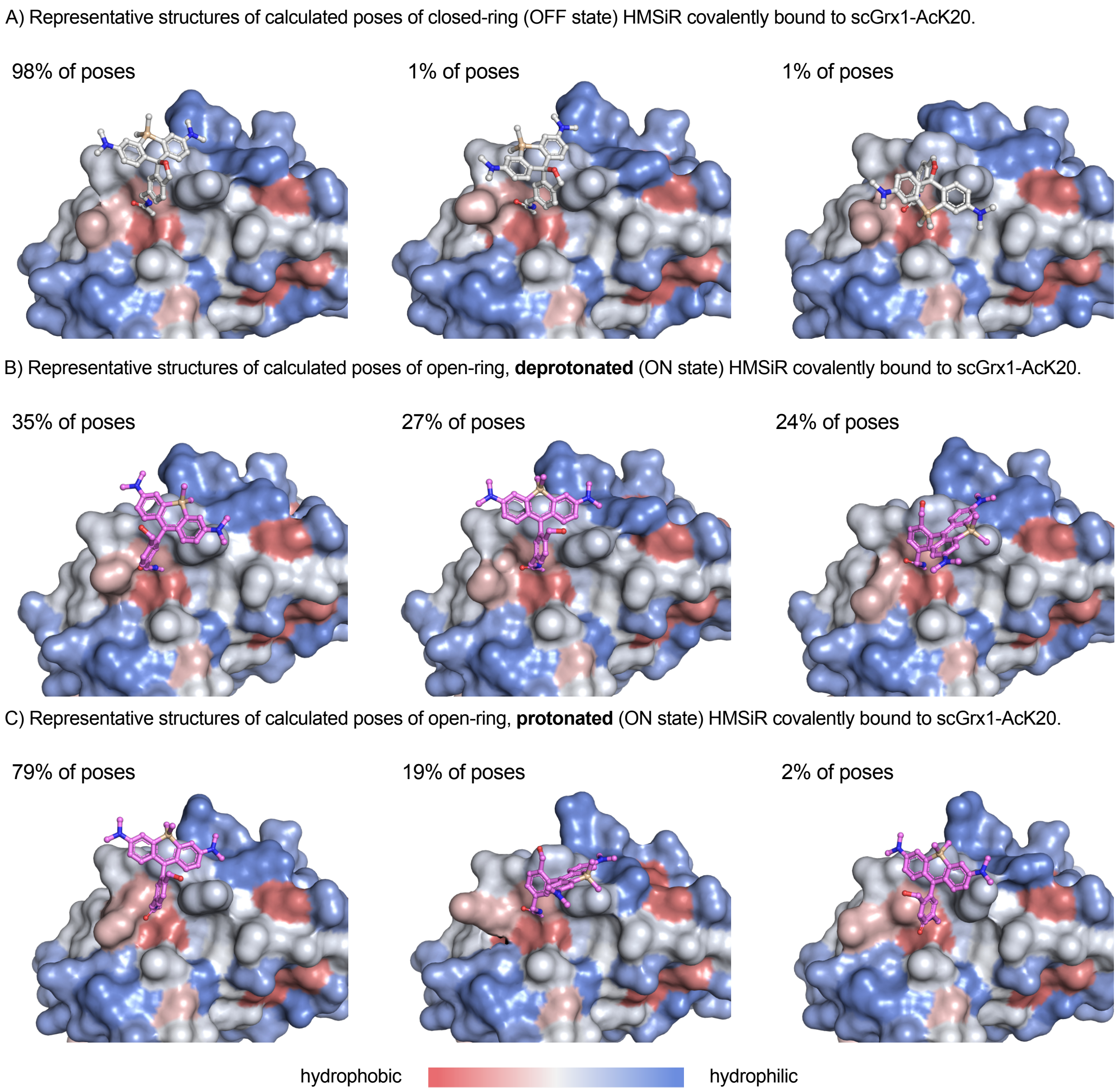

**Figure S28**. Boltz-2[17] generated poses of HMSiR in its OFF and ON states in AcK20-scGrx1. Structures displayed are medoid (most central) members of clusters obtained by average linkage hierarchical clustering with an average RMSD threshold of 5 Å on all ligand heavy atoms. Percentages represent the number of calculated poses that clustered around the displayed medoid from a total of 500 calculated poses. Protein surface colored according to the Kyte-Doolittle[19] hydrophobicity scale.

A) Random forest classification of scGrx1 vs AcK20-scGrx1 using hand-picked features

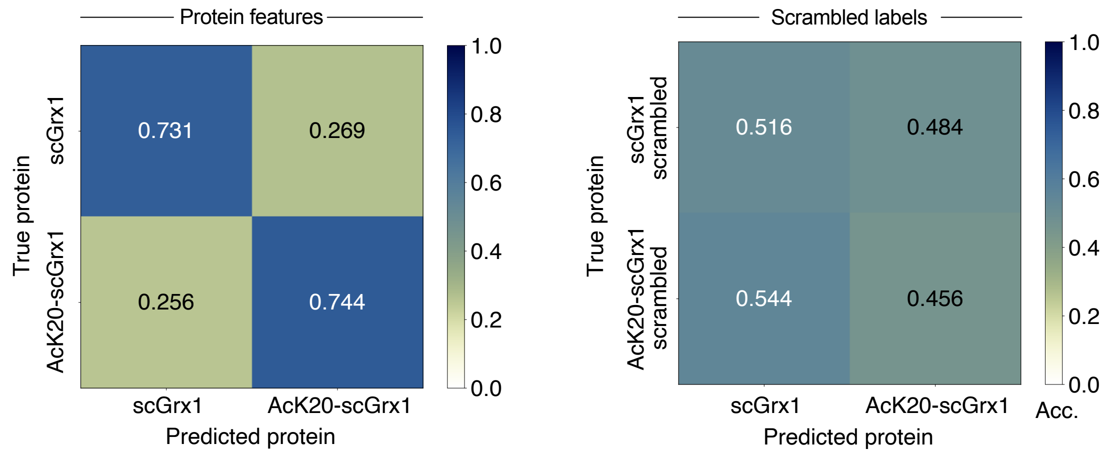

B) Gini importance for the six hand-picked features

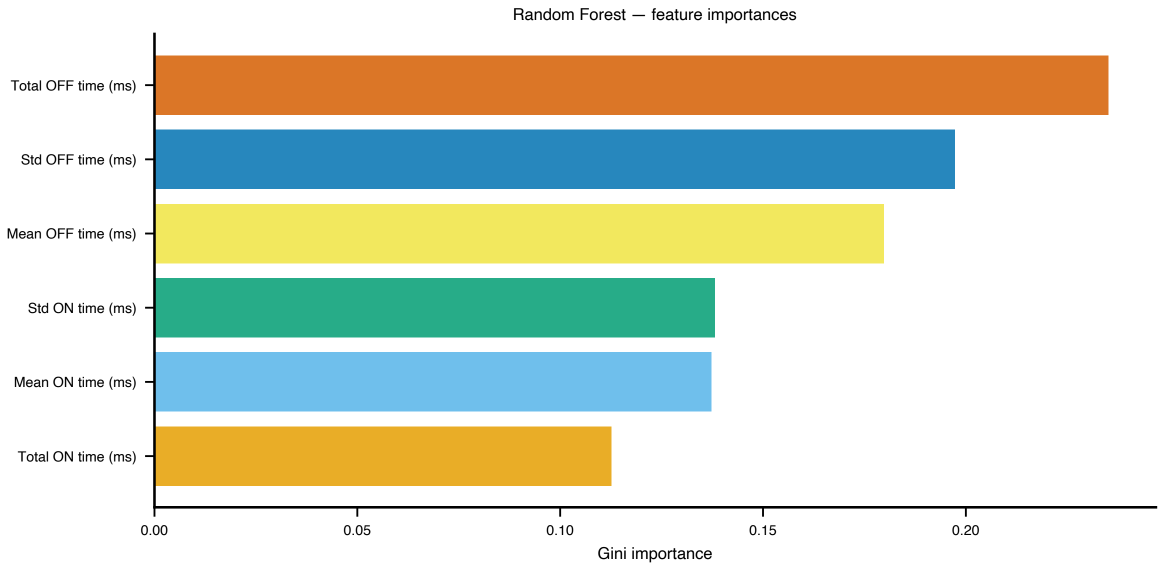

**Figure S29**. RF classification of scGrx1 vs AcK20-scGrx1 using only six selected features. A) Confusion matrices of a single training and classification task using either the actual features of proteins or features assigned to scrambled labels (control). B) Feature importances based on their mean decrease in impurity.

### Supporting Tables

**Table S1**. Reaction conditions of the labeling reactions of HaloTag, $SNAP_f$-tag, and scGrx1 protein samples.

| **Protein** | **Dye** | **Labeling time** | **Labeling efficiency determined by LC-MS** |
|---|---|---|---|
| 42 μM HaloTag | 1.1 equiv. HMSiR-CA | 2 h | 99%[a)] |
| 322 μM HaloTag | 1.1 equiv. SiR-Br | 2 h | 95%[a)] |
| 19 μM HaloTag | 10 equiv. HMSiR-IA | 4 h | 58%[a)] |
| 20 μM SNAP-tag | 5.0 equiv. HMSiR-IA | 2 h | 76%[a)] |
| 83 μM scGrx1 | 2.5 equiv. HMSiR-IA | 1 h | 67% [a)]/ 79%[b)] |
| 32 μM *s*cGrx1-AcK20 | 3.5 equiv. HMSiR-IA | 2 h | 61% [a)]/ 70%[b)] |

a) determined by spectral accurate modeling of multi-charged ions with the deconvolution algorithm SAMMI[4]

b) determined by integration of separated peaks in the Diode Array Detector (DAD) chromatogram at 280 nm

**Table S2**. Proteins used in protein labeling and classification experiments. The protein name, sequence, molecular weight, and origin of the purified protein are given. Masses were calculated using the Expasy online tool.[20] The labeled amino acid is highlighted in red.

| Protein | Protein sequence | Calculated average molecular weight (Da) | Origin |
|---|---|---|---|
| **HaloTag7[21]** <br> **(HaloTag7-6His)** | AEIGTGFPFDPHYVEVLGERMHYV DVGPRDGTPVLFLHGNPTSSYVW RNIIPHVAPTHR**C**IAPDLIGMGKSD KPDLGYFFDDHVRFMDAFIEALGL EEVVLVIHDWGSALGFHWA**K**RNP ERVKGIAFMEFIRPIPTWDEWPEFA RETFQAFRTTDVGRKLIIDQNVFIE GTLPMGVVRPLTEVEMDHYREPFL NPVDREPLWRFPNELPIAGEPANIV ALVEEYMDWLHQSPVPKLLFWGT PGVLIPPAEAARLAKSLPNCKAVDI GPGLNLLQEDNPDLIGSEIARWLST LEISGHHHHHH | 34,301.21 | Recombinant expression. <br> Initial samples: <br> Courtesy of Dr. Henriette Lämmermann and Dr. Sarah Emmert |
| **SNAP-tag[22]** <br> **(6His-SNAP$_f$-tag)** | MHHHHHHDKDCEMKRTTLDSPLG KLELSGCEQGLHRIIFLGKGTSAAD AVEVPAPAAVLGGPEPLMQATAWL NAYFHQPEAIEEFPVPALHHPVFQ QESFTRQVLWKLLKVVKFGEVISY SHLAALAGNPAATAAVKTALSGNP VPILIP**C**HRVVQGDLDVGGYEGGL AVKEWLLAHEGHRLGKPGLG | 20,265.32 | Courtesy of Dr. Annabell Martin |

| | | | |
|---|---|---|---|
| ***Human* scGrx1 (GSSG-scGrx1).**<br>**UniProt: P35754.** | GSSGMAQEFVNSKIQPGKVVVFIKPT**C**PYSRRAQEILSQLPIKQGLLEFVDITATNHTNEIQDYLQQLTGARTVPRVFIGKDSIGGSSDLVSLQQSGELLTRLKQIGALQ | 11,999.76 | Recombinant expression. DNA was supplied by *Twist Bioscience.* |
| ***Human* scGrx1 (GSSG-scGrx1-K20Ac)** | GSSGMAQEFVNSKIQPGKVVVFI**K(Ac)**PT**C**PYSRRAQEILSQLPIKQGLLEFVDITATNHTNEIQDYLQQLTGARTVPRVFIGKDSIGGSSDLVSLQQSGELLTRLKQIGALQ | 12,041.80 | Recombinant expression. DNA was supplied by *Twist Bioscience.* |

### NMR spectra

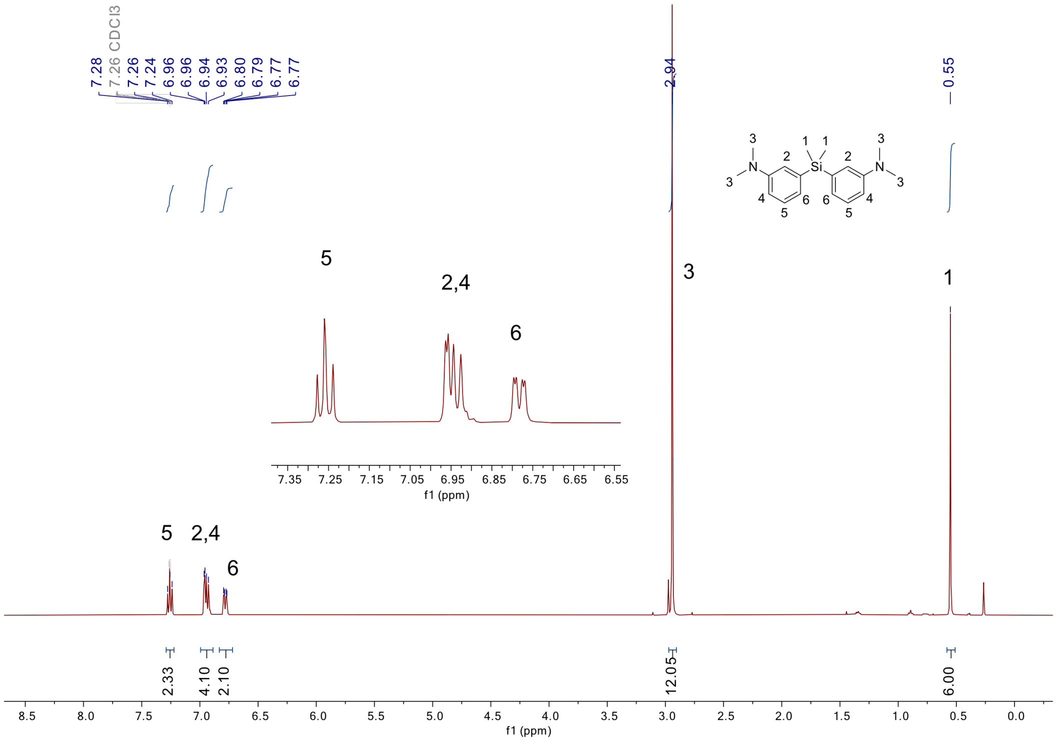

**Figure S30**. $^{1}$H NMR ($CDCl_3$, 400 MHz) spectrum of compound **S1.**

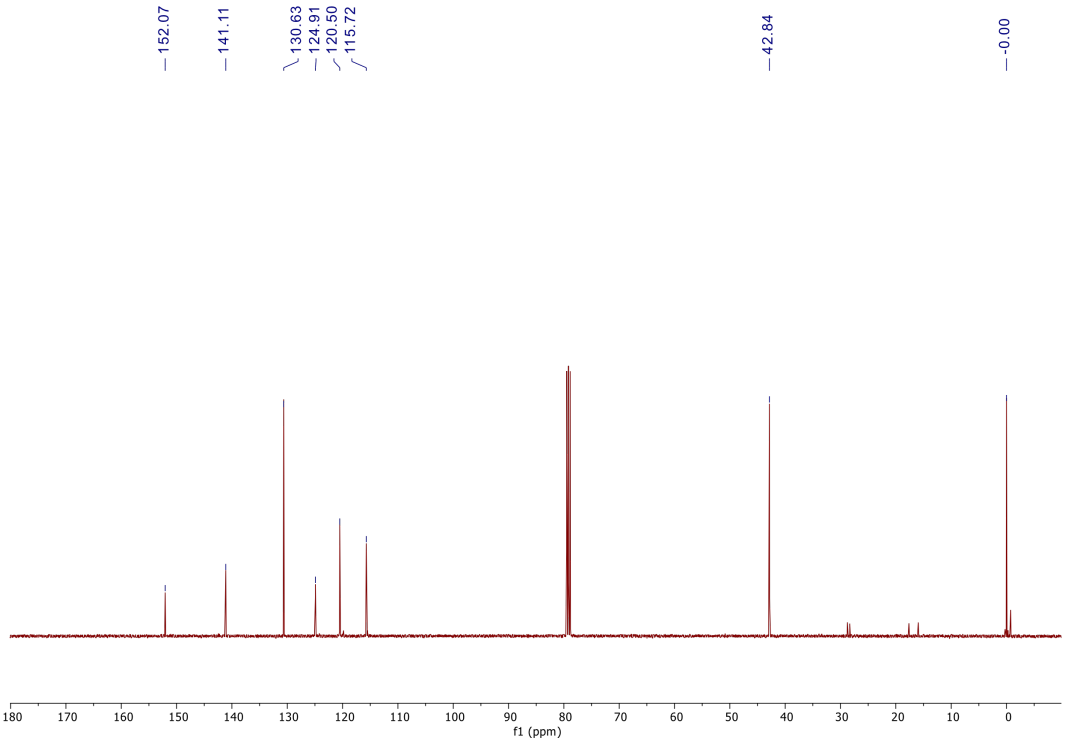

**Figure S31**. $^{13}$C NMR ($CDCl_3$, 101 MHz) spectrum of compound **S1.**

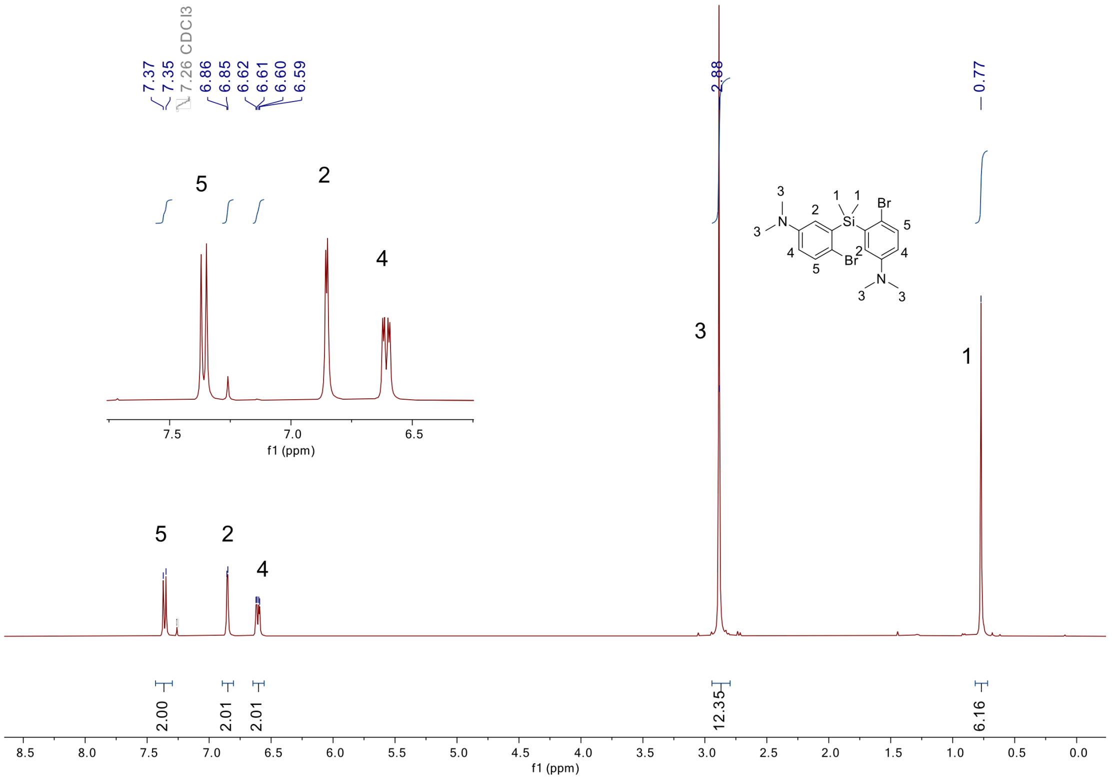

**Figure S32**. $^{1}$H NMR ($CDCl_3$, 400 MHz) spectrum of compound **S2.**

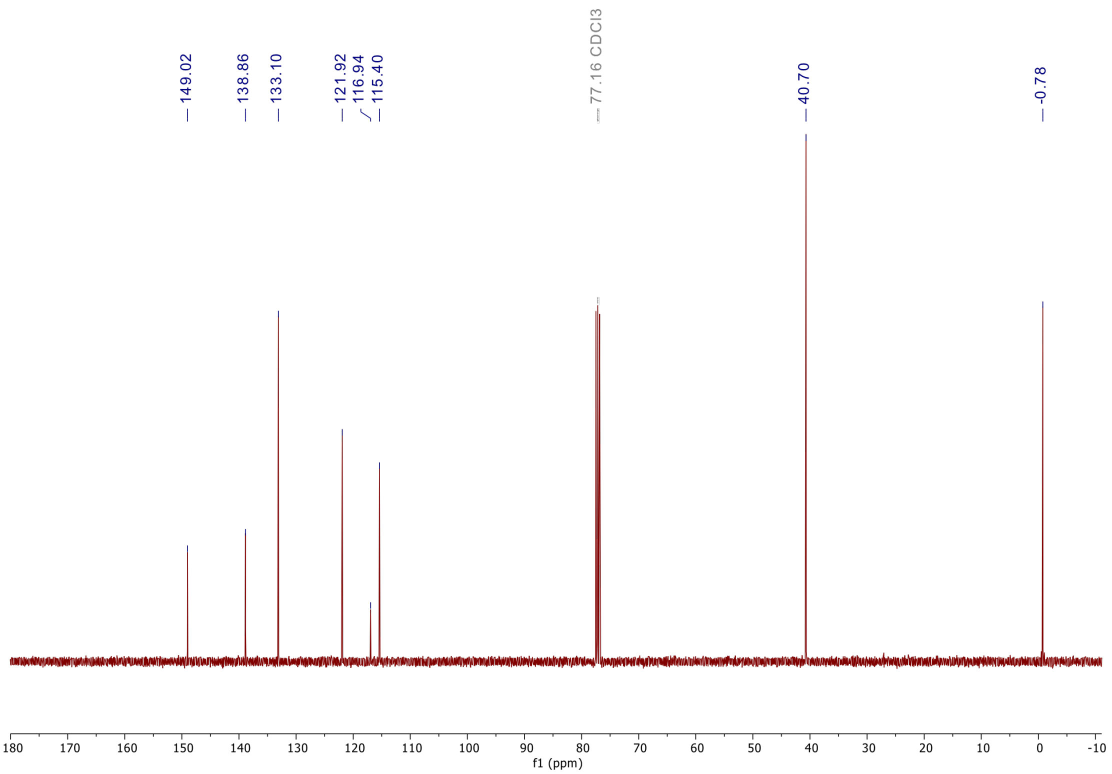

**Figure S33**. $^{13}$C NMR ($CDCl_3$, 101 MHz) spectrum of compound **S2.**

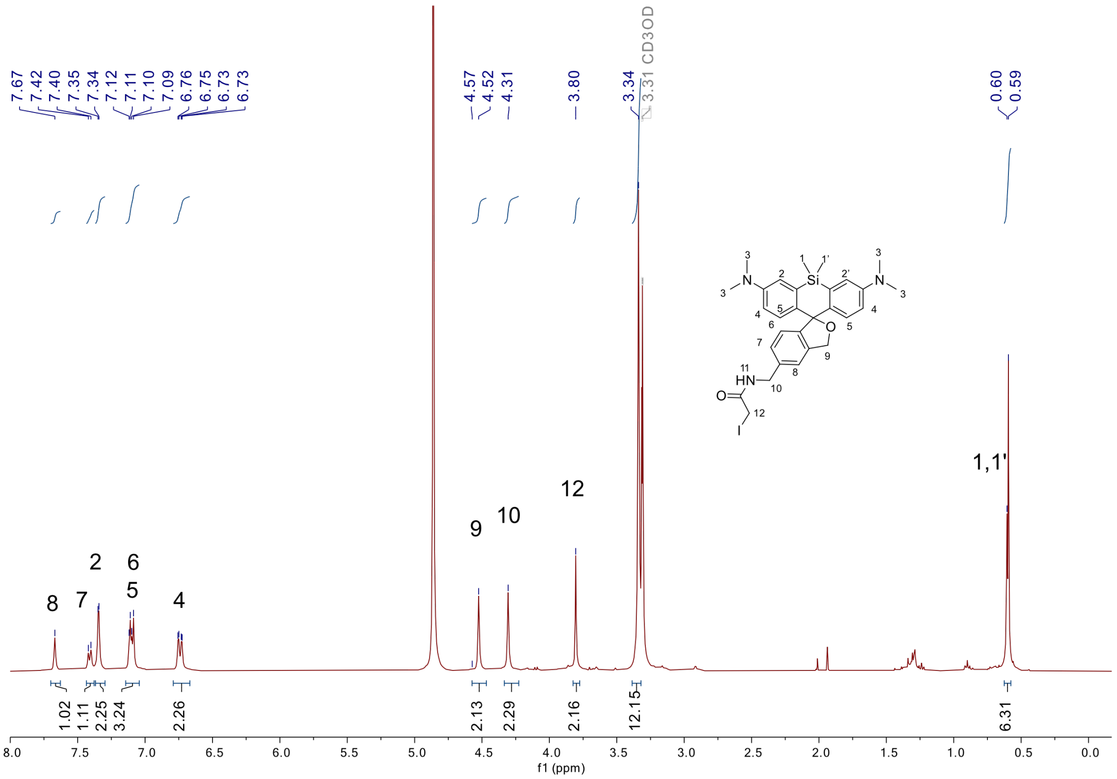

**Figure S34**. $^{1}H$ NMR ($CD_3OD$, 400 MHz) spectrum of compound **HMSiR-IA.**

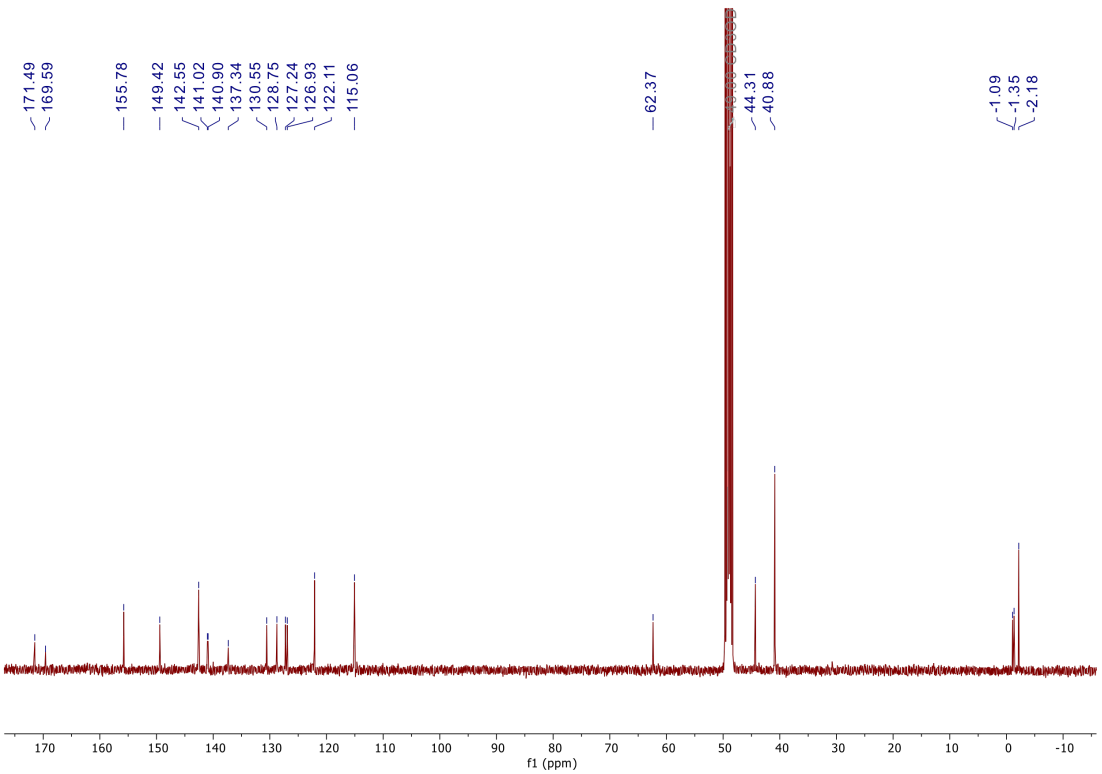

**Figure S35**. $^{13}C$ NMR ($CD_3OD$, 101 MHz) spectrum of compound **HMSiR-IA.**

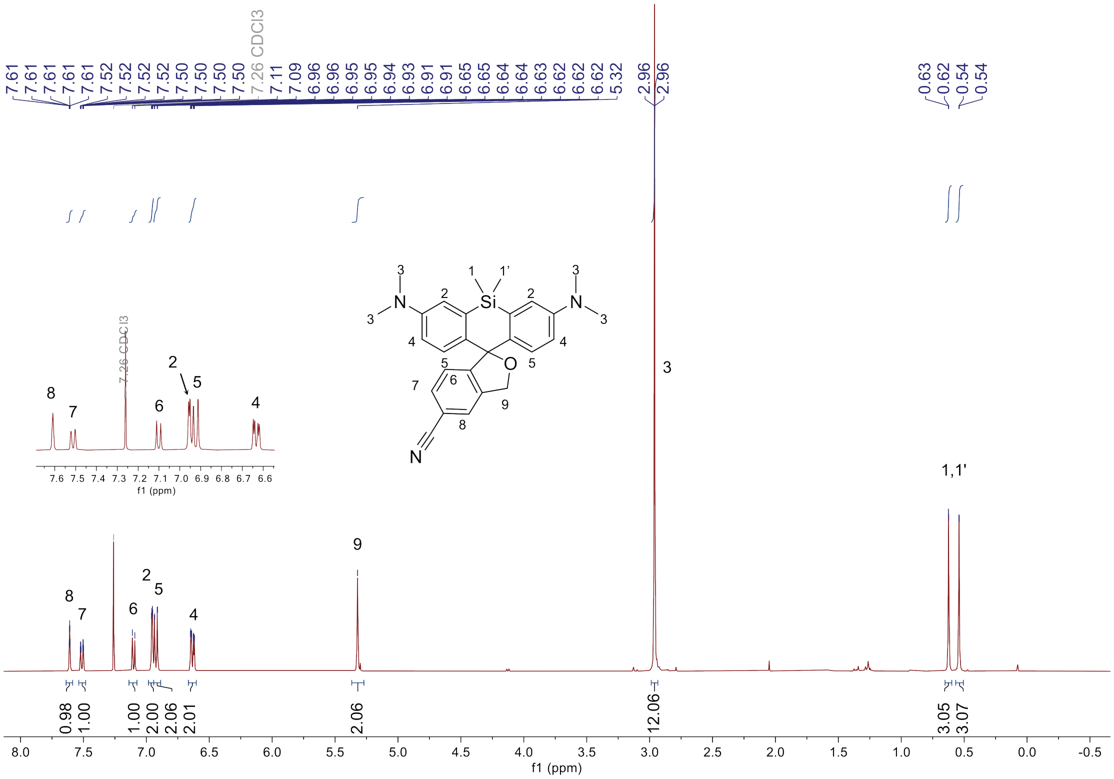

**Figure S16**. $^{1}H$ NMR ($CD_3Cl$, 400 MHz) spectrum of compound **S3.**

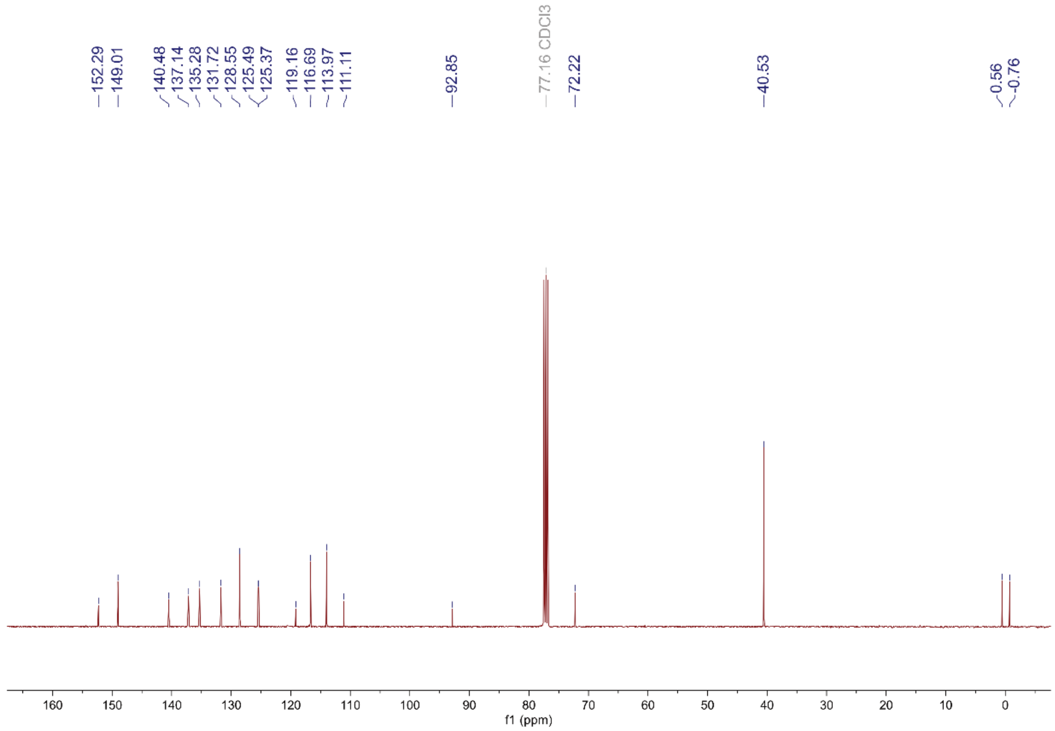

**Figure S2**. $^{13}C$ NMR ($CD_3Cl$, 101 MHz) spectrum of compound **S3.**

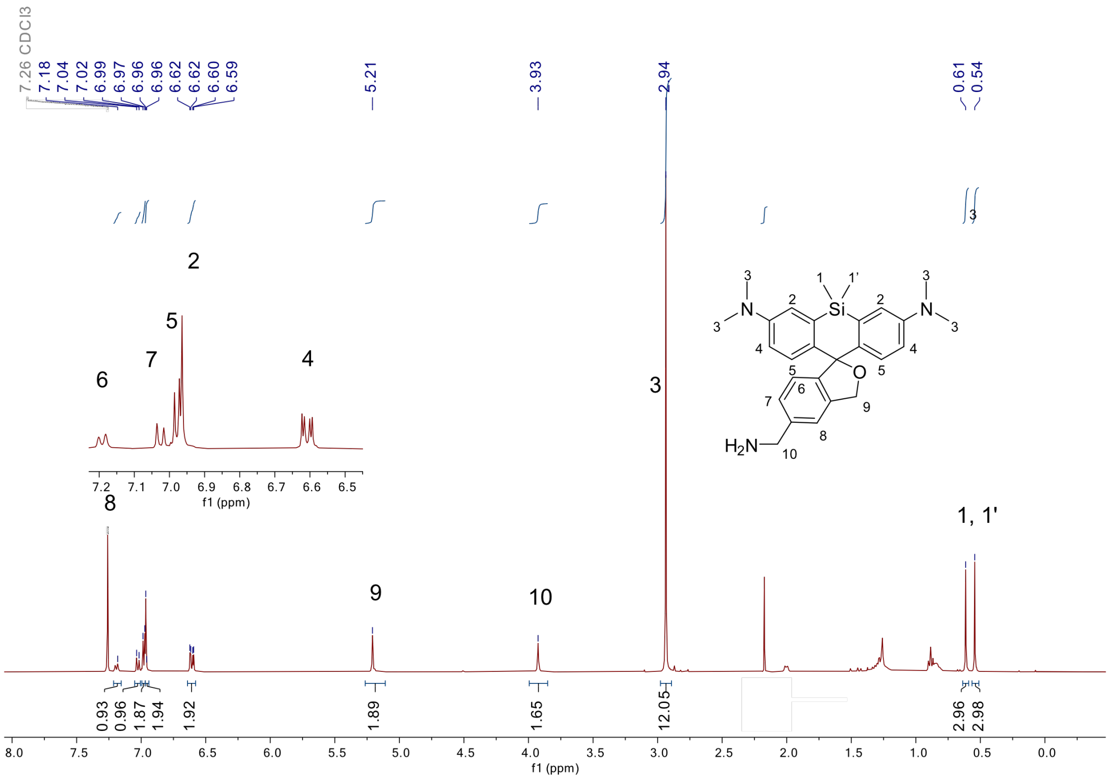

**Figure S3**. $^{1}H$ NMR ($CD_3Cl$, 400 MHz) spectrum of compound **S4.**

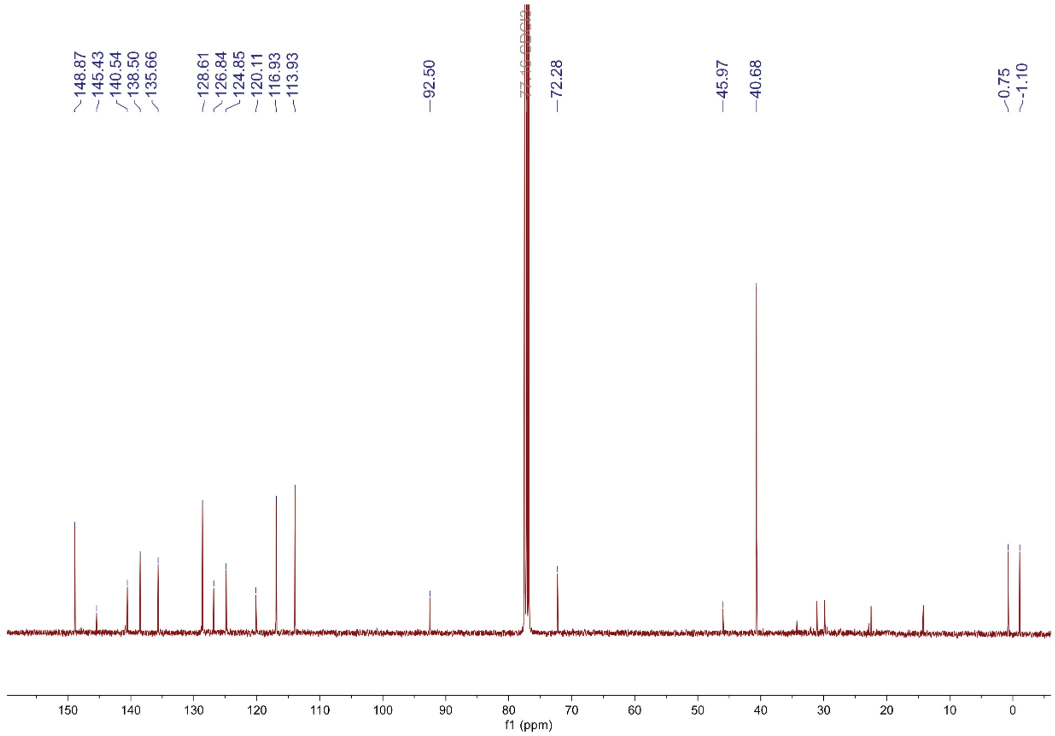

**Figure S39**. $^{13}C$ NMR ($CD_3Cl$, 101 MHz) spectrum of compound **S4.**

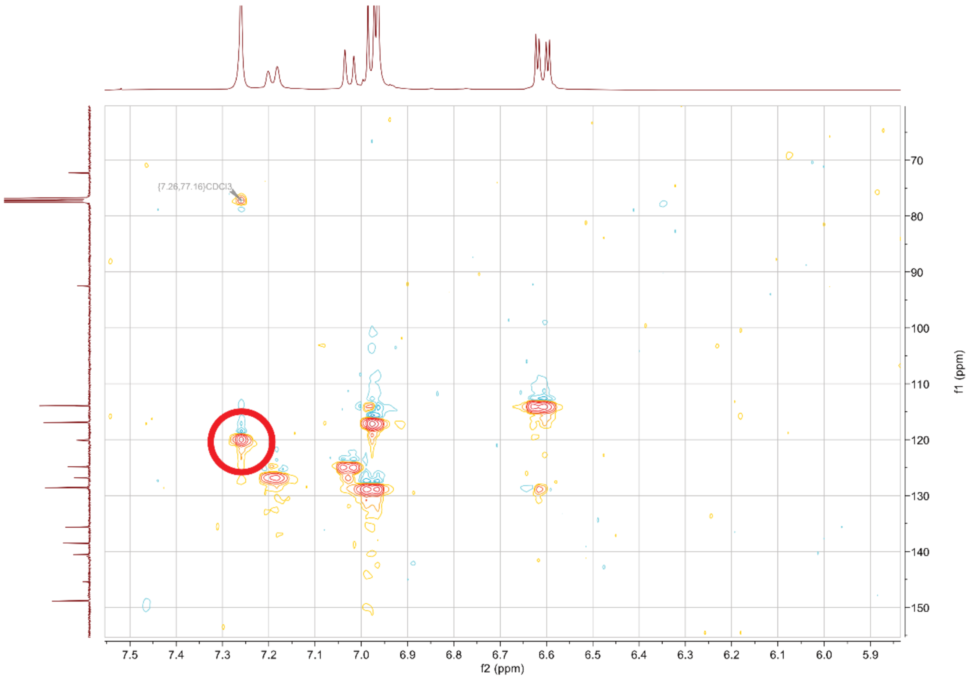

**Figure S40**. HSQC NMR ($CD_3Cl$) Blow-up of region showing the hidden peak of H8 in the HSQC spectrum of compound **S4.**

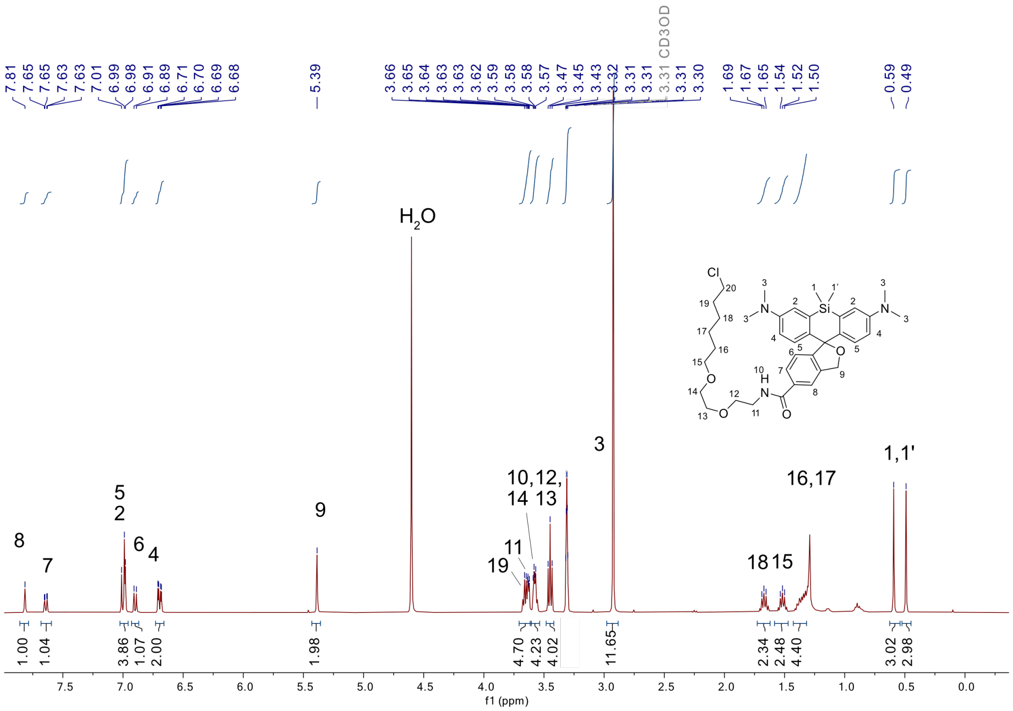

**Figure S41**. $^{1}H$ NMR ($CD_3OD$, 400 MHz) spectrum of compound **HMSiR-Halo** compound **HMSiR-IA.**

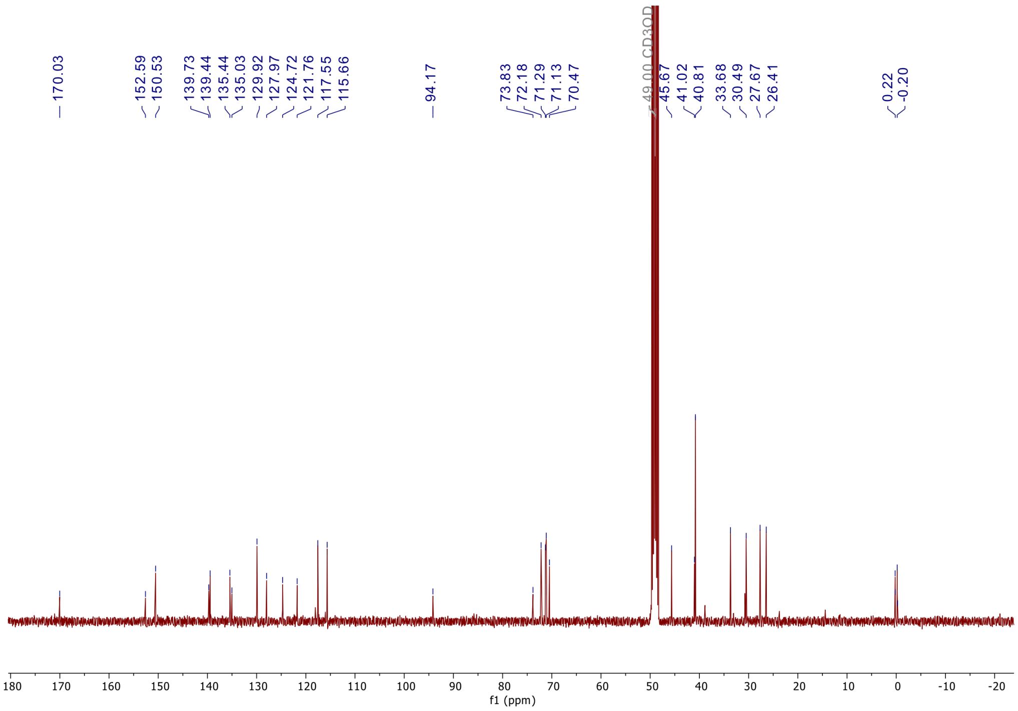

**Figure S42**. $^{13}C$ NMR ($CD_3OD$, 100 MHz) spectrum of compound **HMSiR-Halo** compound **HMSiR-IA.**

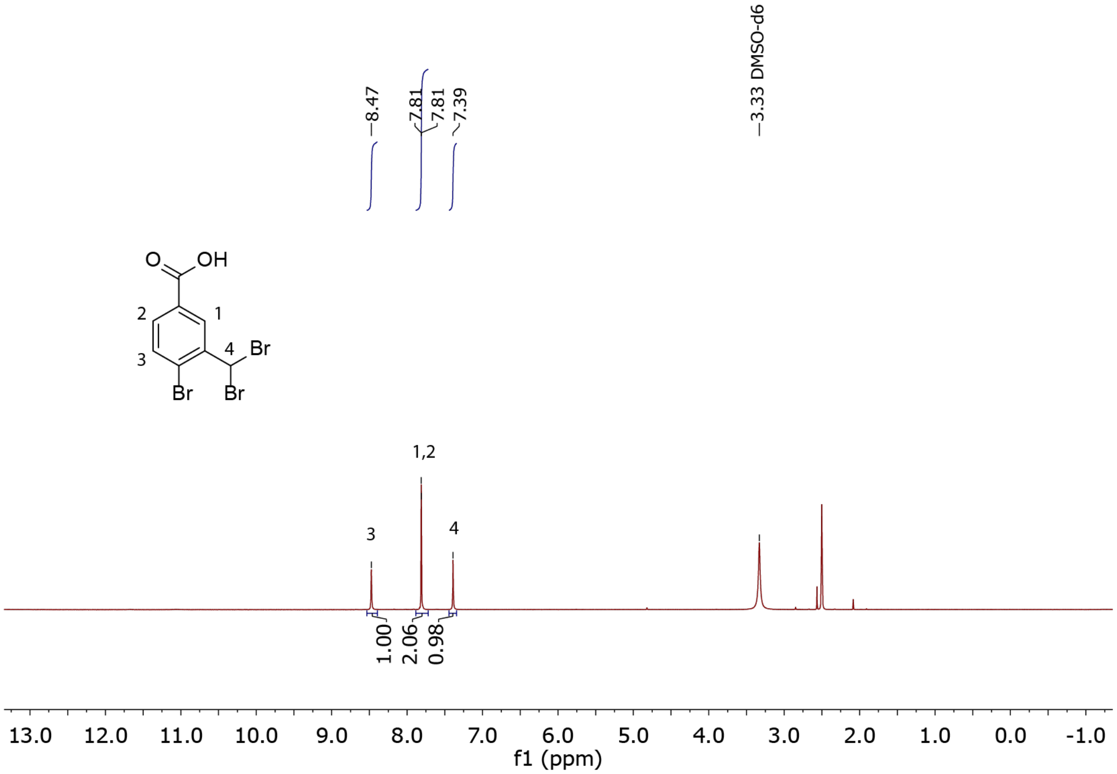

**Figure S43**. [1]H NMR (DMSO-$d_6$, 400 MHz) spectrum of compound **S5.**

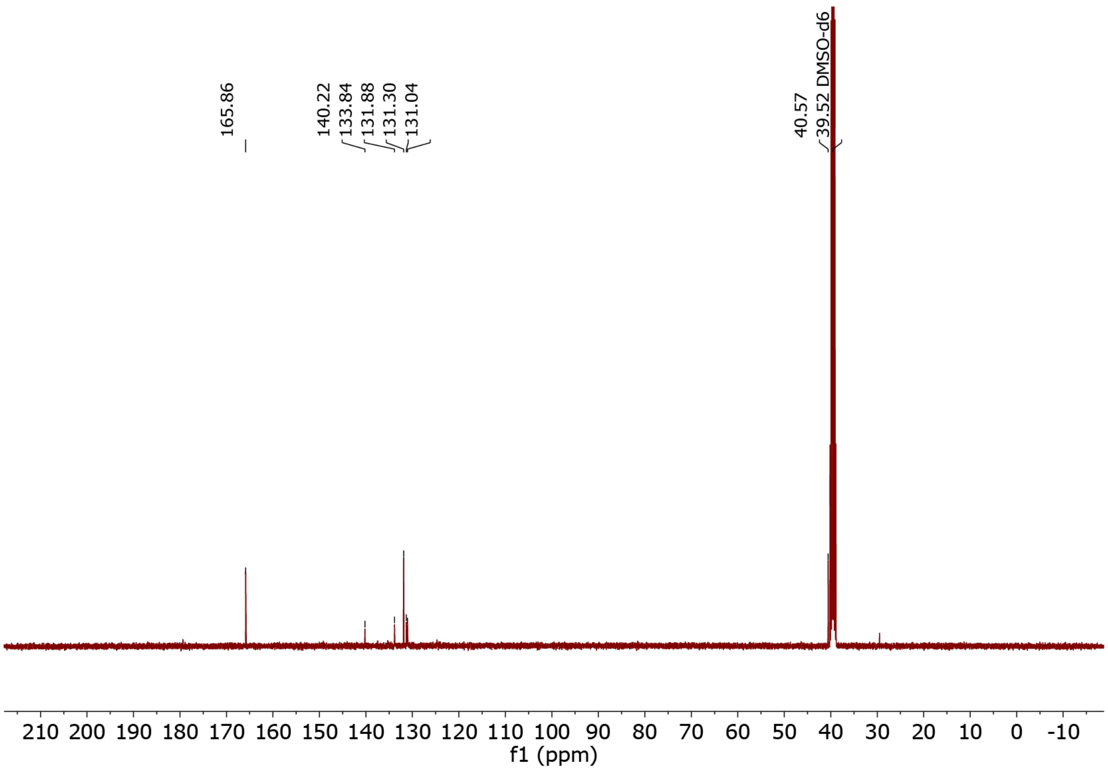

**Figure S44**. [13]C NMR (DMSO-$d_6$, 101 MHz) spectrum of compound **S5.**

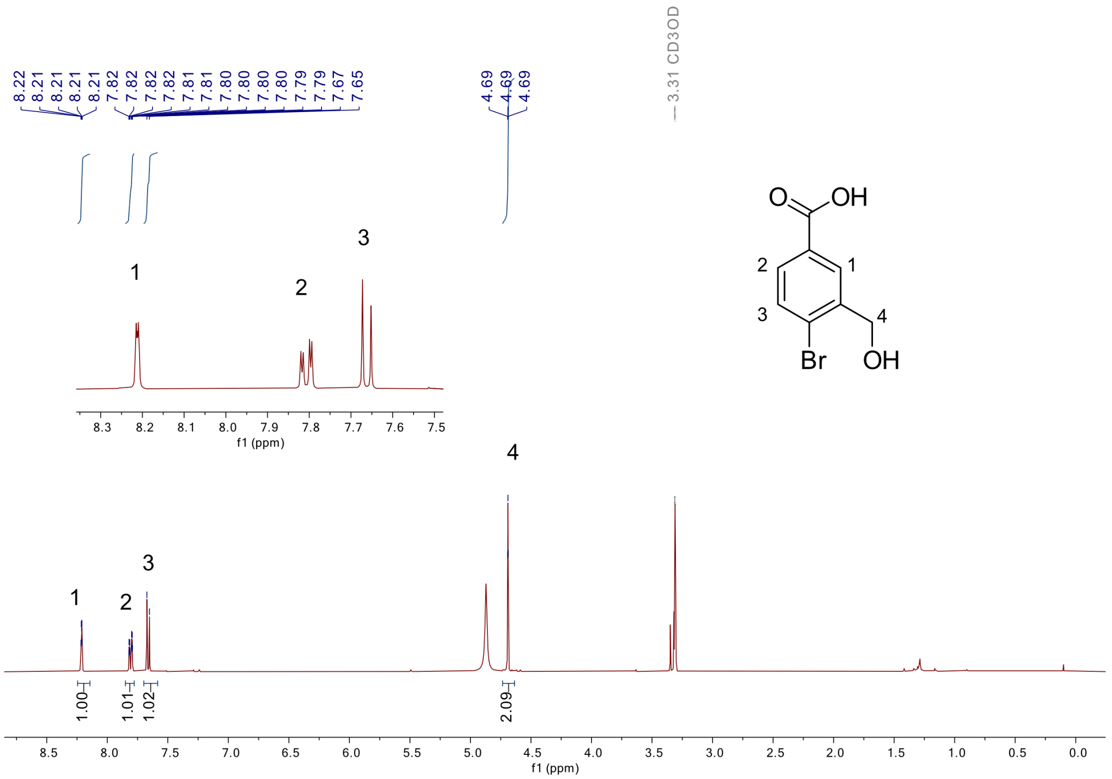

**Figure S45**. [1]H NMR ($CD_3OD$, 400 MHz) spectrum of compound **S6.**

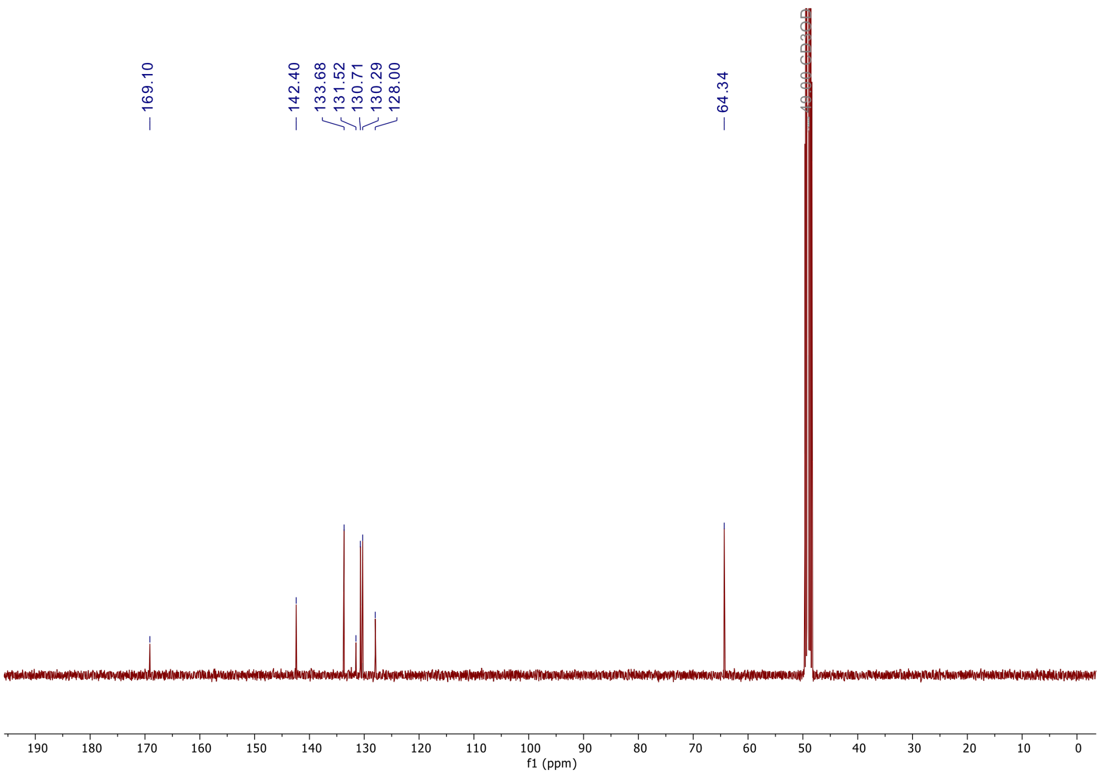

**Figure S46**. [13]C NMR ($CD_3OD$, 101 MHz) spectrum of compound **S6.**

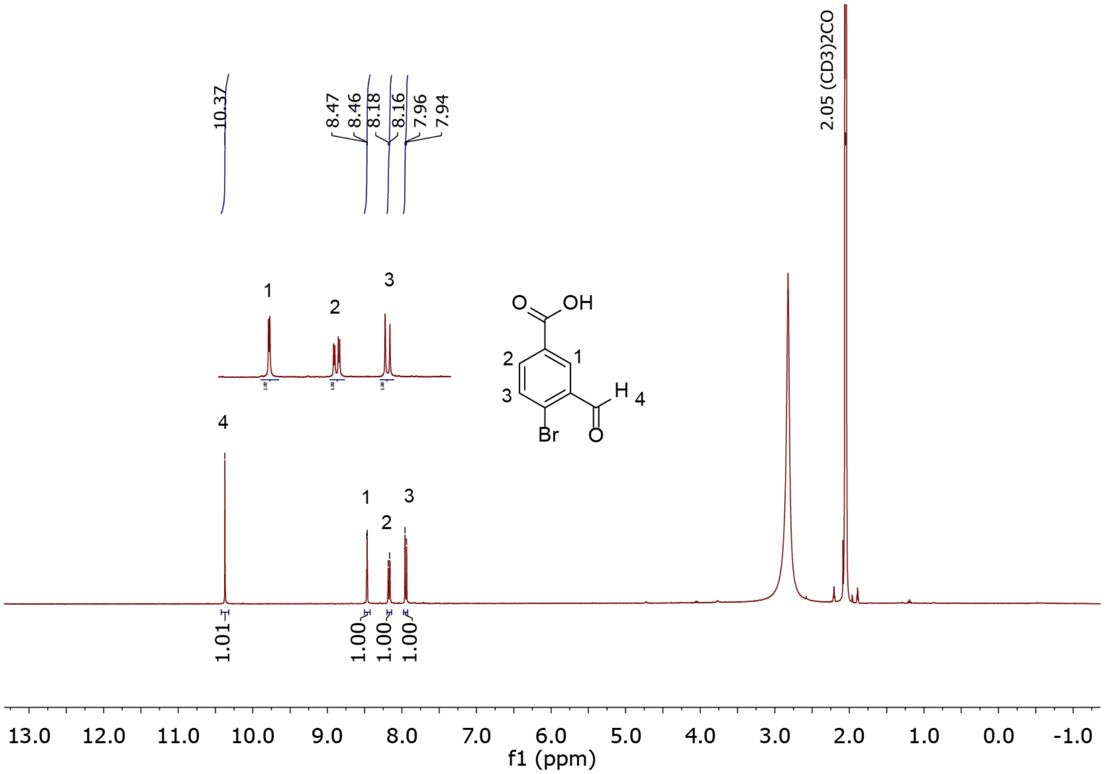

**Figure S47**. $^{1}H$ NMR ($(CD_3)_2CO$, 400 MHz) spectrum of compound **S7.**

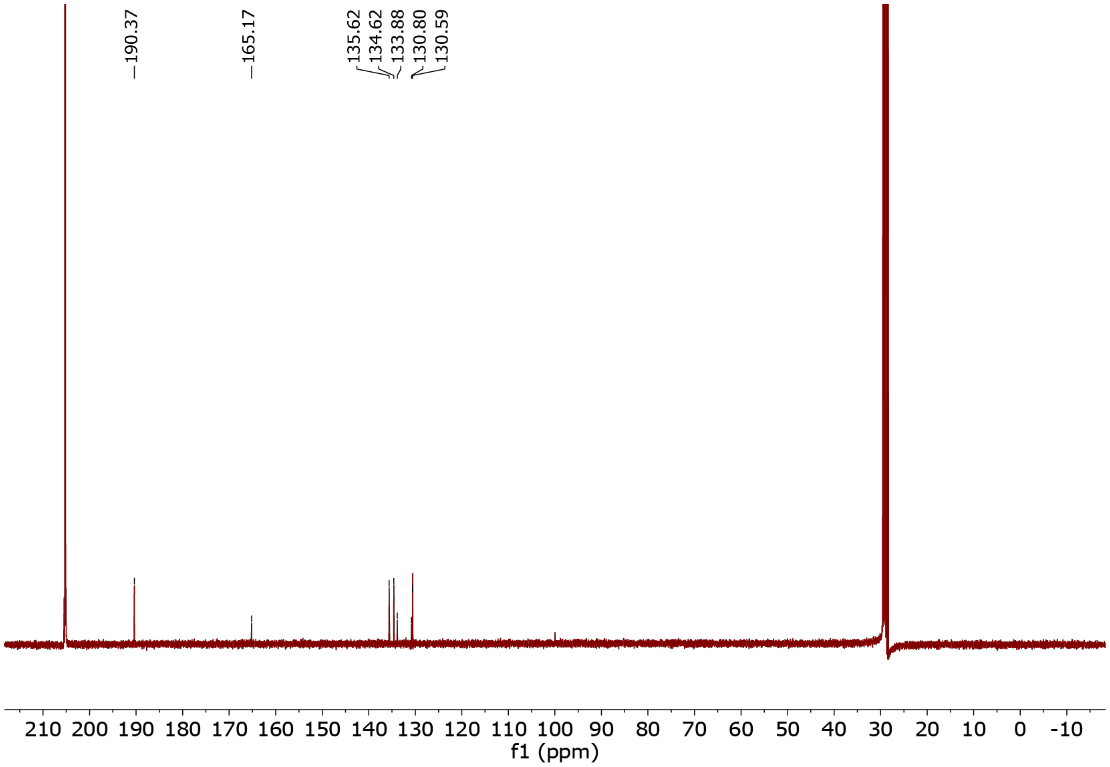

**Figure S48**. $^{13}C$ NMR ($(CD_3)_2CO$, 101 MHz) spectrum of compound **S7.**

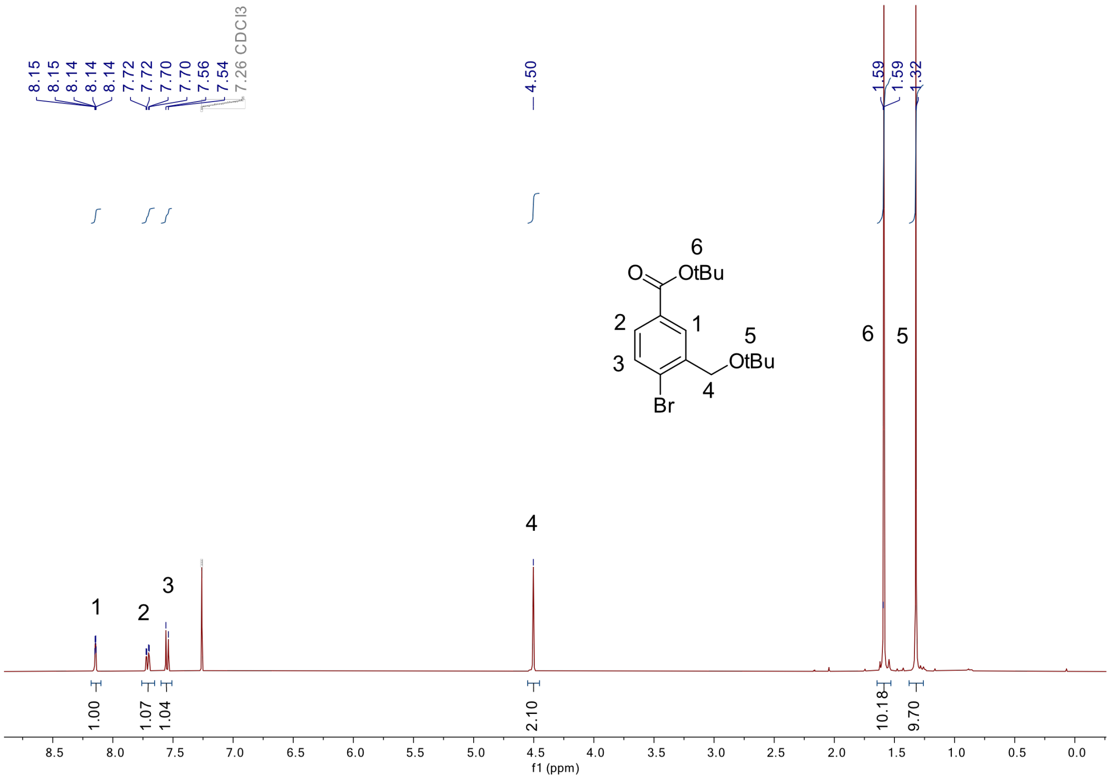

**Figure S49**. $^{1}H$ NMR ($CDCl_3$, 400 MHz) spectrum of compound **S8.**

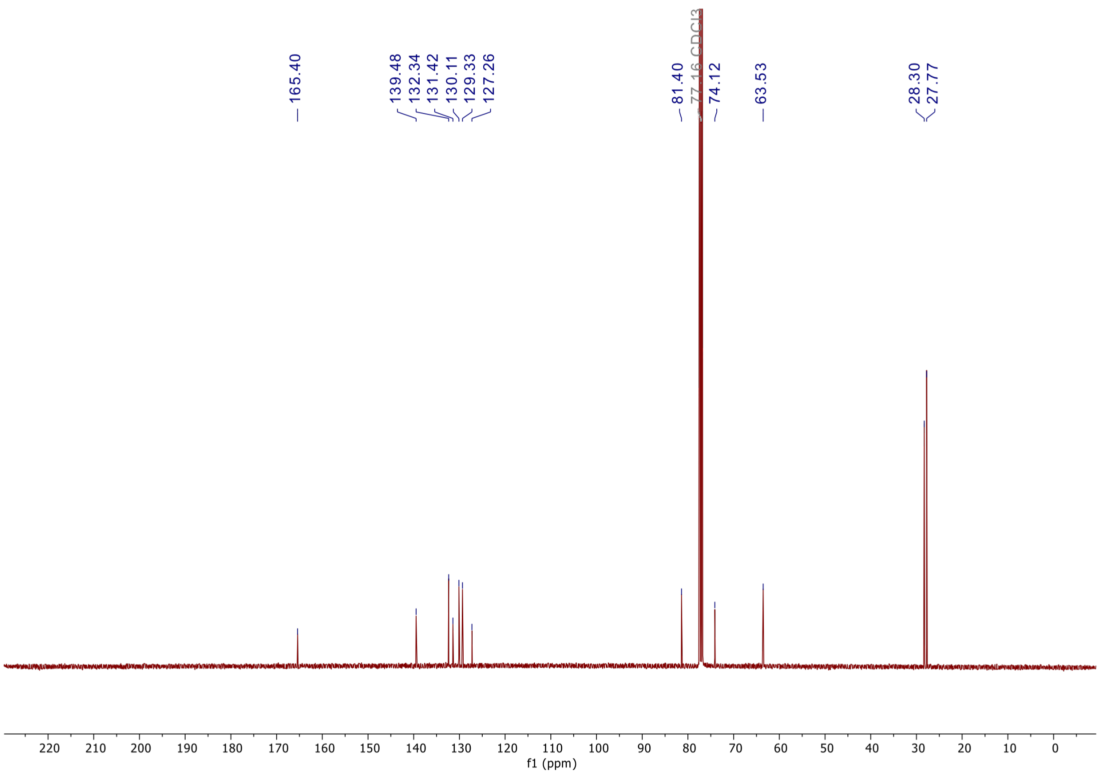

**Figure S50**. $^{13}C$ NMR ($CDCl_3$, 101 MHz) spectrum of compound **S8.**

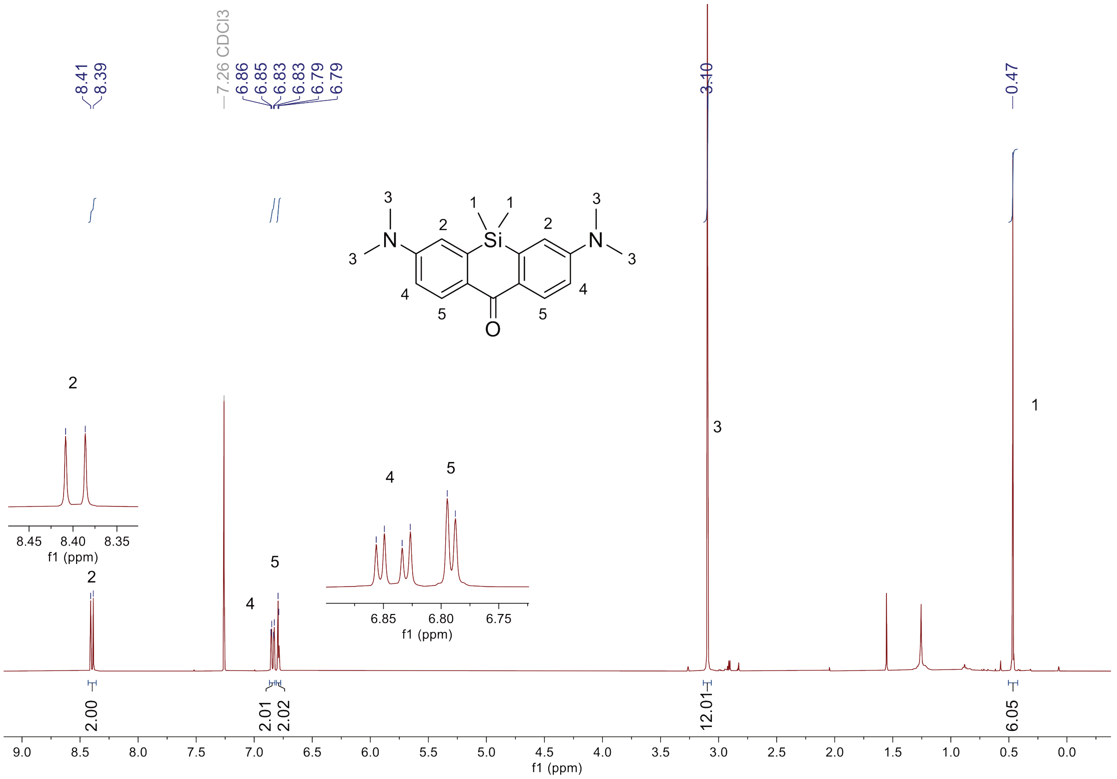

**Figure S51**. $^{1}H$ NMR ($CD_3Cl$, 400 MHz) spectrum of compound **S9.**

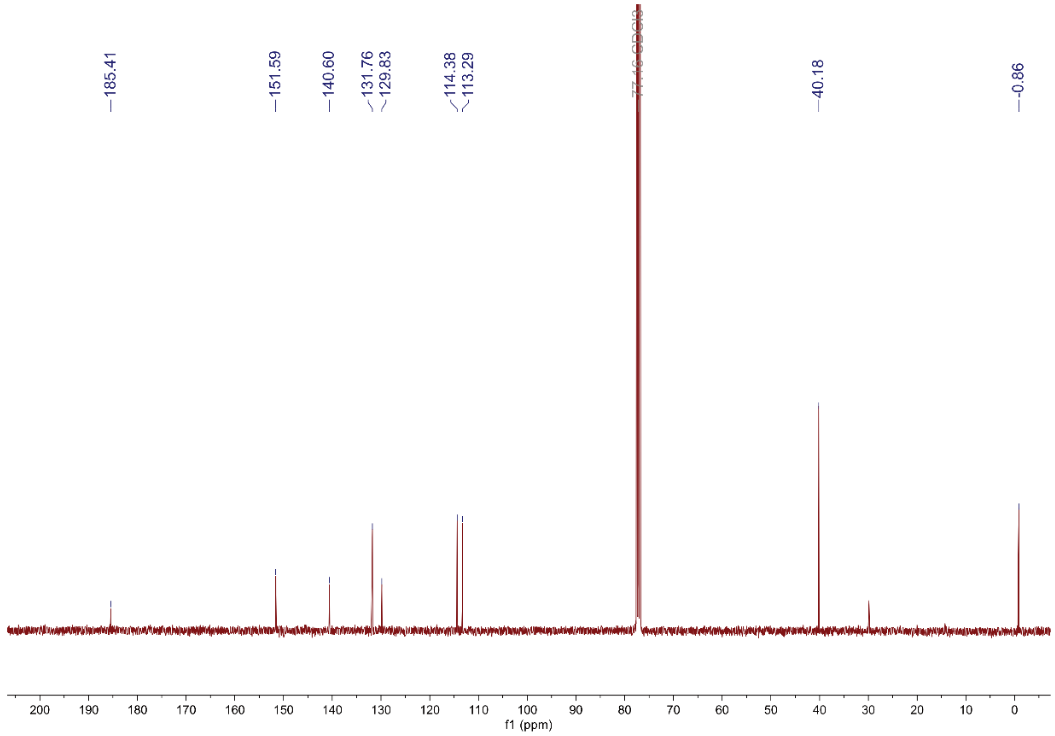

**Figure S52**. $^{13}C$ NMR ($CD_3Cl$, 100 MHz) spectrum of compound **S9.**

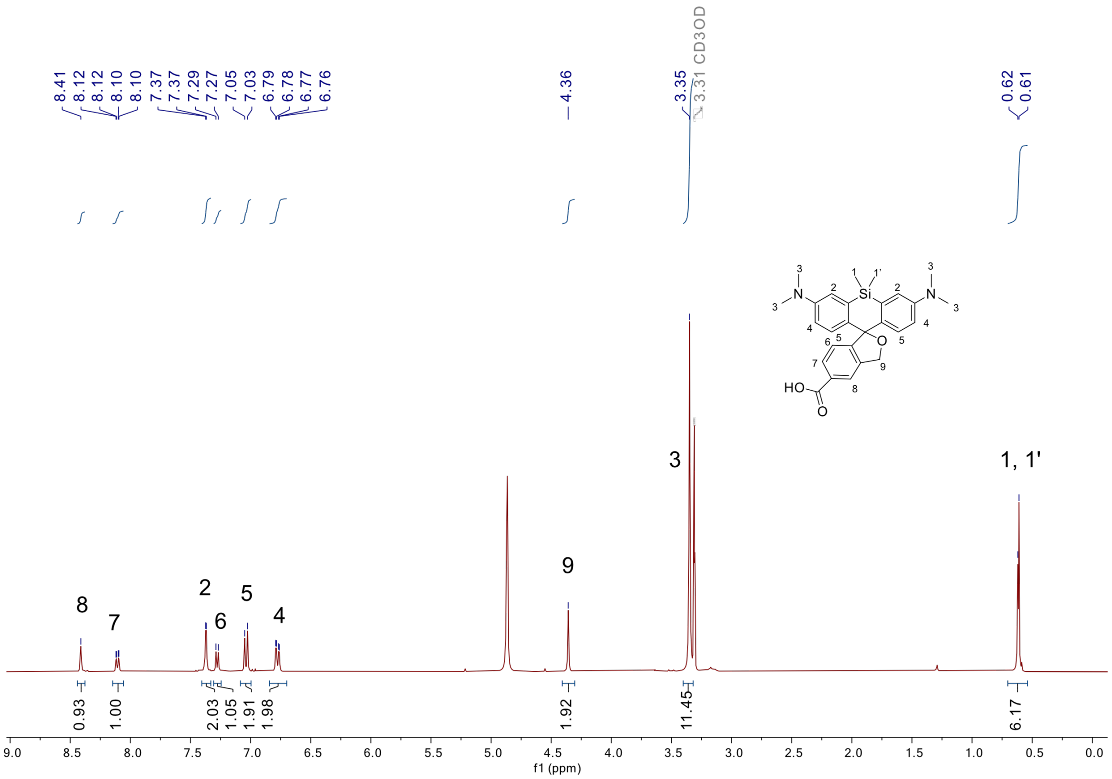

**Figure S53**. $^{1}H$ NMR ($CD_3OD$, 400 MHz) spectrum of compound **S10**.

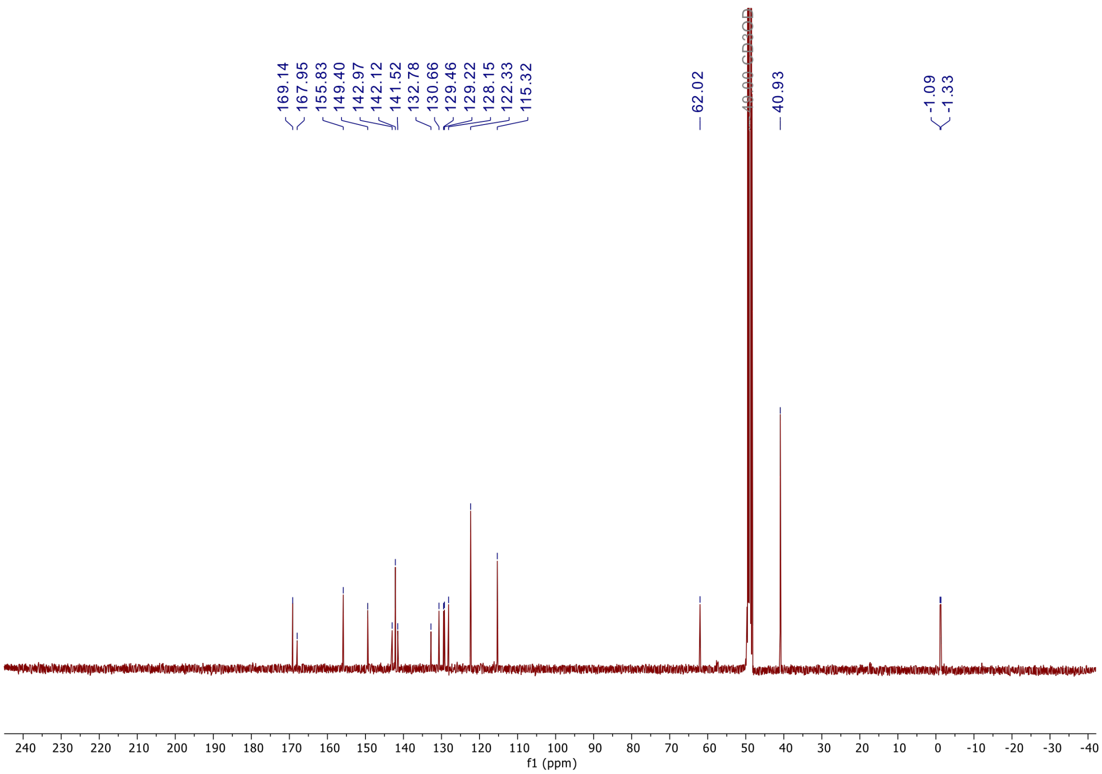

**Figure S54**. $^{13}C$ NMR ($CD_3OD$, 101 MHz) spectrum of compound **S10**.
